# The SKBR3 Cell-Membrane Proteome: Role in Aberrant Cancer Cell Proliferation and Resource for Precision Medicine Applications

**DOI:** 10.1101/2021.10.24.465642

**Authors:** Arba Karcini, Iulia M. Lazar

**Author notes:** **Correspondence:** Iulia M. Lazar, **E-mail:**.

## Abstract

The plasma membrane proteome resides at the interface between the extra- and intra-cellular environment and through its various roles in signal transduction, immune recognition, nutrient transport, and cell-cell and cell-matrix interactions plays an absolutely critical role in determining the fate of a cell. Our work was aimed at exploring the landscape of the cancer cell-membrane proteome responsible for sustaining uncontrolled cell proliferation, and its intrinsic resources for enabling detection and therapeutic interventions. SKBR3/HER2+ breast cancer cells were used as a model system and mass spectrometry for characterizing the proteome. The number of identified cell-membrane proteins exceeded 2,000, with ~1,300 being matched by two or more unique peptides. Classification in four major categories, i.e., proteins with receptor or enzymatic activity, CD antigens, transporters, and cell adhesion proteins, uncovered overlapping roles in biological processes that drive cell growth, apoptosis, differentiation, immune response, adhesion and migration, as well as capabilities for signaling crosstalk and alternate pathways for proliferation. The large number of tumor markers (>50) and putative drug targets (>100) exposed a vast potential for yet unexplored detection and targeting opportunities, whereas the presence of 15 antigen immunological markers enabled an assessment of epithelial, mesenchymal or stemness characteristics. Serum-starved cells displayed altered processes related to mitochondrial OXPHOS/ATP synthesis, protein folding and localization, while serum-treated cells exhibited attributes that support tissue invasion and metastasis. Altogether, our findings advance the understanding of the biological triggers that sustain aberrant cancer cell proliferation, survival and development of resistance to therapeutic drugs, and reveal the vast innate opportunities for guiding immunological profiling and precision medicine applications aimed at target selection or drug discovery.

## Introduction

Breast cancer is a common form of cancer that continues to lead, even in the present day, to a large number of deaths among women worldwide (1). The different breast cancer subtypes are defined based on the presence of ER, HER2 or PR receptors, whether alone or in combination. HER2+ and triple negative breast cancers have the worst prognosis due to the fact that some HER2+ tumors are either non-responsive or develop resistance to anti-HER2 therapies, while triple negative cancers are non-responsive to hormonal therapies or drugs that target HER2 receptors (1,2). As a result, focus has been placed on the development of novel therapeutic approaches that rely either on the use of various drug cocktails and treatment regimens that target multiple receptors or compensatory and downstream crosstalk signaling pathways of HER2, or, more recently, on triggering immune system responses that attack the cancer cells (2).

The heavy interest in the study of cancer cell-membrane receptors has been fueled by their central role in initiating cellular signaling cascades that lead to aberrant cell proliferation, as well as by their potential as cancer markers or drug targets. Cell-membrane receptors include three traditional protein categories, i.e., G-protein-coupled receptors (GPCRs), ion channels, and enzyme-linked receptors, mostly represented by receptor tyrosine kinases (RTKs) (3). GPCRs represent the largest class of receptors (4), while the enzyme-linked receptors the most studied one (5), and together they comprise the majority of drug targets. The aberrant activity of these receptors was linked to many diseases including inflammation, metabolic disorders, and cancer (6). Targeting, for example, HER2 receptors has been at the core of targeting HER2+ tumors. Proteomic analysis of cell-surface (CS) proteins has revealed, however, many important, additional roles for other CS proteins in cancer proliferation (7). The detection and characterization of these cell-surface targets has been, nevertheless, challenging due to compounding factors such as low abundance, hydrophobicity, presence of post-translational modifications (PTMs), and heterogeneity (8,9).

Several methods have been developed for the isolation of cell-surface proteins relying mainly on ultra-centrifugation, coating of the plasma membrane with silica-beads, and chemical labeling of N-linked glycosylated proteins or of protein amine, sulfhydryls or aldehyde groups, followed by affinity pulldown (8–12). After isolation, the state-of-the art for detecting the enriched CS protein fractions involves mass spectrometry (MS) analysis. The advanced capabilities of the MS technology (i.e., high sensitivity, high mass accuracy, and quantification capability) enabled the detection of thousands of proteins per cell line, the compilation of comprehensive cell-surface protein data into interactive databases such as The Cell-Surface Protein Atlas (13) ~1500 human proteins, and the development of even more comprehensive lists constructed with machine learning based predictor tools (~2900 human proteins, 14). Altogether, these studies have contributed to the overall knowledge of what has been named the “surfaceome,” and its associated signaling networks in humans (14,15), that altogether was captured in comprehensive public repositories (16–19).

To capitalize on the wealth of information that can be generated through mass spectrometric analysis, this study was aimed at characterizing the cell-surface proteome of SKBR3/HER2+ breast cancer cells by using orthogonal methods for cell-surface protein enrichment and isolation, categorizing these proteins based on their functional role and relevance to cancer, identifying key drivers of aberrant proliferation, and exploring the opportunities presented by such cells for the development of effective diagnostic and therapeutic approaches. We also report on the remodeling of the cell-membrane proteome under serum-starved and serum-supplemented conditions, and, lastly, we draw insights into the signaling cascades initiated at the plasma membrane and the potential crosstalk activities that fuel the development of resistance to treatment with therapeutic drugs.

## Methods

### Reagents and materials

SKBR3 cells, trypsin (0.25 %)/EDTA (0.53 mM) and PenStrep solution were purchased from ATCC (Manassas, VA), and fetal bovine serum (FBS) from Gemini Bio-Products (West Sacramento, CA). McCoy’s 5A (Modified) medium, Dulbecco’s Phosphate Buffered Saline solution (DPBS), DPBS with calcium and magnesium (+Ca^2+^/Mg^2+^), and TrypLE Select Enzyme solutions were purchased from Gibco (Carlsbad, CA). Sample processing reagents such as NaF, Na_3_VO_4_, dithiothreitol (DTT), urea, ammonium bicarbonate (NH_4_HCO_3_), acetic acid, and trifluoroacetic acid (TFA) were from Sigma (St. Louis, MO). Sequencing grade trypsin and trypsin/LysC were purchased from Promega (Madison, WI). Protease inhibitors cocktail (HALT), EZ-Link Sulfo-NHS-SS-Biotin Pierce Cell-surface Biotinylation and Isolation Kit, EZ-Link Alkoxyamine-PEG4-Biotin, Pierce Sodium meta-Periodate, EasyPep Mini MS Sample Prep Kit, and Streptavidin Alexa Fluor 488 conjugate were purchased from Thermo Scientific (Rockford, IL). Aniline was from BeanTown Chemical Corporation (Hudson, NH). SPEC-PTC18, SPEC-PTSCX sample cleanup pipette tips and Bond Elut C18/ 3 mL cleanup cartridges were from Agilent (Santa Clara, CA), cell culture slides (8-chamber) for fluorescent visualization of cells from MatTek (Ashland, MA), and Nunc cell culture flasks from Thermo Scientific. HPLC-grade solvents such as methanol and acetonitrile were purchased from Fisher Scientific (Fair Lawn, NJ). Water for the preparation of sample solutions and LC eluents was either produced by a MilliQ Ultrapure water system (Millipore, Bedford, MA) or was distilled from DI water.

### Cell culture

The SKBR3 cells were cultured in McCoy’s 5A medium and FBS (10 %) in T175 Nunc flasks, at 37 °C and in the presence of CO_2_ (5 %). After reaching ~70-80 % confluence, for the first set of culture conditions, the cells were washed twice with serum-free medium and incubated in McCoy 5A for 48 h without any supplements. For the second set of culture conditions, after 48 h serum starvation, the cells were incubated for 24 h in McCoy 5A supplemented with FBS (10 %). Penstrep (0.5 %) was added to all culture media to prevent bacterial contamination. Two T175 flasks of serum-free or serum-treated cells (~90 % confluence) were prepared for each cell-surface protein harvesting procedure, by either chemical labelling or proteolytic cleavage methods, as described below. Three distinct biological replicates (n=3) of each condition were generated for analysis.

### Cell-membrane protein labeling and harvesting

SKBR3 cell-membrane protein enrichment was performed by using three complementary procedures, two including biotin labeling and one relying on the trypsinization of cell-membrane extracellular protein domains. All reagent solutions were prepared fresh before use, and the reagent and rinse solutions that were used for biotin labeling were cooled to 4 °C before adding to the cells. The first labeling procedure involved the use of EZ-Link Sulfo-NHS-SS-Biotin (0.5 mg/mL) for labeling the protein N-terminal and Lys side-chain amino groups. Cells were rinsed twice with DPBS (+Ca^2+^/Mg^2+^) and then incubated at 4 °C for 30 min, in the dark, with the biotin reagent. After incubation, the biotin reagent was removed, and each flask was washed twice with 20 mL Tris quenching buffer solution (0.1 M) provided in the kit. The cells were collected by scraping in Tris-buffer (10 mL per flask), and centrifuged for 5 min at 800 x g and 4 °C. The second approach involved the labeling of cell-surface glycoproteins with EZ-Link Alkoxyamine-PEG4-Biotin (0.5 mg/mL) following protocols described by the manufacturer and in previous manuscripts (12), with some modifications. Briefly, the cells were rinsed twice with DPBS (+Ca^2+^/Mg^2+^) and incubated at 4 °C for 30 min, in the dark, with 20 mL sodium meta-periodate solution (1 mM, pH 6.5) to oxidize the glycan moieties of cell-surface proteins. The cells were rinsed again, twice, with DPBS (+Ca^2+^/Mg^2+^), and incubated with 12 mL biotin reagent solution in the presence of 10 mM aniline at 4 °C for 30 min, in the dark. After the completion of the labeling reaction, the biotin reagent was removed, and each flask was washed twice with 20 mL DPBS (+Ca^2+^/Mg^2+^). Cell-surface protein biotinylation of cells was visualized with an inverted Eclipse TE2000-U epi-fluorescence microscope (Nikon, Melville, NY), after staining the cells with Streptavidin Alexa Fluor™ 488 (4 ug/mL). The cells were collected by scraping in 10 mL DPBS and centrifuged for 5 min at 800 x g and 4 °C. The labeled cell pellets generated by either procedure were frozen at −80 °C for further processing or subjected to immediate lysis. The third approach consisted of shaving the cell-surface protein ectodomains with TrypLE, a reagent that contains recombinant enzymes for cell dissociation that are free of animal origin trypsin. For this procedure, the SKBR3 cells were washed twice with serum-free medium, and incubated with 10 mL TrypLE solution at 37 °C, with 5 % CO_2_, for 2-4 min. The incubation time was short, to prevent cell detachment. The cell supernatant containing the cell-surface protein ectodomains was then collected, centrifuged for 5 min at 500 x g and 4 °C for the removal of floating cells, and frozen at −80 °C. The samples generated through the three enrichment procedures will be referred from now on as the amine, glyco, and trypsin samples.

### Cell-membrane protein recovery and processing

To isolate the cell-surface proteins of the amine-biotinylated samples, the cells were lysed with 500 uL Lysis Buffer (Pierce) supplemented with HALT protease inhibitor cocktail (5 uL), for 30 min, on ice, with intermittent vortexing and sonication. The lysate (~500 uL) was collected by centrifugation (15,000 x g, 5 min, 4 °C) and incubated with 250 uL NeutrAvidin beads at room temperature for 2 h, followed by 4 washes with Wash Buffer (Pierce) and 3 washes with NH_4_HCO_3_ (100 mM). After each wash, the beads were isolated by centrifugation (1,000 x g, 1 min). Protein recovery from the beads was performed by proteolytic digestion, on the bead, overnight, at room temperature, in 200 uL NH_4_HCO_3_ (100 mM) supplemented with 25 uL trypsin/Lys C solution (10-12 ug enzyme). After centrifugation (1,000 x g), the beads were further treated with 200 uL DTT (10 mM) for 1 h at room temperature to recover the di-thiol, covalently bound remaining protein fragments. Both on-bead protein digest and DTT-released fractions were collected and denatured with urea (8 M) for 1 h at 57 °C (the on-bead digest solution was also added DTT, 5 mM). After dilution with NH_4_HCO_3_ (100 mM) to reduce the urea concentration to <1 M, the samples were subjected to a second digestion in solution with 10 uL trypsin (~5 ug enzyme) for 4 h at 37 °C. After quenching the enzymatic reaction with TFA, the cell-surface peptide extracts were processed for salt and detergent disposal with SPEC-PTC18 and SPEC-SCX cartridges. Protein concentration measurements could not be performed due to limited sample availability and the low abundance of the cell-surface proteins in solution. Isolation of the cell-surface proteins of the biotinylated glyco samples was performed by following a similar procedure to the one that was used for the amine-labeled samples. The cell lysate was incubated with NeutrAvidin beads, the beads were treated with 200 uL DTT (45 mM, 1 h, room temperature, dark), and after the removal of the DTT solution by centrifugation (1000 x g, 1 min), on-bead proteolytic digestion was performed overnight, at room temperature, in 200 uL solution of NH_4_HCO_3_ (100 mM) with 25 uL trypsin/Lys C (10-12 ug enzyme) in the presence of urea (1 M). An additional 4 h digestion at room temperature was performed by adding to the beads 100 uL NH_4_HCO_3_ (50 mM) and 10 uL trypsin solution (~5 ug enzyme). The collected glycoprotein fraction was then processed with SPEC-PTC18 and SPEC-SCX cartridges. To isolate the cell-surface proteins of the trypsinized samples, the collected solution (~10 mL) was digested in a preliminary stage, overnight, with 20 ug trypsin at 37 °C, and concentrated then on a Bond Elut C18 column to remove the large volume of TrypLE solution. The sample was then reconstituted in Tris-buffer (120 uL, 50 mM, pH=8) and denatured with urea (8 M) and DTT (5 mM) for 1 h at 57 °C. After reducing the urea concentration to <1 M with NH_4_HCO_3_ (100 mM), the sample was subjected to a second digestion with trypsin (~5 ug) for 4 h at 37 °C. After cleanup, all peptide samples were dissolved in 30 uL CH_3_CN/H_2_O/TFA (95-98):(2-5):0.01 v/v for LC-MS analysis.

### LC-MS analysis

The peptide samples were analyzed with an EASY-nLC 1200 UHPLC system (ThermoFisher Scientific) by using a heated nano-electrospray ionization (ESI) source (2 kV) and a Q Exactive hybrid quadrupole-Orbitrap mass spectrometer (ThermoFisher Scientific). An EASY-Spray column ES802A (150 mm long, 75 μm i.d., 3 μm C18/silica particles, ThermoFisher Scientific) was used at flow rates of 250 nL/min. The mobile phases were prepared from H_2_O:CH_3_CN:TFA, and mixed in ratios of 96:4:0.01 v/v for mobile phase A and 10:90:0.01 v/v for B. During a separation gradient of 85 mins, the eluent B concentration was increased from 3 % to 30 % (5-65 min), 45 % (65-72 min), 60 % (72-73 min), and 90 % (73-74 min), where it was kept for 5 min, and then decreased to a final concentration of 3 %. The MS data were acquired over a range of 400-1600 m/z with resolution set to 70,000, AGC target to 3E6, and maximum IT to 100 ms. Data-dependent MS2 acquisition (dd-MS2) was enabled by using higher-energy collisional dissociation (HCD), isolating the precursor ions with a width of 2.4 m/z, and fragmenting them at 30 % normalized collision energy (NCE). dd-MS2 acquisition parameters were set to resolution 17,500, AGC target 1E5 (minimum AGC target 2E3 and intensity threshold 4E4), maximum IT 50 ms, and loop count 20. Charge exclusion was enabled for unassigned and +1 charges, apex trigger was set to 1 to 2 s, dynamic exclusion lasted for 10 s for chromatographic peak widths of 8 s, and the features of isotope exclusion and preferred peptide match were turned on. For parallel reaction monitoring (PRM) validation, the peptides of interest were searched within a time-window of +/-10 min of the precursor ion retention time following the same separation gradient as described above. The precursor ions were isolated with a width of 2.0 m/z, and fragmented at 30 % normalized collision energy with PRM parameters set as follows: resolution 35,000, AGC target 2E5, and maximum IT 110 ms.

### MS raw data processing

The MS data were processed by the Proteome Discoverer 2.4 package (Thermo Fisher Scientific, Waltham, MA) and searched with Sequest HT against a *Homo sapiens* database (DB) of 24,433 reviewed, non-redundant UniProt protein sequences (March 2019 download). The processing workflow spectrum filter was set for a peptide precursor mass range of 400-5,000 Da, and the Sequest HT node parameters allowed for the selection of fully tryptic peptides (6-144 aa length) with maximum two missed cleavages, 15 ppm precursor ion tolerance, b/y/a ion fragments with 0.02 Da tolerance, and dynamic modifications (maximum 4 per peptide) on Met (15.995 Da/oxidation) and the protein N-terminal amino acids (42.011 Da/acetyl). Carbohydrate group labeling and trypsinization do not alter the chemical structure of the cell-surface proteins, but labeling of amine groups with the biotinylation reagent forms a 3-mercapto-propanamide derivative, for which a dynamic modification of 87.998 on the Lys residues was also enabled. The raw files were processed independently for the fraction of proteins generated by direct on-bead digestion and DTT reduction, but for reporting, the results were merged. Overwhelmingly, though, the majority of protein identifications were enabled by the on-bead digestion step, rendering the additional DTT recovery step unnecessary. The peptide spectrum match (PSM) validator node used a target/decoy concatenated database strategy to calculate the FDR targets of 0.01 (strict) and 0.03 (relaxed) based on search engine Xcorr scores (input data of maximum DeltaCn 0.05 and maximum rank 1). Additional parameters were set in the consensus workflow for both peptide and protein levels. The peptide group modification site probability threshold was set to 75. Peptide confidences were represented by the corresponding best PSM confidences. Only peptides of at least medium confidence and proteins matched by only rank 1 peptides were retained in the peptide/protein filter node. The peptides were counted only for top scoring proteins. The protein FDR validator node used the protein scores from the target and decoy searches to calculate the FDRs and rank the proteins, and then calculate the q-values from the FDRs at each score threshold. The FDRs were set to 0.01 (high) and 0.03 (medium) for PSMs, peptides, and proteins, and the strict parsimony principle was enabled for protein grouping. The PRM data were processed by Skyline 20.2 (20) by using a mass spectral library generated from cell-surface protein samples produced by the glycoprotein enrichment method. The *b* and *y* ions were selected from “ion 2” to “last ion” with precursor charges of 2 and 3, and fragment charges of 1 and 2. The library ion match tolerance was 0.02 m/z, and the 5 or 10 most intense product ions were picked from the filtered product ions. The presence of a peptide was considered validated when the peptide displayed a minimum of 5 transitions and a dot product (*dotp*) score >0.9.

### Bioinformatics data interpretation and visualization

An in-house database of cell-membrane proteins was built by extracting relevant entries from the UniProt database based on controlled vocabulary terms (16), from the Human Protein Atlas (HPA) Cellular and Organelle Proteome (17,18), and from the scientific literature (12–15). GeneCards (21) and UniProt (16) were used to assess protein functionality. STRING 11.5 was used to build protein-protein interactions (PPI) networks and assess GO enrichment in biological processes (22), with interaction score confidences set to medium/high and enrichment FDR<0.05. Cytoscape 3.8.2 was utilized to visualize protein networks (23) based on interactomics data exported from STRING, RAWGraphsan open source data visualization framework-for building the dendograms (24), InteractiVenn.net for building Venn diagrams, and Protter for visualizing the location of a protein relative to the cell-membrane bilayer (25).

### Statistical analysis of changes in protein abundance

For each of the three biological replicates (n=3), three LC-MS/MS technical replicates were performed, and the results of the three technical replicates were combined in one multiconsensus protein and peptide report. Protein detection reproducibility and quantitation was performed based on spectral counting. The strength of the bivariate (linear) relationship between any two sets of biological replicates was evaluated with the Pearson correlation coefficient “r”. For evaluating changes in protein abundance, missing values were handled by adding one spectral count to each protein from the dataset. Data normalization was performed based on spectral counting, in two steps. In the first step, normalization was performed at the global level by averaging the total spectral counts of the six samples taken into consideration (i.e., three SF and three ST biological replicates), and using the resulting average as a correction factor (CF1) for adjusting the counts of individual proteins in each sample. In the second step, normalization was performed at the cell-surface protein level by calculating a second correction factor (CF2) based on the spectral counts of only a short list of 10 endogenous cell-surface proteins that were already corrected by CF1 (see equations below). Proteins that changed abundance in the cell-surface proteome were selected by calculating the Log2 values of the ST/SF spectral count ratios and using a two-tailed t-test for assessing significance. Proteins matched by two unique peptides with fold change (FC)≥2 in normalized spectral counts and p-value<0.05 were considered for discussion.

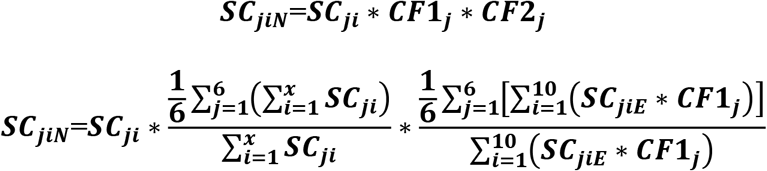

SC_ji_ = spectral count of protein “i” in data set “j” (j=1-6); SC_jiN_ = normalized SC_ji_; SC_jiE_ = spectral count of endogenous membrane protein “i” in data set “j” (10 endogenous proteins were considered); x=total number of proteins identified in the 6 sample sets taken for comparison (3 x ST vs. 3 x ST).

## Results and Discussion

### Cell-membrane protein database

To create a theoretical framework for mapping the cell-membrane proteome, an in-house database containing 7,760 proteins was assembled by using information from the literature and two public resources, i.e., UniProt (16) and the Human Protein Atlas (HPA) (17,18) (**Supplemental File 1**). UniProt proteins were derived by using the advanced search interface that returned 5,440 protein IDs. Controlled vocabulary terms were used for searching the cellular compartment (CC), Gene Onthology (GO) and the Keyword (KW) fields, all filtered for cell membrane localization. Lists of proteins localized to the cell-membrane, cell-surface, cell junction, cell projection, and peripheral proteins, with roles in signaling (i.e., receptor and catalytic activity), immune response (e.g., CD antigens), adhesion, and transport, were extracted. Plasma membrane proteins from the HPA were retrieved in bulk (2,068 IDs), and additional cell-surface proteins detected *in-vitro* by using various experimental enrichment techniques (12,13) or predicted via *in-silico* studies (14,15) were added to the list (4,717 IDs). Complementary information about GPCR families was acquired through IUPHAR (19). A classification of the cell-membrane proteins included in the database is presented in **Figure 1A**. The overlap between the various protein categories was rather minimal (~10-15 %), but unavoidable, due to the complex roles that the cell-membrane proteins play in several biological processes. With improvements in sample preparation technologies, MS detection sensitivity, and search engine machine learning capabilities, a more consistent consensus between the various information sources is also expected.

**Figure 1.**
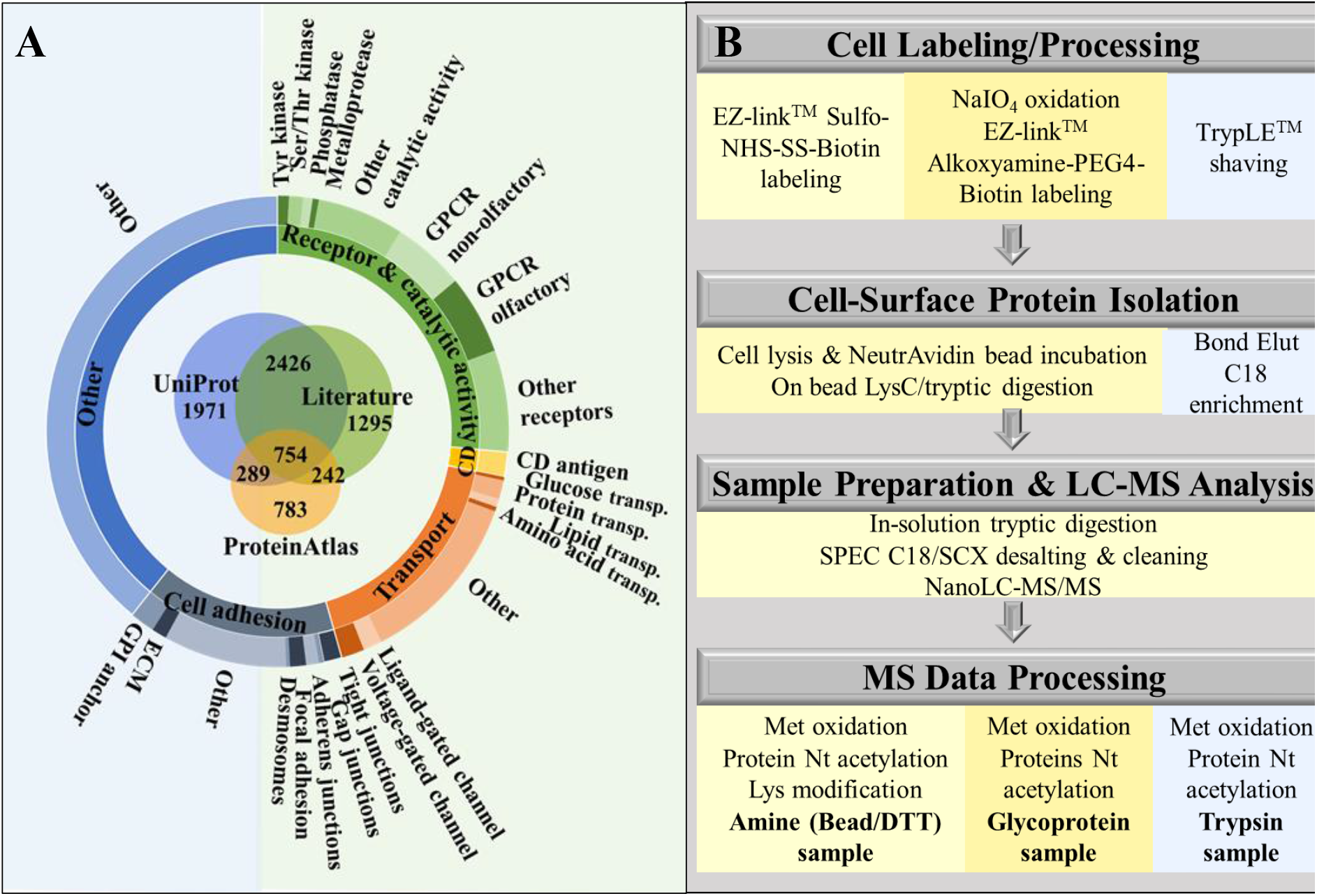
Cell-membrane protein database classification and isolation flowchart. **(A)** In-house built database of 7,760 cell-membrane proteins classified based on GO controlled vocabulary terms. **(B)** SKBR3 cell-membrane protein isolation and processing workflow via three distinctive methods: biotin labeling of protein primary amine groups, biotin labeling of glycoproteins, and enzymatic shaving.

### Effectiveness of cell-membrane protein isolation

To isolate the cell-membrane fraction of SKBR3 cells, a combination of chemical labeling and enzymatic approaches was followed (**Figure 1B**). Based on reported yields and processing times (8–13), three methods, relying on protein isolation by biotinylation and affinity pulldown of amino groups and of glycan posttranslational modifications, as well as on tryptic shaving of receptors in cell culture, were chosen. Sulfo-NHS-SS-biotin based isolation of proteins enabled the labeling of primary amino groups at the protein N-terminal (α-amino) and Lys (ε-amino) residues, while alkoxamine-PEG4-biotin based isolation of carbohydrate moieties that are commonly encountered on the cell-surface proteins. Trypsinization of cells in culture was the least time-consuming method due to minimal processing prior and after sample collection.

The efficiency of the biotin labeling reaction was evaluated in a separate study by using two BSA protein samples and SKBR3 tryptic peptides. The sample:Sulfo-NHS-SS-biotin molar ratios for the BSA protein samples were 1:56 and 1:560, respectively, and for the SKBR3 tryptic peptides was 1:10 (**Figure 2A**). The labeling efficiency of the tryptic peptides was very high, reaching ~99 % at the N-terminal (Nt) and ~82 % for all Lys (K)-containing peptides. Only ~1.5 % of the peptides were non- or partially labeled. However, when labeling was performed at the protein level and followed by proteolytic digestion, the labeling efficiency of Lys-containing BSA peptides dropped progressively to ~75 % and ~54 %, respectively, with the decrease in the molar ratio of the added biotinylation reagent. This was matched by a concomitant increase in the non-labeled or partially labeled peptides to 52 % and 69 %. N-terminal labeling of multiple BSA peptides was observed, as well, presumably due to incomplete quenching of the labeling reagent prior to proteolytic digestion. Nonetheless, the labeling of the BSA protein N-terminal amino acid could not be detected. The results underscore the impact of limited reagent accessibility to hindered Lys sites in a protein, which becomes a much more challenging factor in the case of live cells when the extracellular domain of intact membrane proteins often displays a heavily modified and entangled structure (26). Accordingly, Lys biotinylation of cell-surface proteins on live cells was observed at a substantially reduced level (~1.9 %), even less than that of Met oxidation (~4.5 %). Protein N-terminal acetylation was also detected at a low level (~0.4 %), but biotinylation was not observable (**Figure 2B)**. As a result, in the final analysis, the biotinylation-induced modification on the N-teminus of proteins was not included in the list of enabled DB search modifications. In the case of alkoxamine-PEG4-biotin-based labeling and isolation of glycosylated proteins, the labeling efficiency could not be evaluated by MS as there was no change in mass involved, however, the attachment of the labeling reagent to the cell-surface proteins could be visualized by microscopy and indicated a uniform coverage (**Figure 2C**).

**Figure 2.**
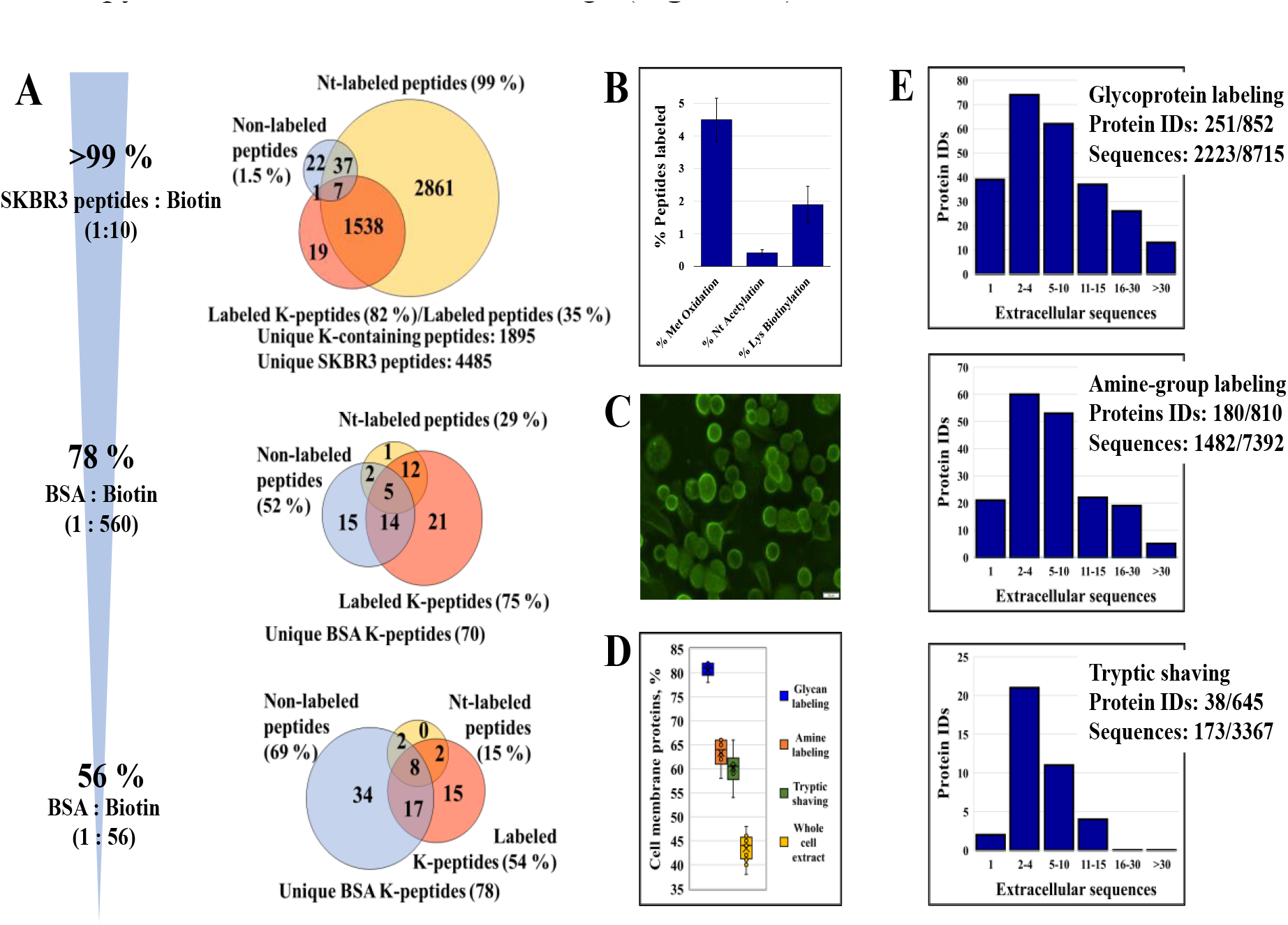
Cell membrane protein labeling efficiency and enrichment in extracellular sequences. (**A**) Percent peptides carrying a biotinylation-induced label in an SKBR3 cell extract and in BSA tryptic digests using various peptide/protein:biotin molar ratios (applicable to the amine group labeling method); the Venn Diagrams represent the number of labeled Lys-containing peptides (+87.998 Da), peptides labeled at the N-terminus (+87.998 Da), and the number of non-labeled peptides. (**B**) Percent peptides carrying a PTM: Met oxidation, peptide Nt acetylation, and Lys biotinylation (case of the amine group labeling method with on-bead proteolytic digestion); the error bars represent the SD of biological replicates. (**C)** SKBR3 cells labeled by alkoxyamine-PEG4-biotin and conjugated with streptavidin antibody-Alexa Fluor™ 488. (**D**) Cell membrane protein enrichment effectiveness represented by the number of cell membrane proteins in the top 100 most abundant proteins (abundance determined by the number of matching unique peptides). (**E**) Histograms of protein IDs matched by different numbers of extracellular peptide sequences; (with extracellular sequences/total detected); extracellular sequence assignments were made based on topological domain information extracted from UniProt.

The enrichment efficiency in cell-membrane proteins was assessed by MS, by calculating the proportion of membrane proteins in the top 100 most abundant proteins, with abundance defined by the counts of unique peptides per protein. According to the controlled vocabulary annotations in the database that was described above, the percentage of cell-membrane proteins in typical whole cell extracts was ~43 %. Upon enrichment, this percentage increased to ~60 %, ~64 % and ~81 % for tryptic shaving, amino group, and glycan labeling, respectively (**Figure 2D**). Cell-surface protein enrichment based on glycan labeling provided the highest yield, most likely due to the heavy glycosylation of the extracellular protein domains that could be more efficiently labeled than the protein N-termini and Lys residues in the case of the amine labeling method (27). Poor penetration of trypsin through the cell-surface protein coat, slow tryptic activity, and the possible contribution of lysed cell content to the pool of identified proteins may have led, on the other hand, to the lowest enrichment yield for the tryptic shaving method. For proteins for which topological information was available in UniProt (i.e., for 2,923 proteins from the in-house built DB, matched by 7,546 extracellular sequences), the topological domain assignments validated the presence of numerous cell-membrane proteins from the dataset. The histograms from **Figure 2E** indicate that many of the detected proteins were identified by multiple extracellular sequences, confirming thus their presence in the cell-membrane or on the cell-surface, and also that in comparison to trypsinization the chemical labeling methods were more effective for capturing the cell-membrane proteome.

### Cell-membrane proteome data analysis

The combination of orthogonal enrichment approaches led to the identification of a total of 2,054 cell-membrane proteins, 1,921 in the serum-free (SF) and 1,435 in the serum-treated (ST) cell states, of which 1,316, 1,254 and 1,030, respectively, were identified by at least two unique medium or high confidence peptides (**Figures 3A/3B, and Supplemental File 2**). The three methods were complementary to each other, however, as shown in the Venn diagrams from **Figure 3C**, cell-membrane protein enrichment based on the labeling of glycoproteins enabled the identification of the largest number of proteins and with the best reproducibility, i.e., 65-67 % overlap between three biological replicates. Protein isolation based on Sulfo-NHS-SS-biotin labeling of amino groups performed the worst (~40-47 % overlap), likely due to less efficient labeling of proteins. The reproducibility of the tryptic shaving method was somewhat higher than that of the amino group labeling approach (~52-54 % overlap), nevertheless, the effectiveness in isolating cell-membrane proteins was the worst (**Figure 2D**). In terms of peptide spectrum matches, the quality of protein identification was high and consistent across all three methods with correlation factors ranging from 0.95 to 0.99 (~0.9 for some tryptic samples) (**Figure 3D**). Despite differences in effectiveness and reproducibility, the use of orthogonal enrichment methods facilitated the identification of a larger set of proteins and contributed to a more detailed characterization of the cell-surface proteome.

**Figure 3.**
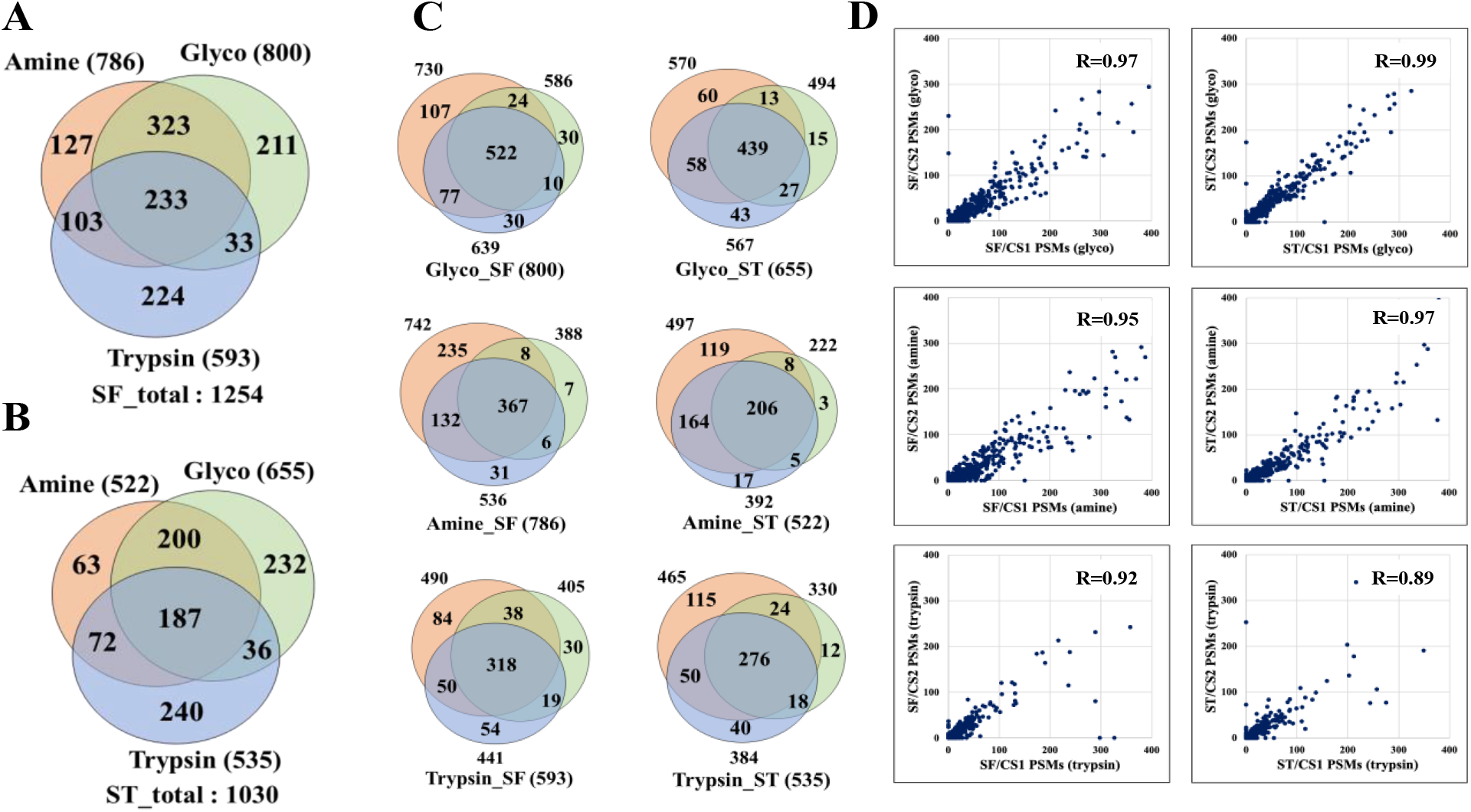
Venn and correlation diagrams representing the complementarity and reproducibility of the three labeling methods in detecting cell-membrane proteins matched by at least 2 unique peptides (FDR< 3 %). (**A**) Serum-free cultured cells. (**B**) Serum-treated cells. (**C**) Reproducibility of protein detection between three biological replicates for each labeling method and cell treatment conditions (SF and ST). (**D**) Correlations of peptide spectrum matches (PSMs) between any two biological replicates for each of the three cell-membrane protein enrichment methods; the correlations are shown for the 0-400 PSM range in which the vast majority of proteins could be found (R=Pearson correlation coefficient).

### The SKBR3 surfaceome

The remarkable ability of cancer cells to enact aberrant proliferation programs and metastasize to distant sites is mediated via an altered cell-surface proteome that facilitates in-and-out cell signaling processes as well as adhesion and migratory functions. To gain a better insight into these processes, a functional characterization of the SKBR3 surfaceome was performed by categorizing the set of 1,316 proteins into four major groups: receptors and proteins with catalytic activity, transporters, cell adhesion/junction proteins, and proteins with immune functions such as CDs. Less abundant categories included receptor substrates and cell-surface binding proteins (e.g., peripheral membrane proteins, GPI anchors, matrix metalloproteinases/MMPs, and ECM molecules). The dendogram from **Figure 4A** provides an overview of a representative subset of 525 cell-membrane proteins that could be placed in specific compartments, with protein IDs being included in **Supplemental File 3**. Cell-surface protein enrichment by glycan or amino group labeling yielded the largest number of receptor/catalytic proteins and CDs (**Figures 4B and C**), while trypsinization enabled the identification of a more abundant fraction in cell adhesion and transport proteins (**Figure 4D**).

**Figure 4.**
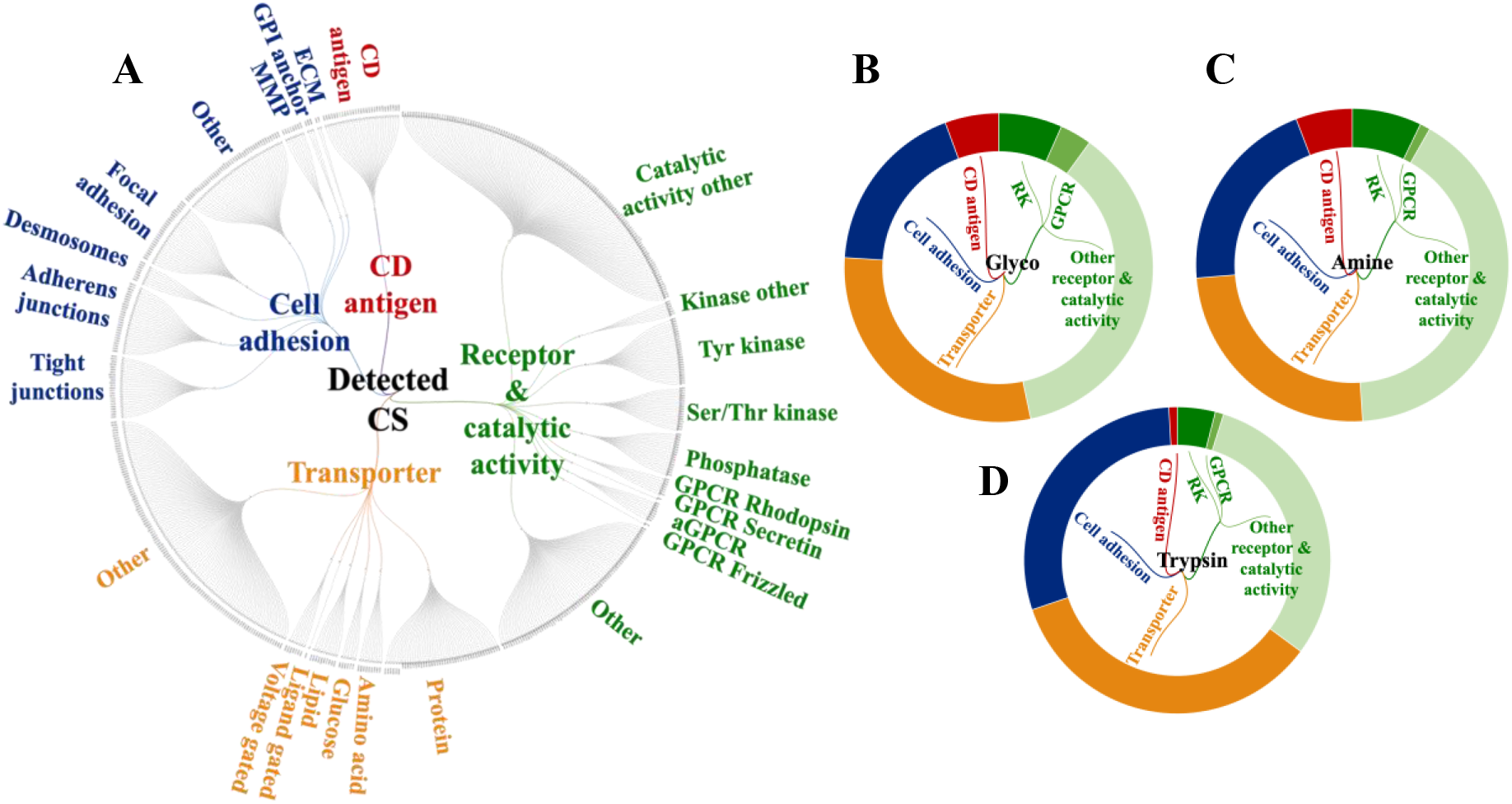
Functional categorization of the detected cell-membrane proteins based on GO controlled vocabulary terms (proteins detected by at least 2 unique peptides, FDR<3 %). (**A**) Dendrogram of cell-membrane proteins detected by all labeling methods and conditions. (**B**), (**C**), and (**D**) Doughnut charts of detected cell-membrane proteins enriched via glycoprotein labeling, amino group labeling, and tryptic shaving methods, respectively.

To further assess biological utility, the combined results of the three enrichment methods were compared to the output of three additional independent experiments, one including cell-surface protein enrichment from proliferative cells, and two including whole cell analysis of SF and ST cells without enrichment in cell-surface proteins (**Table 1**). As expected, when enrichment was performed, a larger number of cell-surface proteins were identified. A clear advantage, however, was observable only when proteins matched by two unique peptides were counted. In particular, the enrichment process enabled the high confidence detection of a much larger number of signaling receptors (GPCRs, Tyr receptor kinases), GPI anchors and CD antigens (columns 5,6 vs 7,8). Notably, the glyco enrichment method alone enabled the identification of the vast majority of kinase/GPCR receptors and CD antigens, rendering it, therefore, the method of choice for profiling valuable targets for therapeutic treatment and immunophenotyping (**Figure 5A**). In contrast, cell-membrane Ser/Thr kinase receptors were detectable in higher numbers and with a larger number of unique peptides without performing cell-surface protein enrichment, likely due to the prevalently longer cytoplasmic tails in comparison to the shorter extracellular N-terminal domains. The Protter visualization of one representative member of Tyr kinase, Ser/Thr kinase and GPCR receptor categories highlights this trend (**Figure 5B**).

**Figure 5.**
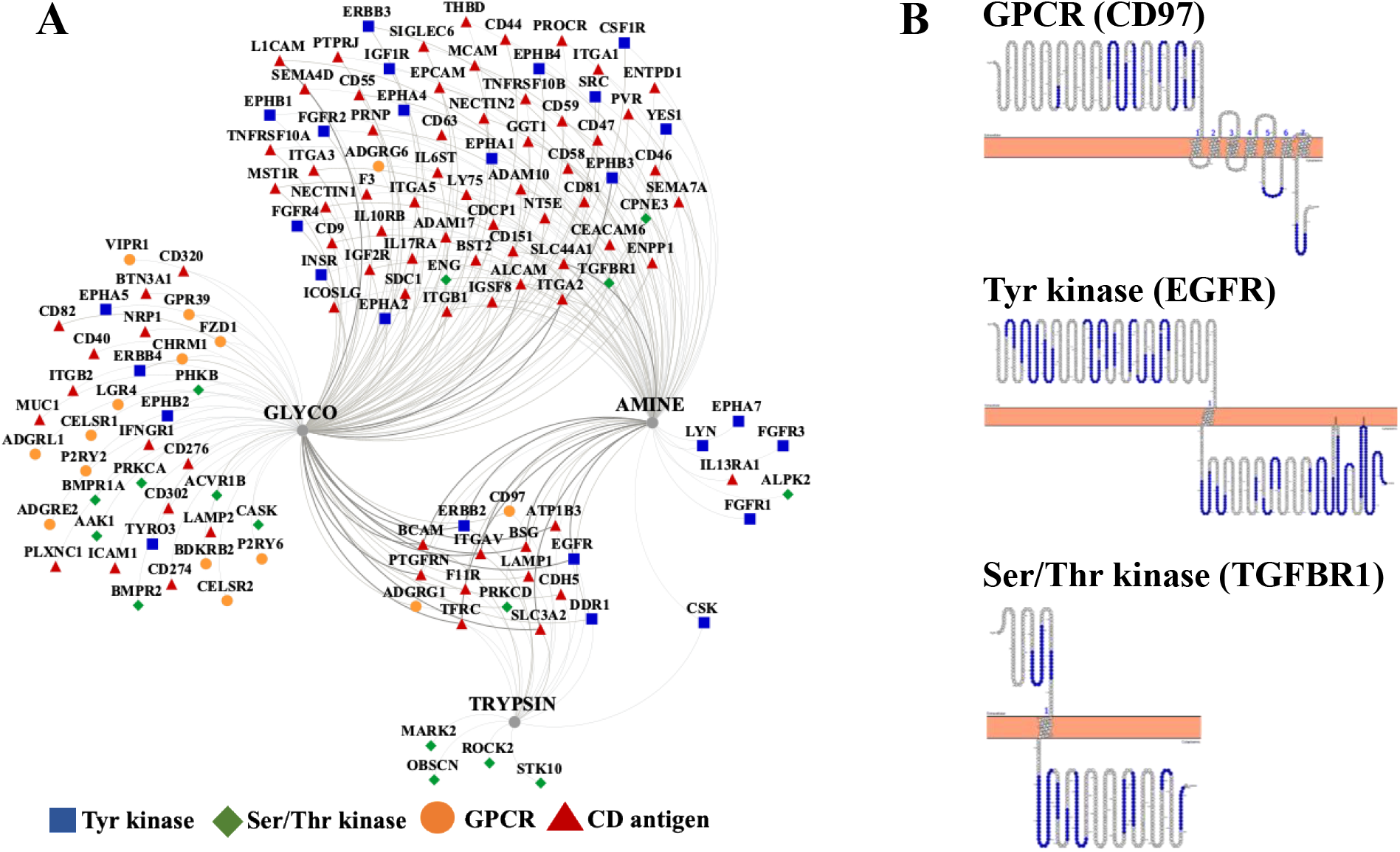
Detectability of cell-membrane receptors and CD proteins. (**A**) Diagram representing the overlap between the detected proteins produced by the three enrichment methods (visualization performed with Cytoscape); the edge thickness reflects the protein abundance as evidenced by PSMs. (**B**) Protter visualization of exemplar protein topologies in the cell membrane for three classes of receptors: Tyr kinase (EGFR), Ser/Thr kinase (TGFBR1), and a GPCR (CD97).

**Table 1.** Cell-surface protein identification effectiveness with and without enrichment in cell-surface proteins, by considering the whole protein set or only proteins matched by two unique peptides.

|  | G1+S<br>cells<br>(CS) | Prolife<br>rating<br>cells<br>(CS) | G1+S<br>whole<br>cells* | G1+S<br>whole<br>cells<sup>#</sup> | G1+S<br>cells<br>(CS)<br>2 pep | Prolife<br>rating<br>cells<br>(CS)<br>2 pep | G1+S<br>whole<br>cells*<br>2 pep | G1+S<br>whole<br>cells<sup>#</sup><br>2 pep |
| --- | --- | --- | --- | --- | --- | --- | --- | --- |
|  | 1 | 2 | 3 | 4 | 5 | 6 | 7 | 8 |
| Matches to<br>CSDB | 2054 | 1359 | 2265 | 2821 | 1316 | 1175 | 1295 | 1108 |
| Matches to<br>CS/UniProt | 1339 | 874 | 1427 | 1831 | 861 | 754 | 772 | 667 |
| Receptors | 168 | 112 | 114 | 211 | 117 | 106 | 24 | 21 |
| GPCRs | 36 | 15 | 36 | 65 | 15 | 11 | 1 | 1 |
| Tyr kinases | 29 | 20 | 16 | 30 | 26 | 20 | 8 | 4 |
| Ser/Thr<br>kinases | 29 | 17 | 40 | 44 | 17 | 15 | 25 | 19 |
| Transport(ers) | 381 | 275 | 348 | 425 | 279 | 247 | 198 | 163 |
| Cell junction/<br>cell adhesion | 348 | 251 | 347 | 421 | 255 | 231 | 187 | 158 |
| GPI anchors | 25 | 21 | 16 | 28 | 17 | 20 | 4 | 2 |
| Signal anchor | 24 | 21 | 28 | 35 | 20 | 19 | 12 | 11 |
| CD antigens | 105 | 84 | 58 | 84 | 89 | 80 | 19 | 13 |
| Secreted | 643 | 470 | 718 | 897 | 454 | 401 | 396 | 366 |

\*Data acquired with a QExactive Plus Orbitrap mass spectrometer. <sup>#</sup>The analysis included many replicates. Greyed columns include protein count data from SKBR3 cells subjected to enrichment in cell-surface proteins. Red cells indicate conditions under which a substantial improvement in cell-surface protein enrichment was observed.

Altogether, based on controlled vocabulary terms, the enrichment process enabled the classification of about 275 proteins with catalytic and receptor activity (including 15 GPCRs, 26 Tyr kinases, 17 Ser/Thr kinases), 89 CD antigens, 255 cell adhesion/junction molecules, and 279 transport proteins (**Figures 6A** and **7**). An interrogation of the biological processes (**Figure 6B**) and associated pathways (**Figure 6C**) represented by these proteins revealed for each category not just one, but multiple and complex roles with broad impact on essential cellular processes such as cell communication/signaling, biological adhesion and migration, transport, immune response, cell growth, death and differentiation (**Supplemental File 2**). Beyond these four categories, additional groups included proteins that were either associated with the cell-membrane (i.e., peripheral membrane, cell junction, cell projection, ECM) or represented contamination from other compartments (secreted and exosome proteins).

**Figure 6.**
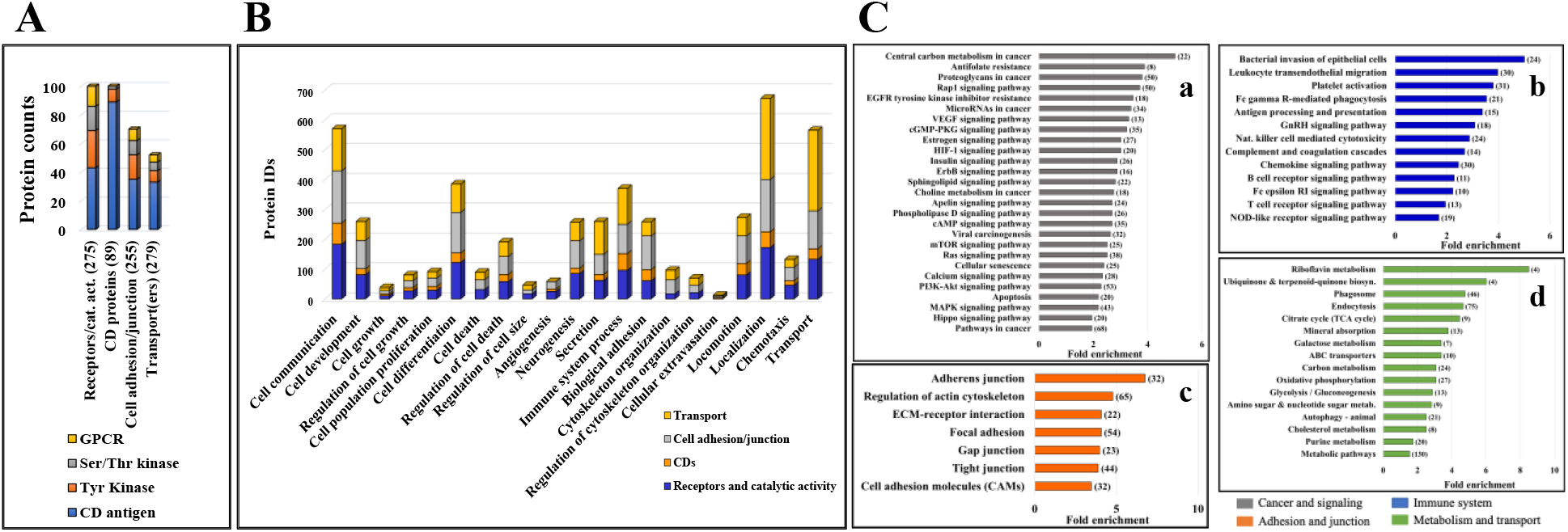
Bar charts of selected functional categories and pathways associated with the detected cell-membrane proteins. (**A**) Categorization of the detected receptors/enzymes, antigens, transporters and cell adhesion/junction proteins into Tyr kinase, Ser/Thr kinase, GPCR and CD groups. (**B**) Cancer-relevant enriched biological processes represented by the cell-membrane proteins. (**C**) Enriched KEGG pathways represented by cell-membrane proteins involved in (**Ca**) signaling and cancer, (**Cb**) immune response, (**Cc**) adhesion/junction, and (**Cd**) metabolism and transport. Notes: Numbers in brackets represent the number of proteins matched to each process; full lists, fold-enrichment and FDR values are provided in **Supplemental File 2**; the background gene sets were the full set of corresponding proteins in the human proteome.

**Figure 7.**
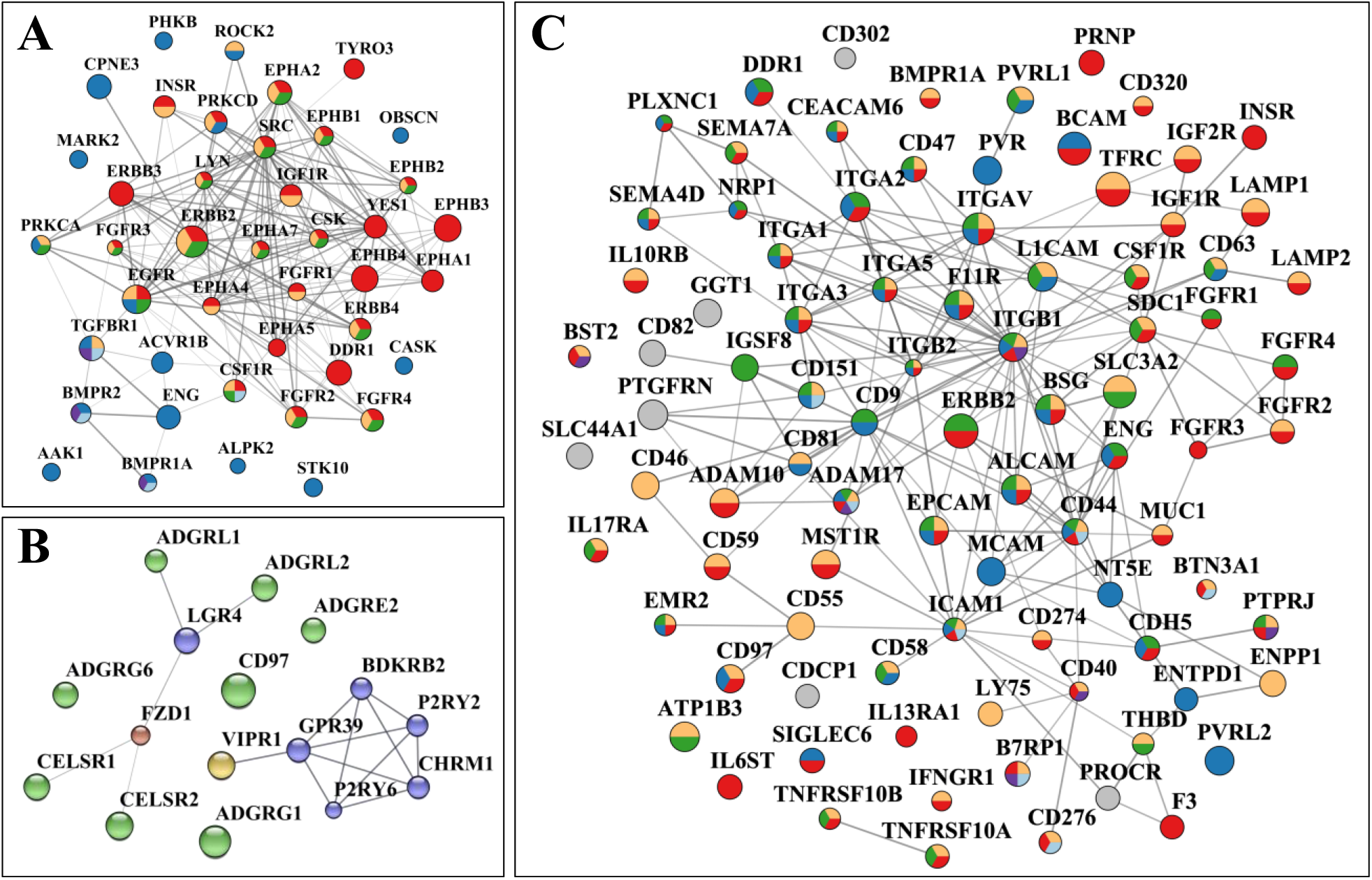
Protein-protein interaction networks of selected cell-membrane protein categories. (**A**) Receptor Tyr and Ser/Thr kinases. Red-Tyr kinase receptor, Blue-Ser/Thr kinase receptor, Yellow-MAPK regulation, Green-ERK1/ERK2 regulation, Light blue-Cytokine-cytokine receptor signaling, Magenta-TGFB signaling. (**B**) G-protein coupled receptors. Blue-Class A Rhodopsin, Yellow-Class B1 Secretin, Green-Class B2 Adhesion, Red-Class F Frizzled. (**C**) CD antigens. Yellow-Immune system process, Blue-Biological adhesion, Red-Cell communication, Green-Locomotion, Magenta-B cell activation, Light blue-T-cell activation. Notes: The PPI networks were generated with STRING and visualized with Cytoscape; node size is proportional to the total spectral counts that matched a protein, from <10 (small) to >10,000 (large).

### Proteins with receptor and catalytic activity

The category of cell-membrane proteins with catalytic and receptor activity included, in addition to a large group of kinase receptors, non-receptor kinases, phosphatases, MMPs, GTPase molecular switches, and proteins with ATPase activity. Together, these proteins are engaged in extensive cell-to-cell signaling and intracellular signal transduction, cell growth, apoptosis, cell locomotion and migration, trafficking of various cellular components, transport (ions, lipids, amino acid, metabolites), and regulation of actin cytoskeleton organization and cell polarization.

Enzyme-linked receptors display extracellular domains for binding growth factors, cytokines or hormones, and initiate the transmission of chemical signals via their intracellular cytoplasmic domains that either have, or interact with proteins that have, catalytic activity. These receptors play major roles not just in signal transduction, but also in cell proliferation, differentiation and development. They also represent widespread drug targets due their aberrant signaling activity linked to many diseases including inflammation, metabolic disorders, and cancer (**Figures 6B** and **6Ca**) (6). Among the detected catalytic receptors, the most relevant to cancer growth, proliferation and differentiation, and breast cancer specifically, were the Tyr protein kinases of the EGFR/ERBB, FGFR and IGFR families of growth factor and hormone binding receptors. These receptors form highly interconnected PPI networks thorough which they control essential cellular functions (**Figure 7A**), are highly mutated in many cancers (28), and represent promising tumor markers and/or drug targets (**Figure 8**). Mutations that lead to gain of function, genomic amplification or chromosomal rearrangements are often responsible for abnormal activation, signaling, and uncontrolled cell proliferation. Among the newest RTK drug targets in breast carcinoma, the discoidin domain receptor (DDR1) involved in the activation of cell proliferation, survival, ECM remodeling, migration and invasion pathways, was detectable (29). In addition, Ser/Thr kinase receptors for a number of TGF-β superfamily of ligands (BMPR1A/BMPR2 bone morphogenic proteins, ACVR1/ACVR2 activin receptors), as well as members of the TGFBR complex (ENG), were present. The activation of TGF-β receptors is controlled by interactions with other proteins and by various posttranslational modifications (e.g., phosphorylation, ubiquitylation, sumoylation, and neddylation). As a result, these receptors are capable of triggering downstream signaling processes via multiple pathways, including SMAD, ERK, JNK and p38MAPK (30). Modulation of TGFBR signaling is accomplished by interactions with a broad range of cell surface receptors and non-receptors, for example, with ENG, NRP1, PDGFRβ, CD44, and integrins, which were all detected in the cell membrane fraction of SKBR3 cells.

**Figure 8.**
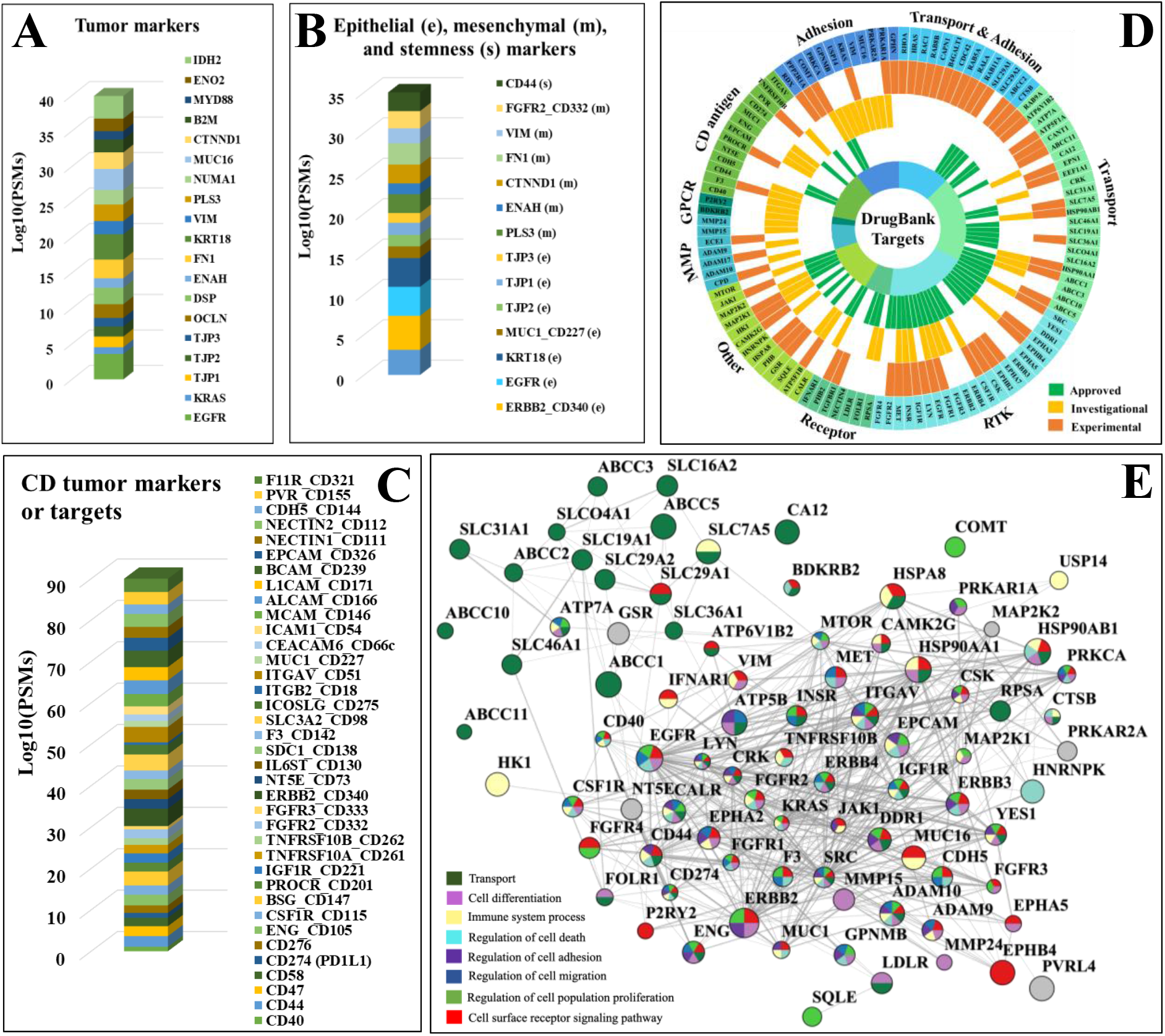
Cancer markers and drug targets detected in the SKBR3 cell-membrane proteome. (**A**) Tumor markers. (**B**) Epithelial (e), mesenchymal (m), and stemness (s) cancer markers. (**C**) CD Tumor markers and drug targets. (**D**) Sunburst chart representing receptors by functional categories identified in DrugBank as potential cancer therapeutic drug targets based on approved (green), investigational (yellow), and experimental (orange) status levels. (**E**) PPI network of the approved and investigational cancer drug targets. Note: Node size is proportional to the log10 spectral counts and edge thickness reflects the STRING interaction score.

A fairly large group of Tyr kinase ephrin receptors, semaphorin plexin receptors, neuropilin (NRP1), and ROBO1 formed a cluster with multiple important roles in developmental processes, cell differentiation, cytoskeleton remodeling and migration. Ephrins are known to be involved in the regulation of multiple signaling pathways (e.g., MAPK, ERK, RAS, estrogen), and together with ROBO1 and NRP1 (a VEGF receptor) play major roles in angiogenesis and vascular development (16,31). Plexin receptors, on the other hand, have been shown to be involved in invasive cell growth and ERBB signaling. For example, PLXNB2 promotes the phosphorylation of ERBB2 at Tyr 1248 (16), a phosphorylation site that downstream affects biological processes related to cell cycle and growth, cytoskeleton organization, motility, apoptosis, and carcinogenesis (32). Together with FGFR1, EGFR/ERBB, HMGB1, BMPR1A/BMPR2, and the non-catalytic NOTCH1/2/3 group, the plexin/ephrin receptors are further massively involved in cell differentiation processes. In addition, NOTCH signaling has been shown to be aberrantly activated in breast cancers via either receptor overexpression or mutations that lead to uncontrolled cell proliferation and survival (33).

Altogether, the ephrin/plexin group was part of a larger functional cluster with roles in chemokine signaling and increased chemotaxis, that included not just well-known receptor/non-receptor signaling kinases (ERBB2, CSF1R, SRC, PRKCD), but also a group of proteins with functionally diverse activities [i.e., small GTPases (RHOA, RHOG, HRAS, RAC1, RAB13), signaling adapters (SHC1), receptor-type tyrosine-protein phosphatases (PTPRA, PTPRJ), nectin receptors (PVRL1), receptor and non-receptor proteins with roles in immune responses (CSF1R, IL17RA, IL17RC, LYN, CXCL1, CXADR), integrin receptors (ITGA1, ITGAV, ITGB2), cell adhesion and cell-ECM binding molecules (EMR2, L1CAM, ALCAM, HMGB1, NEO1, DAG1), Ca-binding proteins (S100A8/9, PRKCA, spectrins), cation channels (TRPMs), and other cytoskeleton reorganization proteins (VASP, ENAH, MYH10, EZR, CORO1A/1B)]. Chemotactic cell migration relies on a signaling system that uses cell-surface receptors to detect and respond to extracellular gradients of chemical cues (e.g., CSF1). In tumors, an altered expression of chemokines is responsible for the recruitment of immune cells and for cellular processes that support angiogenesis, proliferation, cancer dissemination and metastasis (34). As a result, chemokines and their cognate chemokine receptors evolved as valuable drug targets for the development of novel immunotherapeutic interventions (35).

### GPCRs

GPCRs encompass ~800 protein members and are the largest family of cell-surface receptors (36). GPCRs transduce extracellular signals by changing their conformation upon binding of a ligand and transmiting the signal through G-protein modulation. The detected GPCRs were part of the Rhodopsin (class A) and Secretin (Class B) families (**Figure 7B**), and were yielded mainly by the glycan labeling method likely due to their rather low abundance and high glycosylation rate at the N-terminal sequences (37). **Supplemental file 4** provides PRM/MS data that validate the presence of the detected GPCRs. These GPCRs included adhesion (aGPCRs), Wnt signaling and neuroactive ligand-receptors, several with EGF-like (CELSR1, CELSR2, CD97, EMR2) and hormone-receptor (VIPR1, CELSR1, LPHN1) Pfam domains. Most Secretin GPCRs were also adhesion GPCRs that contained GPCR proteolysis sites (GPS). The aGPCRs are evolutionarily conserved (37), and, in addition to their involvement in cell adhesion and migration processes, have emerged for their role in tumorigenesis and metastasis (38). For example, ADGRE5 (CD97) and ADGRG1 (GPR56) are two aGPCRs that have been detected by the largest peptide counts and by all enrichment methods, and have been reported for increased expression in various cancers (38,39). The Rhodopsin class A GPCRs (P2RY2, P2RY6, CHRM1, BDKRB2, GPR39) included a cluster involved in neuroligand-receptor ligand interactions that also comprised two G protein-coupled purinergic nucleotide receptors interacting with Vasoactive intestinal polypeptide receptor 1 (VIPR1). P2RY6 is a known target for colorectal cancer due to its role in protecting cancer cells from apoptotic processes, possibly via AKT and/or ERK1/2 signaling, but less is known about its role in breast cancer (40).

Due to their comprehensive role in a wide range of signaling processes and physiological conditions, the GPCRs also represent one of the largest families of drug targets. The majority of aGPCRs lack, however, an endogenous ligand and their mechanism of action is not fully understood. Emerging crosstalk activity between GPCRs and catalytic receptors (e.g., RTKs such as EGFR) has revealed novel signaling mechanisms with roles in cell proliferation and differentiation (41), the GPCRs being capable of initiating distinct MAPK signaling pathways via stimulation of ERK, JNK and p38MAPK (42). Depending on the transduction mechanism, however, the GPCRs can have either an inhibiting or stimulating role on the downstream pathways, and the mechanistic details of RTK transactivation are yet to be explored (41). To boost the discovery of novel therapeutic targets, the challenge of transactivation studies is twofold, i.e., to demonstrate the presence of existing crosstalk interactions and to clarify the relevance of such crosstalk to disease.

### Immune system receptors, CDs, and antigen characteristics

Cancer cell receptors that trigger cytotoxic innate and adaptive immune system responses are critical to the path of tumor development, and are key determinants of the biological processes that help cancer cells evade destruction by immune attack. The presence of a group of interleukin and interferon cytokine receptors (ILs, IFNs), C-lectin/Fc/scavenger receptors, macrophage stimulating protein receptors (MST1R, CSF1R), HLA class I histocompatibility antigens (HLA-E, HLA-G), and B-cell/T-cell activating proteins (LYN, CD40, CD81) were indicative of SKBR3 triggers capable of eliciting a spectrum of innate, adaptive and inflammatory reactions that included among others cytokine production, positive regulation of innate immunity and defense responses, and elicitation of B-cell proliferation and T-cell killer cytotoxic effects (**Figures 6B** and **6Cb**). Many of these proteins were part of a group of 89 cluster of differentiation (CD) antigens with multiple roles not just in immune system processes but also in cell communication, signal transduction, adhesion, cell locomotion and transport (**Figure 7C**). The detected CD antigens encompassed classical receptors (ERBB2, FGFRs, IGFRs, TNFRs, TFRC, IL and IFN receptors), integrins and integrin binding proteins (ITGAV, ITGAs, ITGBs, semaphorins), cell adhesion molecules (CAMs: EPCAM, BCAM, ICAM1, L1CAM, MCAM, CDH5/CD144, mucins, nectins), and disintegrin metalloproteinase domain-containing proteins (ADAM10/17). The relevance of this cell-surface category was underscored by participation in- or regulation of pathways such as MAPK, PI3K-AKT, ERK, JAK/STAT, ECM-receptor interactions, and B-cell/T-cell activation. The most abundant CDs included members of all protein categories (ERBB2, TFRC, ITGAV, ITGA2, ITGB1, BCAM, EPCAM, SLC3A2), with high degree centrality nodes being represented primarily by adhesion proteins (ITGB1, ITGAV, ICAM1, CD9, and CD44) (**Figure 7C**).

Several CDs are documented tumor biomarkers or drug targets (**Figures 8A-C**) (43–47), but of particular interest was the presence of antigen immunological markers that define the epithelial, mesenchymal or stemness characteristics of cells (**Figure 8B**). A group of 8 epithelial (EpCAM/CD326, ERBB2/CD340, EGFR, KRT18, MUC1/CD227, TJP2, TJP1, TJP3), 6 putative mesenchymal (PLS3, ENAH, CTNND1, FN1, VIM, FGFR2/CD332) and one stemness marker (CD44) were present, with the ERBB2, EGFR, KRT and EpCAM epithelial markers being highly abundant on the cell-surface (43). The presence of non-epithelial markers, however, indicated that the SKBR3 cells were a mixed population of differentiated epithelial cells and cells undergoing EMT with stemness characteristics (43). It must be noted, though, that from the detectable peptide sequences it was not clear whether the mesenchymal or epithelial splice variants and protein isoforms of ENAH and FGFR2 were identified (43). The presence of CD44 was also reflective of the metastatic characteristics of SKBR3 cells, while that of PDL1/CD274 (programmed cell death 1 ligand), a receptor ligand that blocks T-cell activation and that is upregulated by many tumor cells, of the ability to escape immune surveillance (48). The PD1/PD1L1 receptor/ligand pair is the target of thousands of clinical trials that test immune checkpoint inhibitors (49). Possible similar roles have been attributed to CD276, as well.

### Cell adhesion and junction proteins

These molecules are often in a gray area of categorization because they participate not just in cell adhesion and locomotion, but also in a broad range of cancer-relevant processes including cell-cell and intracellular signaling, cell growth/proliferation/differentiation and death, secretion, angiogenesis, endocytosis, and chemotaxis, just to name a few (**Figures 6B** and **6Cc**) (50). The CAM receptors that are involved in signaling belong to several families that include Ca-dependent cadherins, integrins, selectins, and Ca-independent immunoglobulin-like proteins. CAMs do not have catalytic domains, but engage in signaling by association with signaling adaptors and nonreceptor tyrosine kinases. Adhesion molecules that use non-enzymatic mechanisms for signal transduction have been, however, much less studied with respect to the details of signal recognition and transfer. All classical cell-matrix (focal adhesion integrins) and cell-cell adhesion categories (adherens junction cadherins and protocadherins; desmosomal desmocollin, desmoglein, and plakins; tight junction claudins, occludins, JAMs, and ZO proteins; gap junction connexins), as well as a number of additional CAMs (EPCAM, BCAM, MCAM, CDH5/CD144, mucins), immunoglobulin-like CAMs (ICAM1, L1CAM, nectins), disintegrin (ADAM 9/10), and matrix metalloproteinases (MMP15/24) were represented in the dataset.

The integrins and selectins have been shown to be involved in various aspects of the metastatic process (51). Integrins are transmembrane adhesion receptors that recognize a variety of cell-surface or extracellular matrix (ECM) ligands (e.g., fibronectin, vitronectin, laminin, and collagen). The binding is mediated by the 24 α- and 9 β glycoprotein subunits that form noncovalent heterodimers (ITGA/ITGB) with binding activity modulated by various extracellular (e.g., Ca^2+^/Mg^2+^) or cell-type specific factors, and affinity for either cell-matrix or cell-cell interactions. Integrins have multiple and complex roles in cancer progression, and distinct integrin expression patterns were used to predict survival and organ-specific metastases via tumor-derived exosome uptake (52). When activated by the binding of matrix components, the integrins can engage catalytic receptors and co-operate in triggering intracellular signaling pathways that regulate cell growth, survival, proliferation and differentiation. Viceversa, signaling processes initiated by conventional receptors can alter the expression and ligand-binding properties of integrins. By mediating the interactions between the ECM and the actin cytoskeleton, via binding intracellular anchor proteins (α-actinin, talin, filamin, vinculin) and recruiting downstream focal adhesion kinase (FAK) and Src kinases, the integrins regulate cell shape, motility, and further, cell migration and invasion (53). The central role of integrins and CAMs in signaling, immune recognition and cell migration was underscored, as also highlighted above, by the high degree centrality of their nodes in the PPI network of CD antigens (**Figure 7C**). Selectins, on the other hand, are adhesion molecules found on the surface of leukocytes, platelets and endothelial cells that through interactions with ligands expressed on the surface of cancer cells (mucins, glycosaminoglycans or sulfated glycolipids) facilitate metastatic spread within blood vessels (51). Along with other receptors, many adhesion proteins that are tumor markers used in diagnostics and therapeutic decisions were detected, among which IDH2, MUC16 and ITGAV/CD5 in high abundance (**Figures 8A** and **C**) (47).

Cell-cell anchoring junctions were represented by adherence cadherin molecules associated with the actin filaments through cytoplasmic proteins such as catenins (e.g., CTNNA1), and desmosomal desmocollins (DSC2) and desmogleins (DSG2) bound to keratin intermediate filaments via plakin linkers (54,55). These types of junctions have roles in tissue morphogenesis, maintaining tissue architecture and epithelial homeostasis, cell proliferation and differentiation, and in facilitating cell movement. Therefore, the loss of these junctions is considered to be a prerequisite to epithelial-to-mesenchymal transition (EMT), migration and cancer invasion (55,56). Gap junctions-that allow for direct cell-to-cell communication to allow for the passage of small molecules, ions or electrical currents, and tight junctions-that have barrier function to prevent and regulate the passage of solutes, ions or water, have been also implicated in cancer. Altered expression of these junction proteins and damaged junction integrity or functionality have been associated with inflammatory conditions, anchorage-independent growth, EMT, cancer invasion and survival and growth at the metastatic site (57,58).

### Transporters and ion channels

The category of plasma membrane transport included members of the entire range of transport (i.e., ABCs-ATP binding cassette transporters and SLCs-solute carriers superfamilies) and ion channel proteins (i.e., ligand and voltage gated), as well as other receptor/signaling, adhesion, MMP or peripheral proteins that either have transporter activity or are adaptors or accessory components of the transport complexes. Altogether, the pool of membrane transport comprised 95 proteins with transporter activity and 22 with ion channel activity. Transporters carry solutes (nutrients, signaling molecules, metabolites, vitamins, drugs, ions) across the cell-membrane by relying on active transport processes with energy obtained by either ATP turnover or by exploiting ion gradients across the membrane. Ion channels control the selective passage of ions into and out of a cell through pore-forming membrane proteins, with gating provided by either a voltage gradient across the plasma membrane, the binding of intra- or extracellular ligands to the channel, or by mechanical stress. Defective transport has been correlated with a variety of metabolic diseases, and also with cancer (59). As ABC transporters mediate the efflux of drugs, they have been associated with the development of multidrug-resistance (MDR) and failure of chemotherapies. SLCs represent the 2^nd^ largest family of cell-membrane proteins after GPCRs that facilitate the transport of amino acids, peptides, sugars, neurotransmitters, vitamins, metals, inorganic/organic ions and electrolytes (59,60). Six ABC and 42 SLC transporters with symporter or antiporter activity were identified, mostly involved in ion transport, of which three were MDR-associated proteins (ABCC1, ABCC2, ABCC5, ABCC10) and two were relevant to drug uptake (SLCO4A1 and SLC22A18) (60,61). Moreover, among the 22 proteins with ion channel activity (seven voltage-gated and one ligand-gated), ORAI1, ANO1, STIM1, and PANX1 have been found to be involved in evasion of cancer cells from the primary tumor (62). Several recent reviews have summarized the therapeutic potential of cell-membrane transport proteins, and highlighted the anti-tumor or anti-metastatic potential of channel inhibitors (62,63). EMT was found to be associated with a remodeling of the Ca^2+^ signalosome (62), and voltage gated Na^+^ channels were found to be upregulated in breast cancer and to promote tissue invasion (63). Altogether, this group of proteins carries out functions that are related not just to cellular transport and localization, but also to signaling/communication, development/differentiation, immune response, secretion, and adhesion (**Figure 6B** and **6Cd**). The transport/adhesion proteins could be associated with additional functions related to cytoskeletal organization, the vesicle mediated transport with endocytosis, and the secretion proteins with immune responses (SLCs, ATPases, GTPases, TMEMs, TRPMs, MMPs, integrins, ORAI1, VAMP).

### Drug targeting potential

Given the immense therapeutic opportunities offered by cell-membrane proteins, the detected RTKs, GPCRs, CD antigens, MMPs, adhesion and transport proteins were searched in the DrugBank database to identify prospects for targeting HER2+ breast cancer cells (64). A total of 113 proteins were found in this pool, 56 for which approved and 48 for which investigational cancer drug targeting data existed (**Figure 8D** and **Supplemental file 5**). An additional category of 62 detected cell-surface proteins, not necessarily cancer-relevant, targeted by experimental drugs, was added to the list. The approved list included small molecule or monoclonal antibody drugs for various solid or liquid, early or advanced/metastatic cancers, and administered via chemo, targeted, combination, MDR, or topical therapeutic regimes. The RTKs, transporters and adhesion proteins represented the largest class of targets, followed by CD antigens, MMPs and GPCRs. In consensus, the targets of approved and investigational drugs clustered into two main categories, highlighting novel prospects for the development of anticancer drug cocktails (**Figure 8E**). One category encompassed regulators of signal transduction, cell proliferation, differentiation, adhesion/migration, immune response and death, while the other solute transporters through the cell-membrane. Despite the fact that GPCRs are the largest family of cell-surface and also druggable receptors, as observed, only few cancer therapies target the GPCRs. Generally, the discovery of novel drugs for GPCR targets has been limited by a high degree of sequence homology between many GPCRs at the binding site of ligands, and the lack of a clarified structure for GPCRs that are hard to isolate, purify or crystallize (65). Nevertheless, due to involvement in complex downstream signaling processes that lead to cancer proliferation and progression, GPCRs continue to be some of the most sought-after cell-surface targets (6). When contemplating, however, the large surfaceome map that can be revealed by proteomic studies, and the highly intertwined interactions between the various functional membrane protein categories (**Figure 8E**), network targeting of aberrantly behaving signature proteins evolves as a promising approach for the development of synergistic and more effective therapeutic approaches with less side effects and toxicity (66).

### Changes in cell-surface protein abundance

The regulation of plasma membrane protein abundances represents a key biological process through which the cells mediate intercellular communication, preserve cellular homeostasis, or exert their function in response to environmental stimuli. This regulation can be slow when it involves protein de novo synthesis or degradation, or fast when it relies on rapid removal or insertion of proteins from and into the plasma membrane by making use of proteins stored in endosomal compartments or exocytic vesicles (67). Given that harvesting of the cell-membrane proteome in this study occurred after prolonged exposure to SF or ST culture conditions, observable changes were expected to be representative of homeostatic processes rather than fast, transitory or cyclic events. Changes in cell-membrane protein expression between SF and ST cells were investigated after performing a two-step data normalization process, with the first, global normalization step being intended to account for data variability induced by sample processing, and the second step for the possible contamination of the cell-surface proteins by proteins from other cell compartments. Correction factors for global normalization were calculated by using the spectral counts of all proteins identified in each of the six samples under consideration (see Methods section), while for the second step, by using the spectral counts of only 10 endogenous cell-membrane proteins that met the following criteria: (a) the proteins were detected in every biological replicate of every isolation method, (b) the proteins were primarily associated with the cell-membrane but not with other cell fractions (i.e., nucleus, cytoplasm, ECM or secretome) and (c) the proteins had to have a transmembrane domain. The endogenous normalization proteins included solute carriers (SLC2A1, SLC3A2, SLC16A3), adhesion proteins (CDH5, PCDH1, DSG2, F11R), and receptors or proteins with catalytic activity (ITGB5, GNAS, SUSD2). The correction factors calculated by this approach ranged from 0.8 to 1.3 for the 1^st^ set, and from 0.94 to 1.06 for the 2^nd^ set. The method was applied to the dataset generated by the glycoprotein enrichment method that returned the largest number of total proteins IDs, with the highest reproducibility and enrichment effectiveness (i.e., 852 proteins with 2 unique peptides/protein). Changes in abundance were observed for members of all cell-membrane protein categories (**Figure 9A**), however, as the cell-surface protein labeling procedure induced the detachment and lysis of a small fraction of fragile, serum-starved cells, only proteins with cell-membrane, cell-surface or cell-membrane GO annotation were considered for the comparative analysis of the cell-surface glycoprotein ST vs. SF cells (i.e., 581 proteins, **Supplemental file 6**).

**Figure 9.**
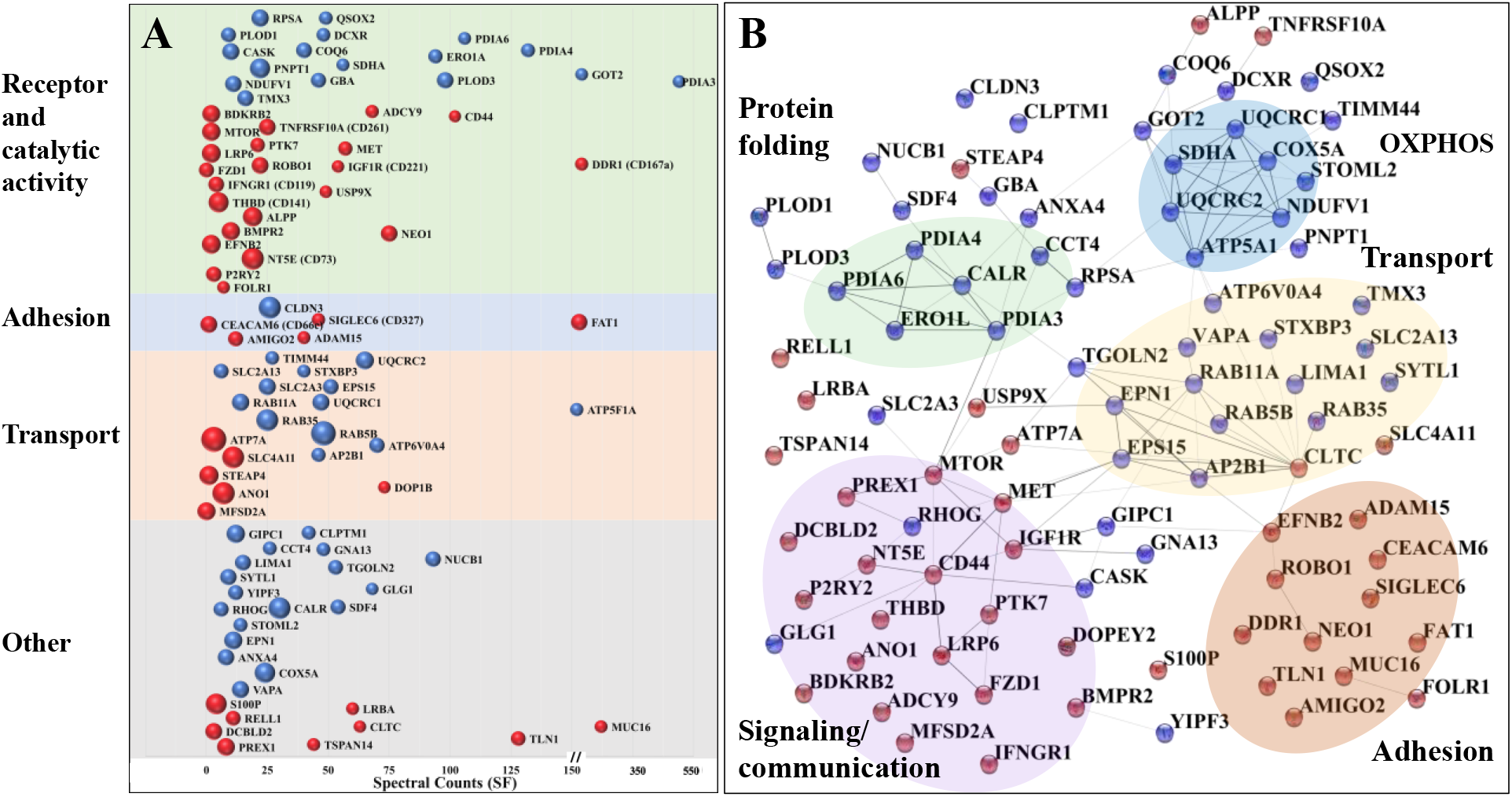
Proteins with changed abundance in the cell-membrane proteome. (**A**) Proteins with increased (red) and decreased (blue) abundance in ST vs. SF cells, categorized by function (y-axis) and spectral counts in SF cells (x-axis); the sphere size is proportional to the log2(fold change) in protein spectral counts. (**B**) Cytoscape visualization of the STRING PPI network created with the proteins that displayed a change in abundance (same color scheme as in A).

The biological processes that were up-regulated in serum-deprived cells were represented by proteins involved in (a) OXPHOS and mitochondrial ATP synthesis coupled electron transport, (b) ER protein folding, negative regulation of unfolded protein response (UPR) and Ca^2+^ homeostasis, and (c) intracellular molecular and vesicle-mediated transport, localization, secretion, and cellular homeostasis (**Figure 9B**). Intensified transport and localization processes were presumably provoked by adaptation and survival responses to serum-deprived stress. Vesicle trafficking and membrane fusion included Rab11-mediated endocytic recycling, clathrin mediated endocytosis and exocytosis, and appeared to target members of the ER protein folding machinery, Golgi apparatus and mitochondrial oxidative phosphorylation (OXPHOS). Calreticulin (CALR) and members of the protein disulphide isomerase (PDI) family are involved in maintaining cellular homeostasis by acting as chaperones that aid the folding of proteins destined for secretion in the ER. Their over expression has been observed in ER stress, but was also correlated with various cancerous cell states, while their translocation to the cell surface was associated with cancer progression and invasion (68–70). Metabolic rewiring of growing and proliferating cancer cells to use aerobic glycolysis for ATP production, instead of OXPHOS, is an established and extensively studied mechanism of energy production (the Warburg effect (71). It has been recognized, however, that different types of cancer cells can use both OXPHOS and glycolysis for ATP production (72), and, as a result, OXPHOS inhibitors have been suggested for targeting metabolic processes in high OXPHOS cancers (72). However, the behavior of cancer cells under nutrient-deprived conditions is not well understood, certain studies pointing toward reinforced OXPHOS activity in serum-deficient cells (73,74). In our study, the upregulation of a number of mitochondrial inner membrane respiratory chain proteins in the serum-deprived cells was also indicative of cells relying more heavily on OXPHOS for producing ATP (i.e., complex I-NDUFV1, complex II-SDHA, complex III-UQCRC1/UQCRC2, complex IV-COX5A, and complex V-ATP5A1). It was not clear, however, whether elevated OXPHOS and/or stress-induced protein re-localization to the plasma membrane was responsible for the increased abundance of the proteins in the cell-membrane fraction of the serum-starved cells. Nonetheless, the presence and activity of the ATPase complex components and of other mitochondrial matrix proteins and OXPHOS complexes in the cell-membrane and lipid rafts has been described before (75,76). The components of the F1F0ATP synthase complex on the surface of certain tumors has been associated with more aggressive, late stage metastatic cancers (77)-as was the case of the SKBR3 cells that were collected from a pleural effusion metastatic site. Cell-surface ATPase activity has been also associated with synthesis of extracellular ATP, binding of various ligands, and purinergic signaling (76). Extracellular ATP acts as an intercellular messenger that can interact with various cell-surface receptors, triggering, depending on conditions, cell death, proliferation, or various immune responses. When acting on P2RY receptors, such as the P2RY2 and P2RY6 detected on the surface of SKBR3 cells, ATP can activate ERK-MAPK, PI3K-AKT and survival pathways, or support EMT, invasiveness and metastatic spreading (78). ATP targeting in the tumor microenvironment has been attempted, therefore, as a cancer therapy in several clinical trials (78). In a similar manner, it has been suggested that OXPHOS complexes in the plasma membrane represent a source of extracellular superoxide which can exert various regulating roles in cellular function (76). It was interesting to note that unlike in the serum-deprived cells, upregulated HIF-1 signaling via IGF1R, MTOR, and IFNGR1 was observable in the serum-treated cells (**Figure 9**). In support of such findings, previous reports have revealed that metabolic reprogramming of cancer cells to use glucose metabolism for energy production uses mechanisms common to HIF-1 accelerated glucose metabolism (73,79).

As expected, the serum-stimulated cells displayed two major categories of up-regulated processes represented by proteins involved on one hand in cell communication and cell-surface receptor signaling (e.g., IGF1R, MTOR, FZD1, P2RY2, MET, IFNGR1, BMPR2, CD44, BDKRB2), and, on the other hand, in cell-cell adhesion and cell-matrix interactions, locomotion and migration (e.g., EFNB2, TLN1, ADAM15, MUC16, CD44, SIGLEC6, FAT1, ROBO1, DDR1, LRP6). MTOR signaling (MTOR, IGF1R, FZD1, LRP6), which plays a central role in regulating cell growth and anabolic/catabolic metabolism, appeared to be activated by the presence of growth factors, hormones and nutrients from serum (80). MTOR hyperactivation is integral to several oncogenic pathways (e.g., PI3K/AKT and MAPK) (80), and, similarly, IGF1R overexpression and signaling has been shown to be implicated in the regulation of survival and proliferation of many cancers (81). Therefore, co-targeting PI3K, mTOR, and IGF1R proved to be effective in reducing tumor growth and decreasing cell migration and invasion (82). Targeting of the mesenchymal epithelial transition (MET) receptor Tyr kinase that is coded by the proto-oncogene *MET*, has also gained momentum, as its aberrant activation has been associated with a number of signaling pathways that promote cell survival, growth, proliferation, morphogenetic effects, and migration (e.g., PI3K/AKT, Ras/MAPK, JAK/STAT, SRC, Wnt/β-catenin) (83). Relevant to note is that MTOR, IGF1R and MET have been implicated in the activation of alternative pathways that drive resistance to therapeutic treatment with EGFR tyrosine kinase inhibitors (TKIs), resulting frequently in the recurrence of tumors (84). Processes related to cell differentiation and immune system responses were also elicited in the stimulated cells by many of the same proteins. In addition, CD44, an adhesion receptor expressed on the surface of many cancer cells that mediates cell-cell interactions and cell migration, has been recognized for its multifunctional roles in survival, angiogenesis, metastasis and activation of immune responses and inflammation (21,85). Altogether, in the presence of nutrients, these proteins were representative of key biological processes that support cancer cell proliferation, adhesion, migration and propensity for tissue invasion and metastasis.

## Conclusions

The metastatic SKBR3 cell-membrane proteome revealed a broad and rich map of receptors, immune response, adhesion and transporter proteins that sustain cancer-cell interactions with the native or drug-altered tumor microenvironment. As evidenced by PPI networks, the concerted action of cell-membrane proteins exposed synergistic capabilities that nourish aberrant cell proliferation and metastatic potential not just through well-known signaling mechanisms, but through all functional roles of the surfaceome. Various aspects of cell growth, proliferation and differentiation were mainly orchestrated by proteins with catalytic activity, kinase receptors, plexins, and some CDs via growth-factor initiated signaling pathways. Several important groups of proteins with newly identified activities in cancer development included the families of metalloproteinases, nectins, ephrins, and bone morphogenetic proteins. Alterations in the abundance of certain cell-membrane proteins in response to serum withdrawal provided novel insights into how cancer cells exploit metabolic mechanisms of energy production to sustain their proliferation. Cell migration, invasion and metastatic propensity were facilitated by proteins with roles in sustaining or regulating angiogenesis, cell-cell or cell-ECM interactions and EMT processes. Essentially, all cell-membrane protein categories, not just the CDs, contributed to mounting innate/adaptive or inflammatory immune responses, with ephrin and plexin receptors supporting chemokine signaling and chemotactic processes. The presence of receptor ligands with T-cell inhibitory functions, such as PDL1, pointed to abilities to stage immune escape. Propensity for altered drug uptake mechanisms and development of multi-drug resistance, mediated by solute carriers or ABC transporters, respectively, were also evident. The availability of a vast range of multi-functional cell-membrane proteins underscored, however, encouraging prospects for the development of more effective combination therapies that co-target proliferative, autocrine/survival, apoptotic, angiogenesis, and cell migratory pathways, as well as more subtle cancer checkpoint immunotherapies. Reassuring was the presence of a large number of immunological markers that reflected yet unexplored opportunities for cancer diagnosis, prognosis and assessment of recurrence after therapy.

## Acknowledgment

This work was supported by an award from the National Institute of General Medical Sciences (Grant No. 1R01GM121920) to IML.

## Author contributions

AK performed the experiments and processed the data; IML and AK analyzed the data and wrote the manuscript; IML coordinated the work. Both authors reviewed and approved the final version of the manuscript.

## Materials & Correspondence

Address correspondence and material requests to Dr. Iulia M. Lazar.

## Data availability

The mass spectrometry raw files were deposited to the ProteomeXchange Consortium via the PRIDE partner repository with the following dataset identifiers: PXD028976, PXD028977, and PXD028978.

