## Supplemental File 4 for "The SKBR3 Cell-Membrane Proteome: Role in Aberrant Cancer Cell Proliferation and Resource for Precision Medicine Applications"

### Supplemental File 5
### GPCR/PRM validations

### Adhesion G protein-coupled receptor L1 (ADGRL1)

GSIYAGDVSSSVK, Charge 2, m/z = 635.3193

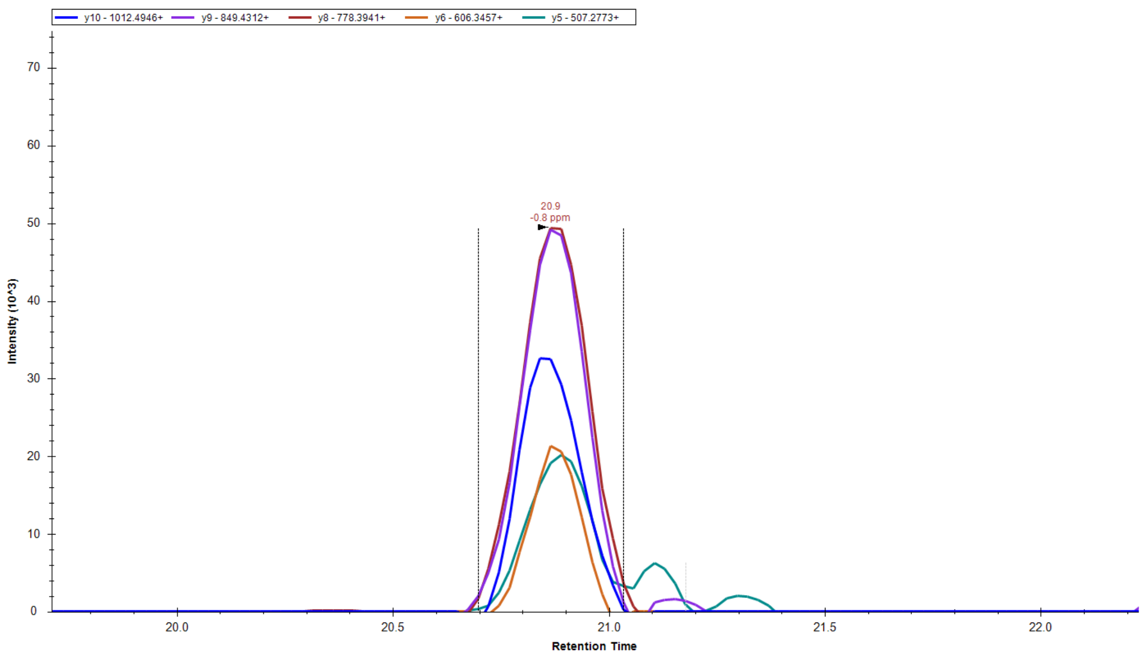


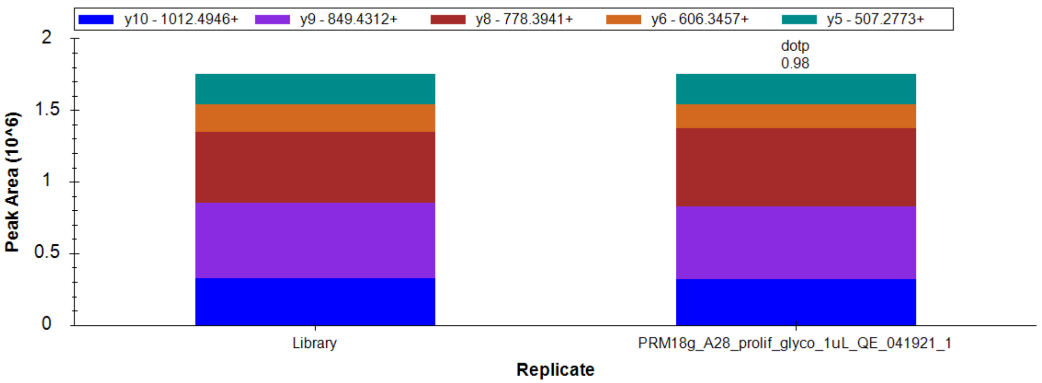


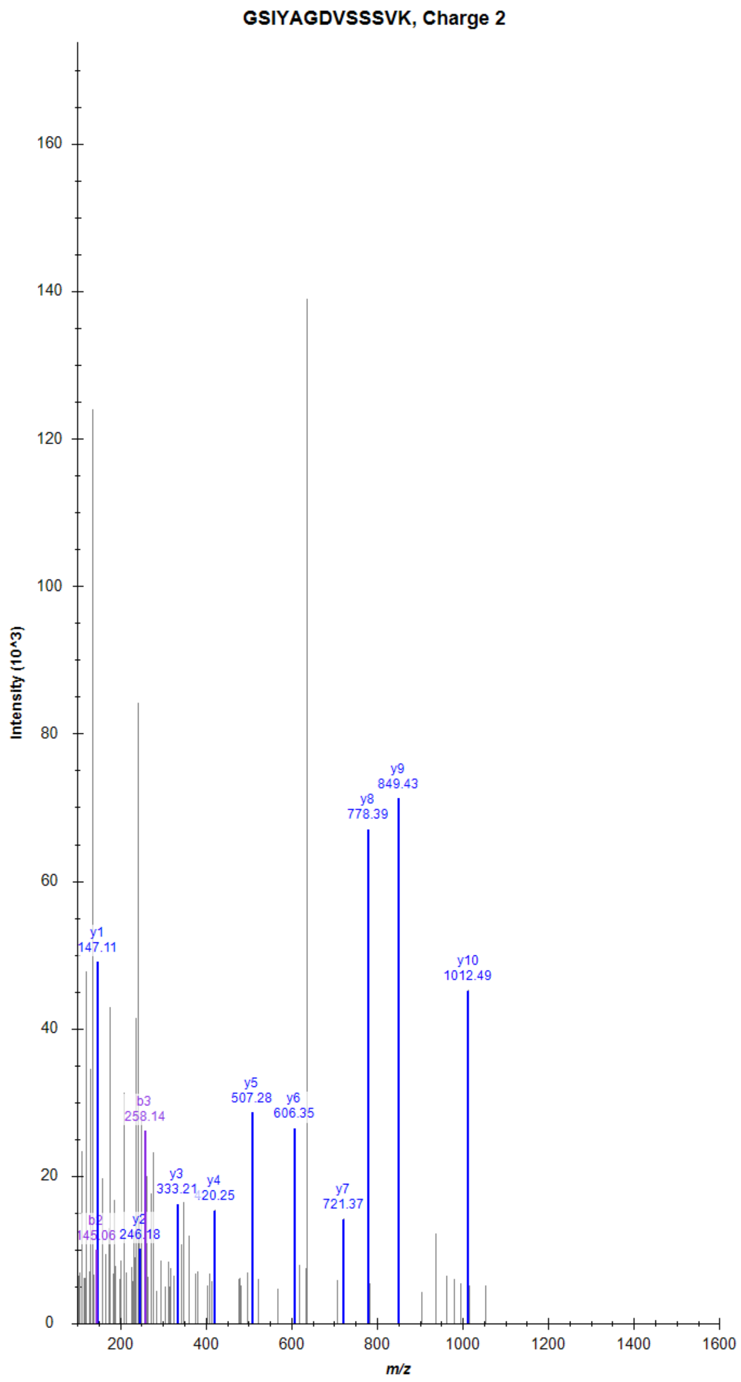

### Adhesion G protein-coupled receptor L1 (ADGRL1)

SGETVINTANYHDTSPYR, Charge 3, m/z = 675.6484

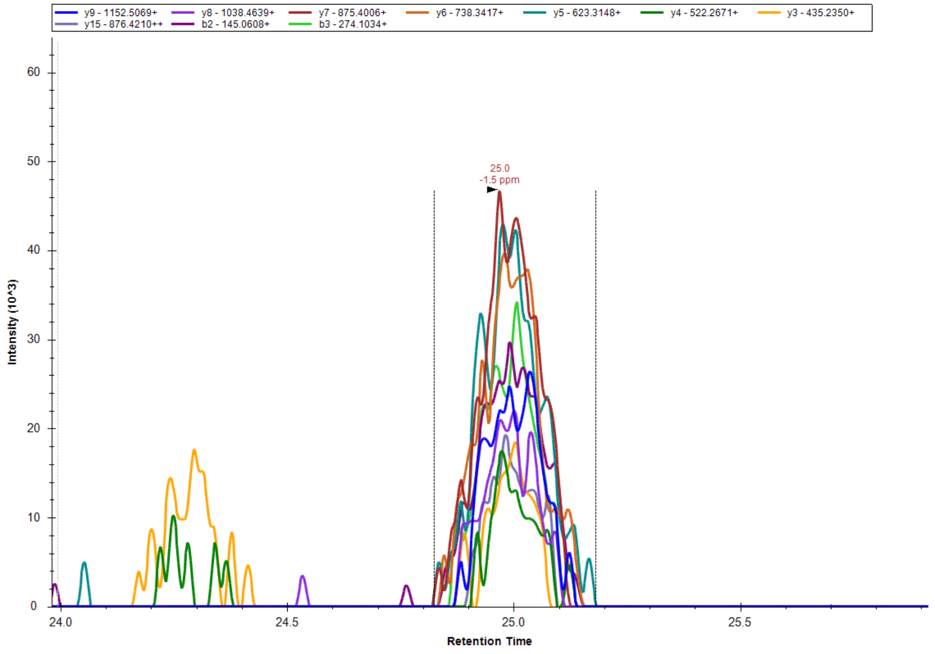


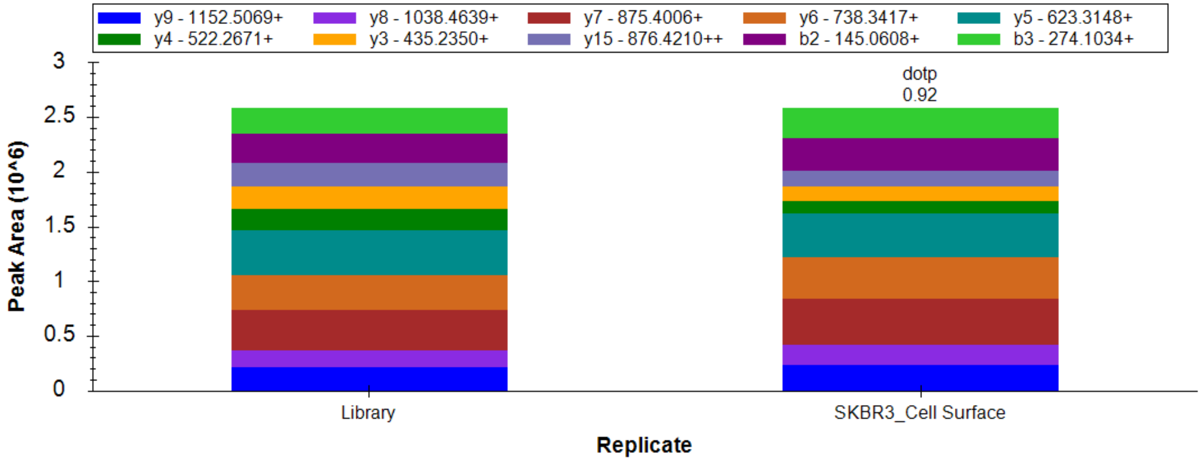


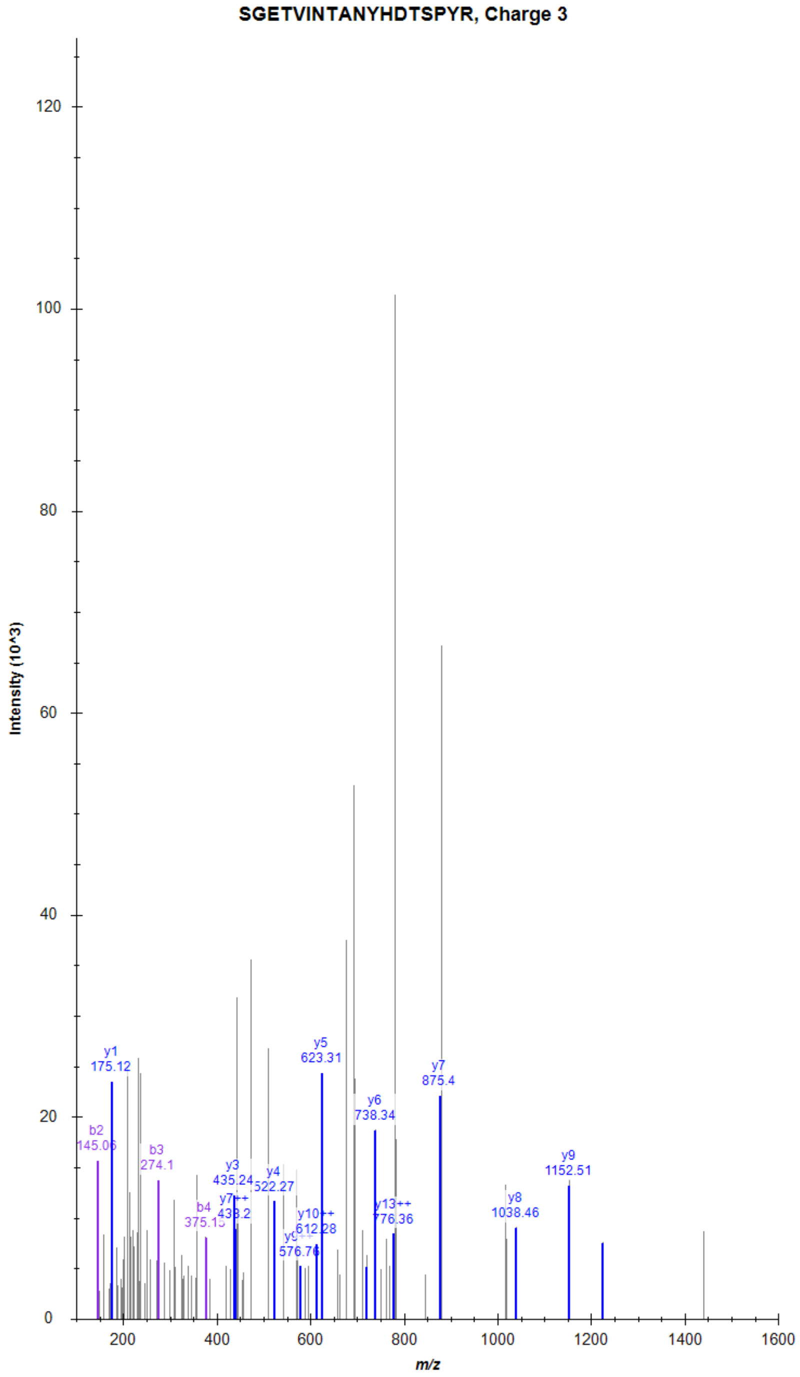

### Adhesion G protein-coupled receptor L2 (ADGRL2)

AALPFGLVR, Charge 2, m/z = 472.2903

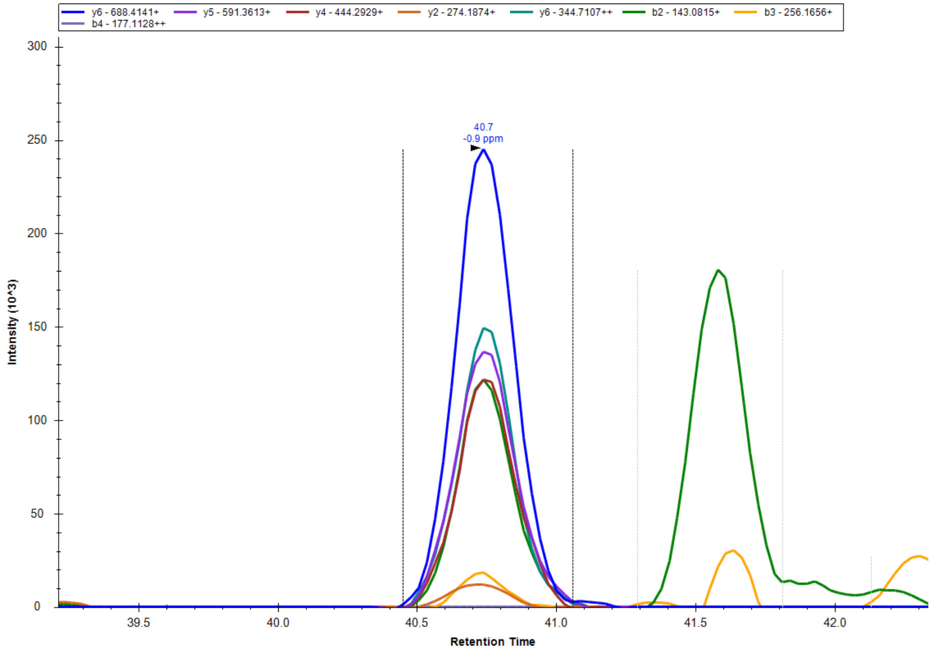


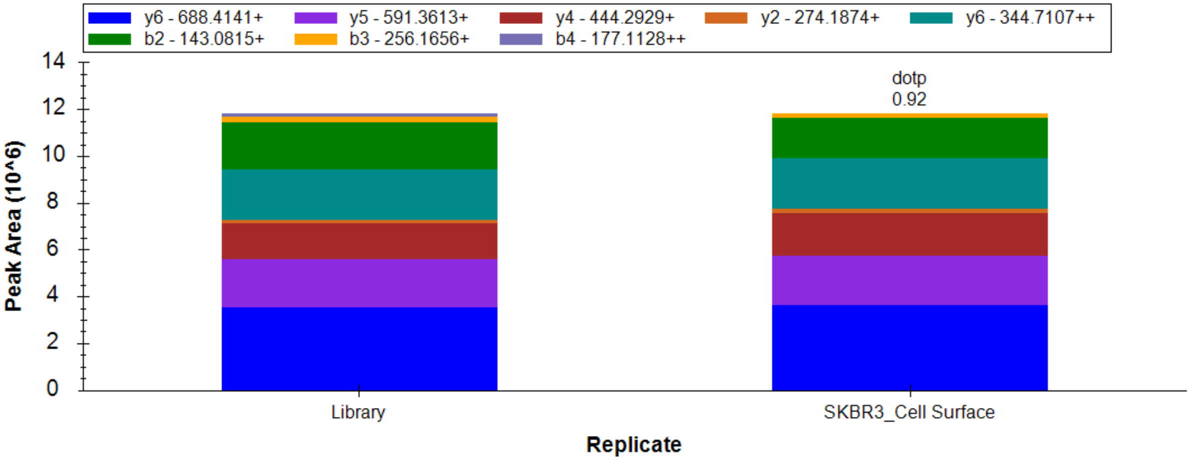


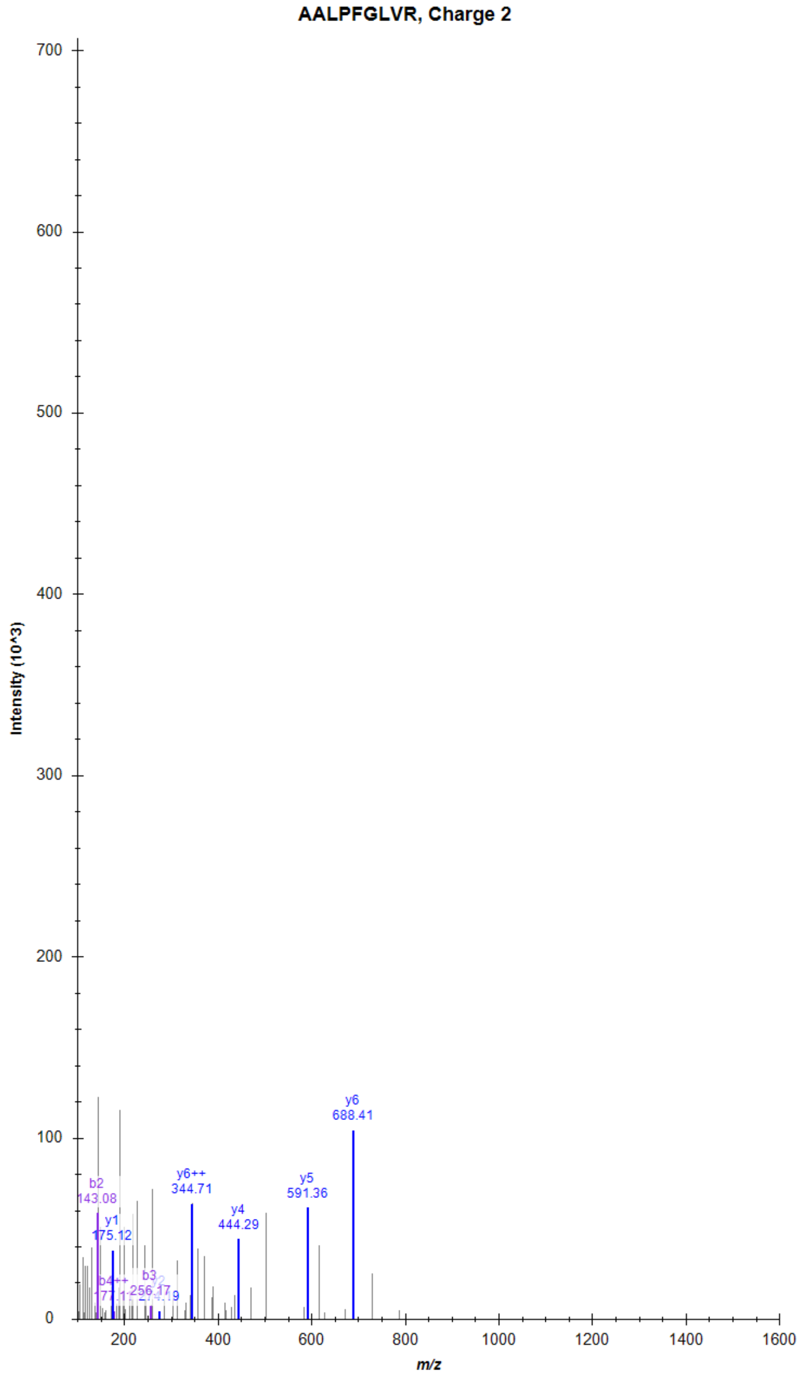

### Adhesion G protein-coupled receptor L2 (ADGRL2)

SGEAIINYANYHDTSPYR, Charge 3, m/z = 690.9888

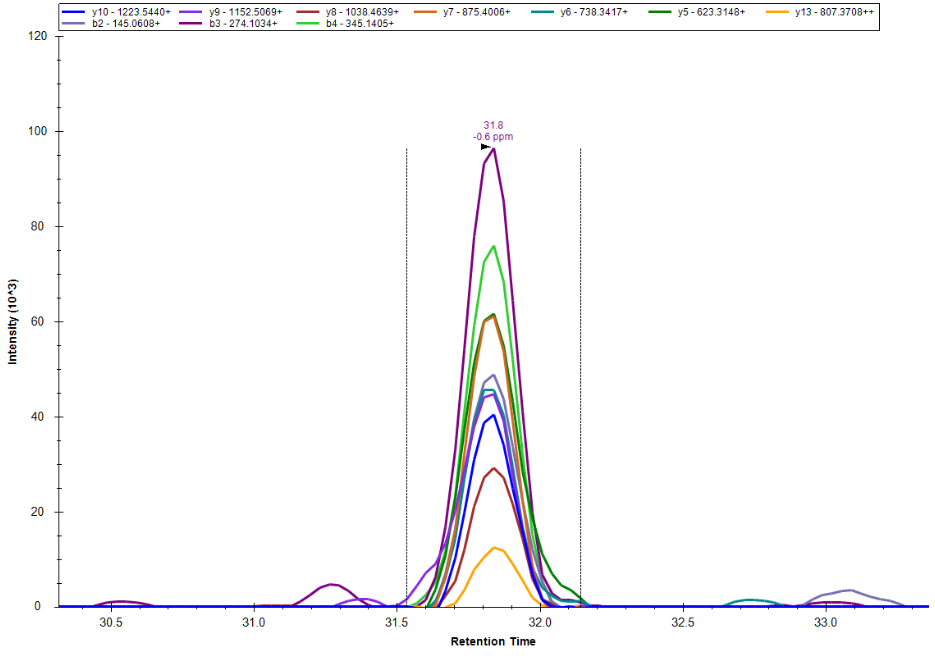


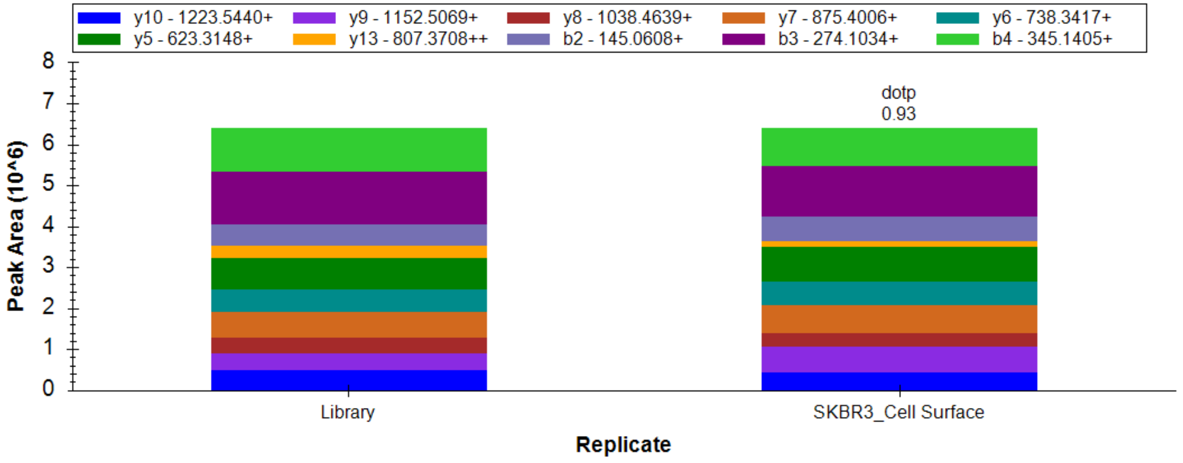


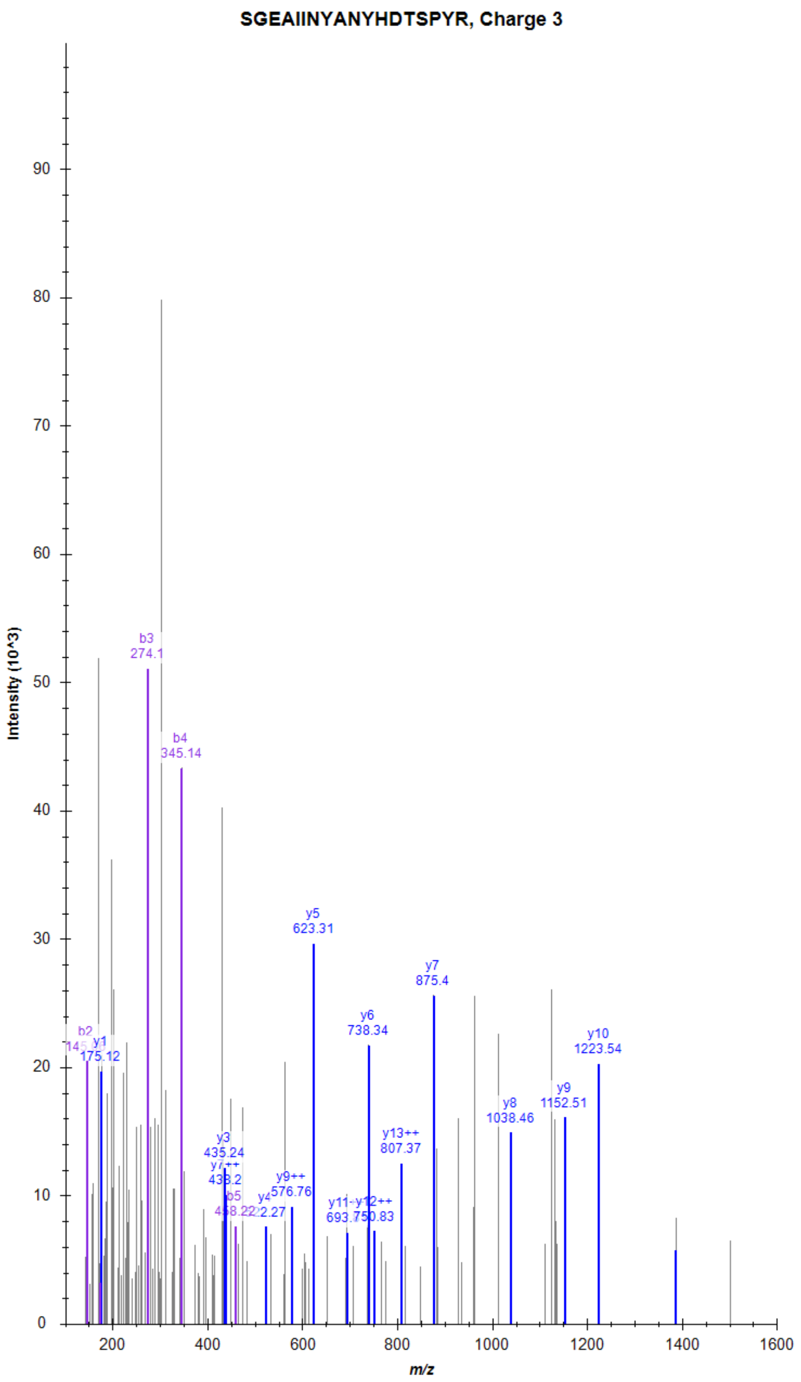

### Vasoactive intestinal polypeptide receptor 1 (VIPR1)

AASLDEQQTMFYGSVK, Charge 2, m/z = 887.9209

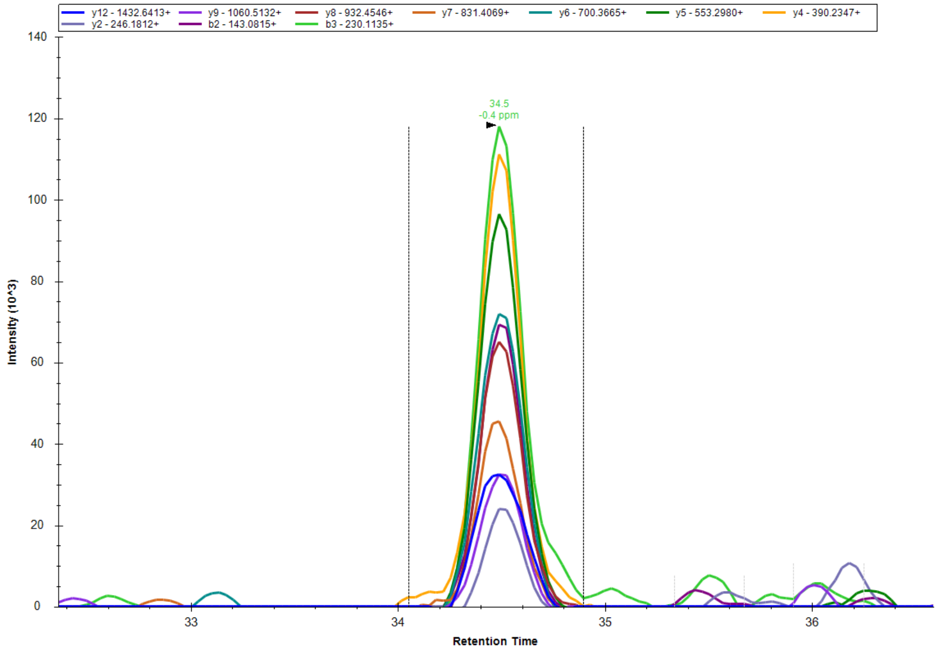


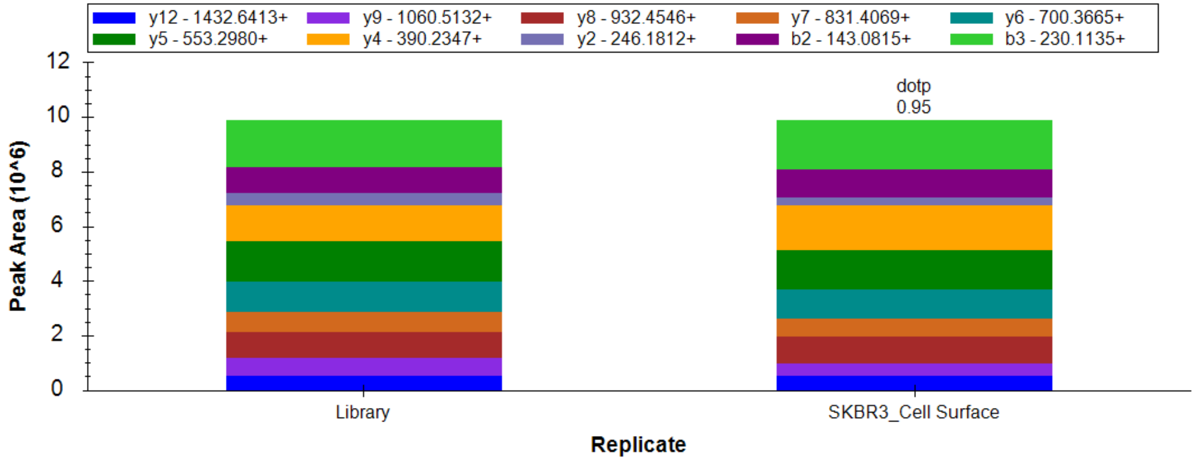


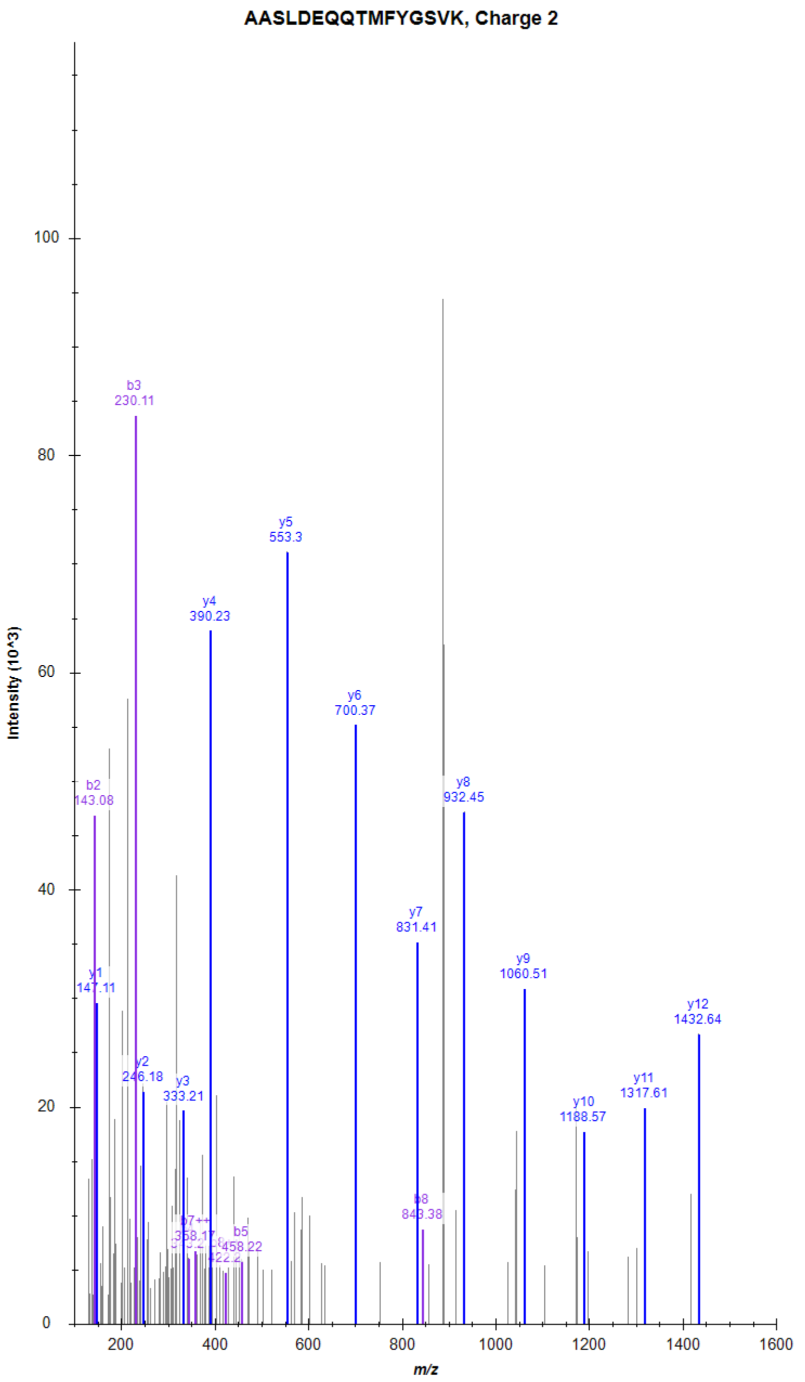

### Vasoactive intestinal polypeptide receptor 1 (VIPR1)

LFSSIQGR, Charge 2, m/z = 454.2533

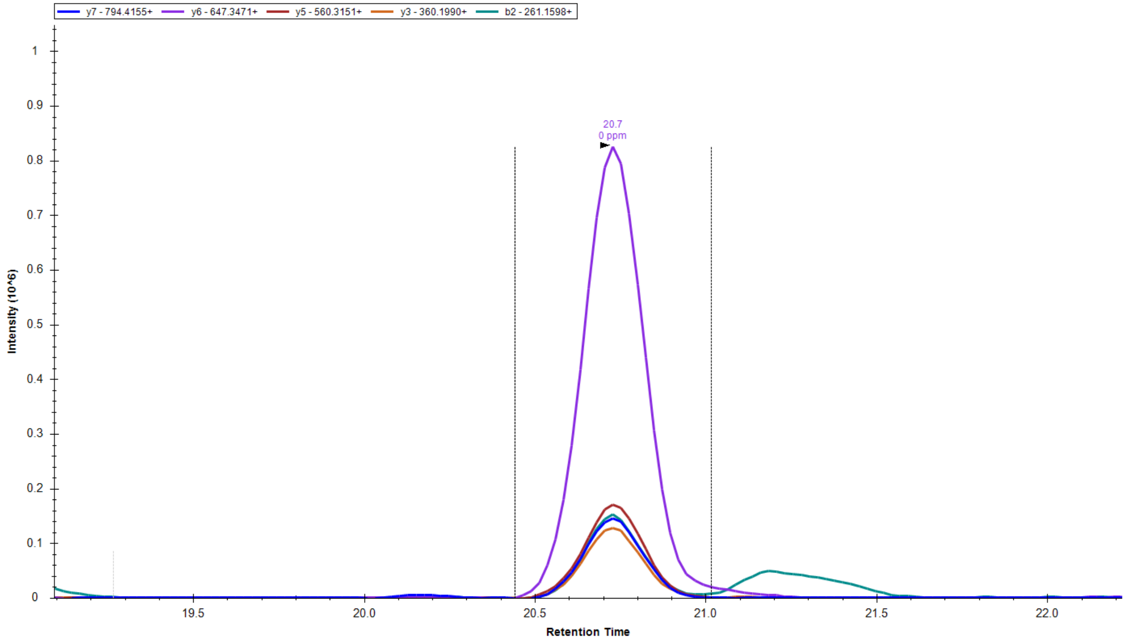


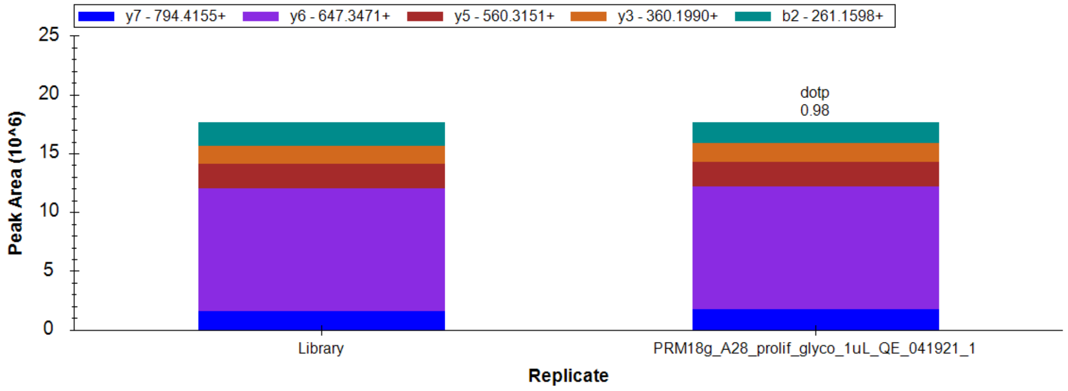


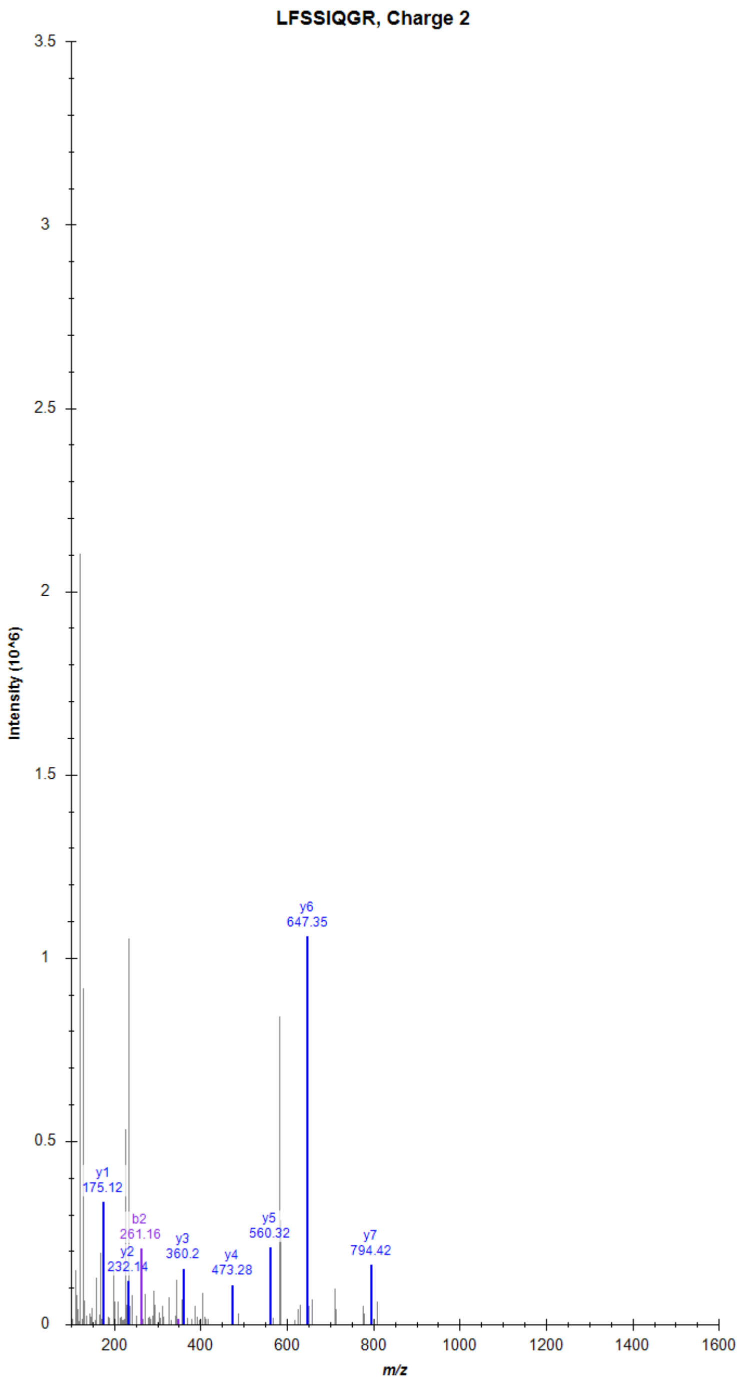

### CD97 antigen (CD97)

TSSAEVTIQNVIK, Charge 2, m/z = 695.3828

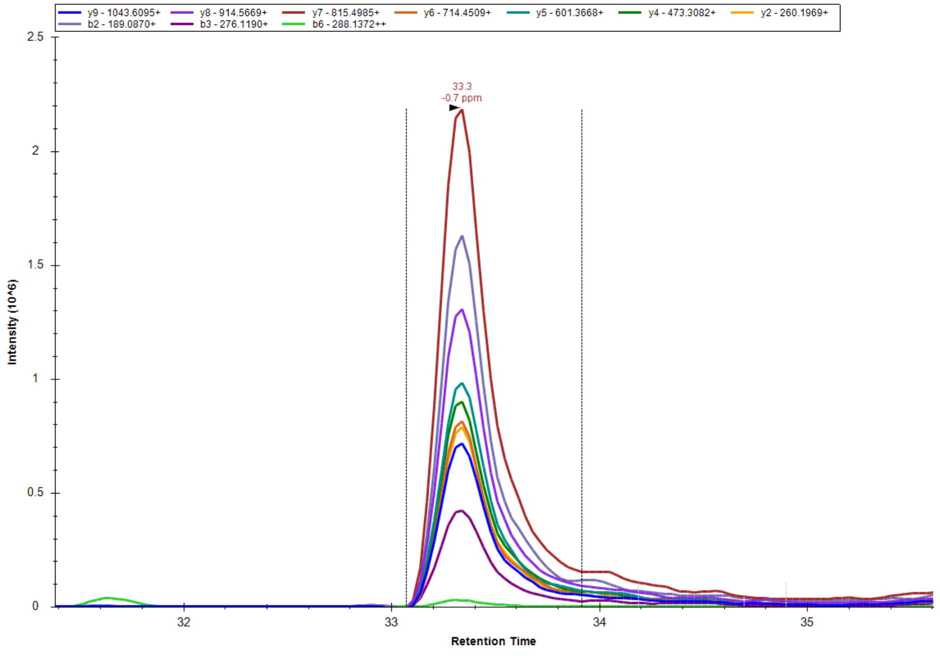


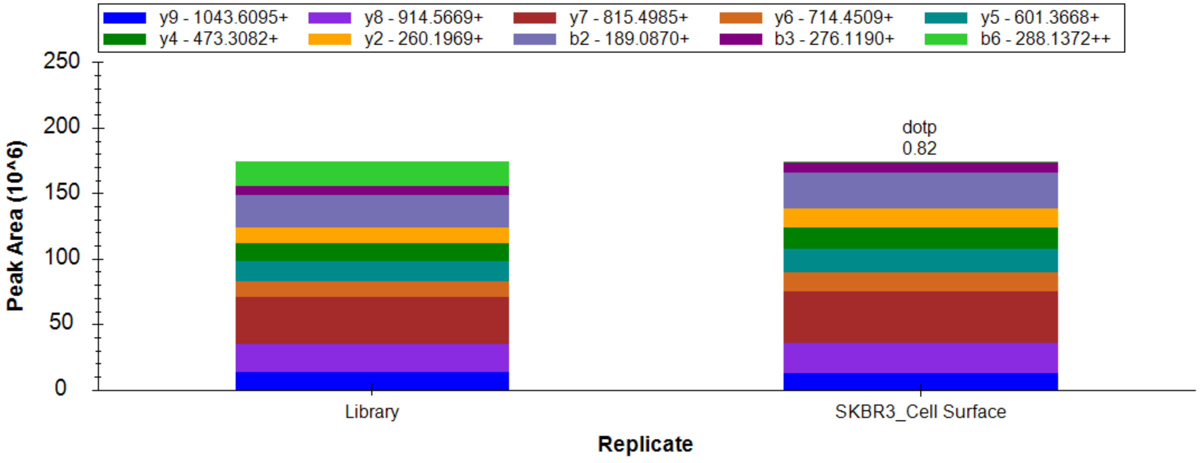


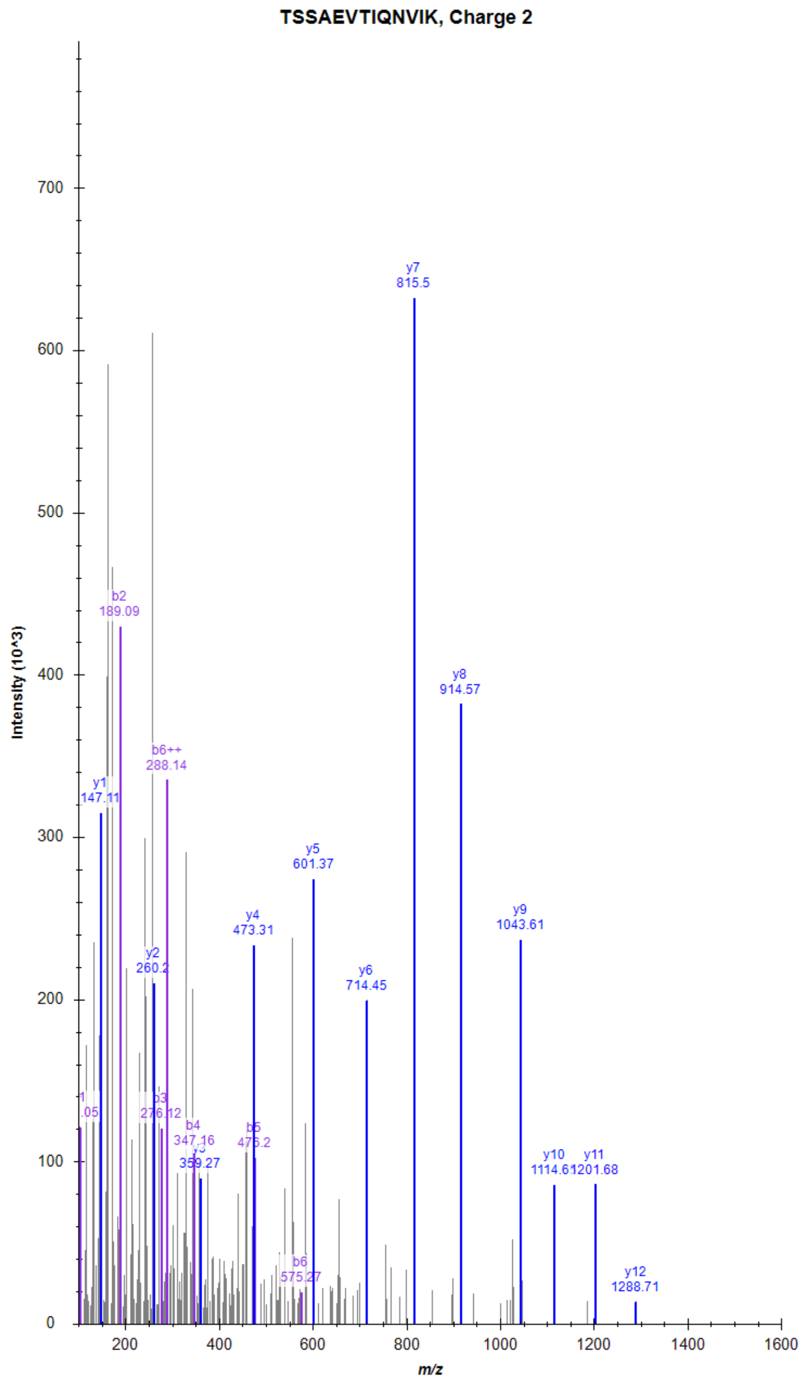

### CD97 antigen (CD97)

KQAELEEIYESSIR, Charge 2, m/z = 847.9334

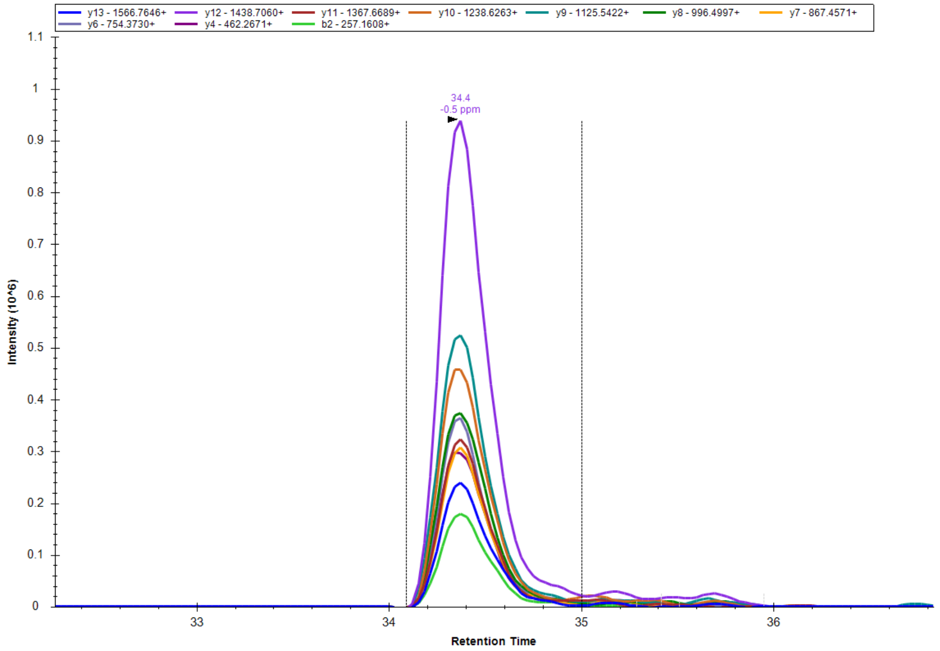


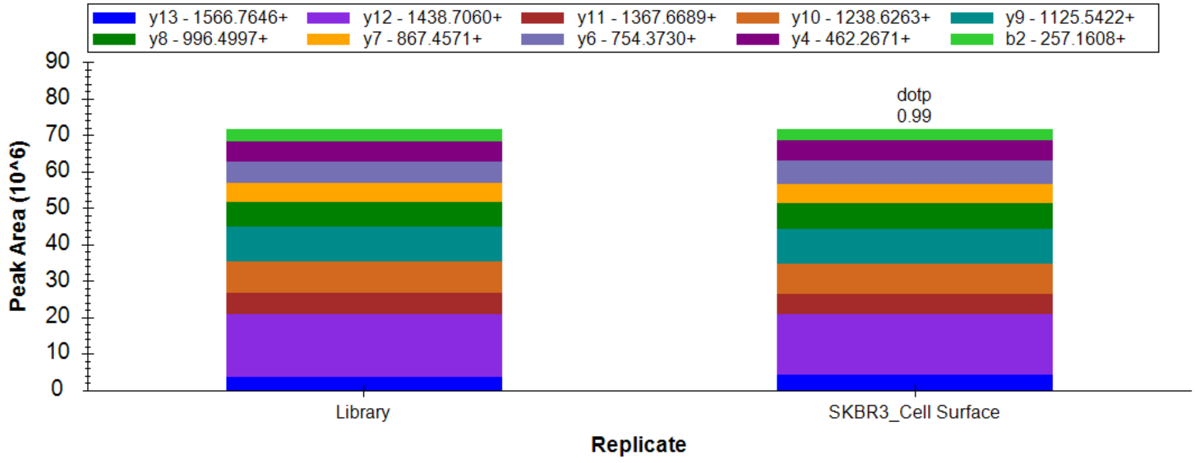


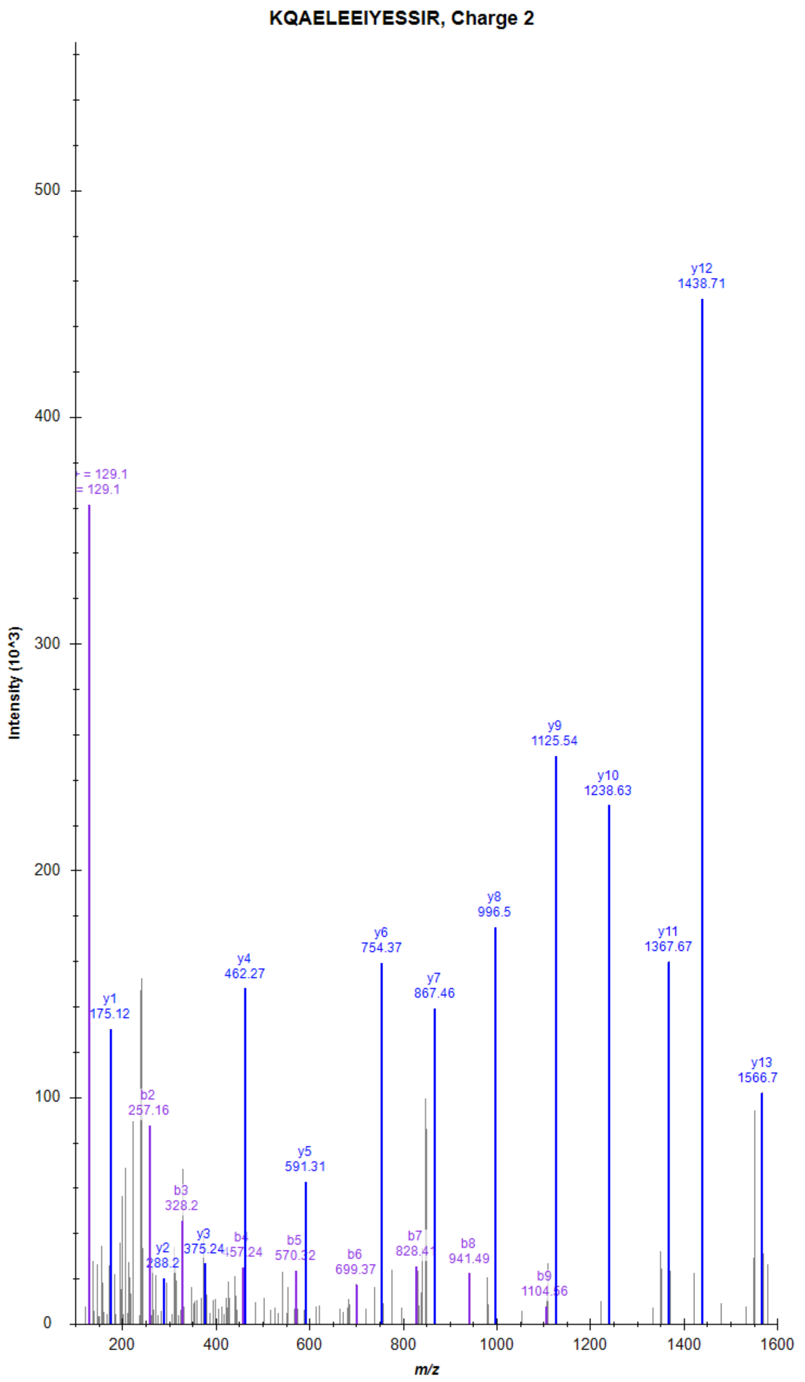

### Adhesion G-protein coupled receptor G6 (ADGRG6)

TGLFQDVGPQR, Charge 2, m/z = 609.3173

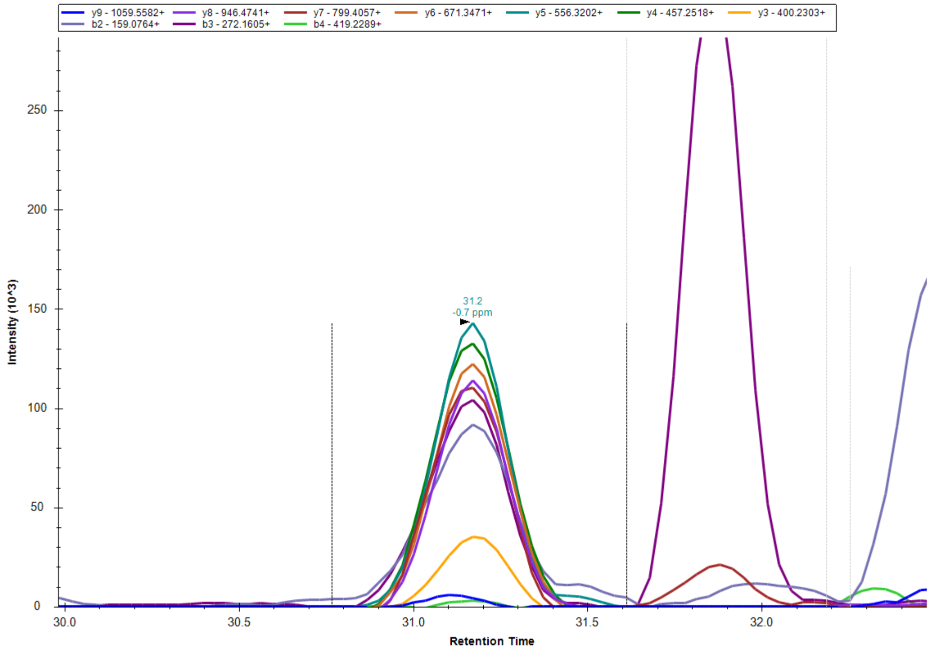


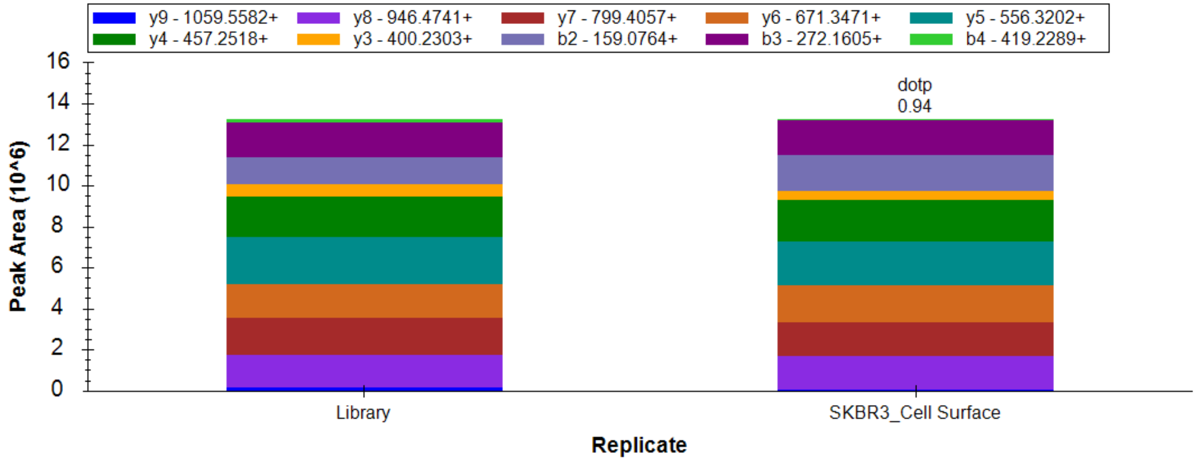


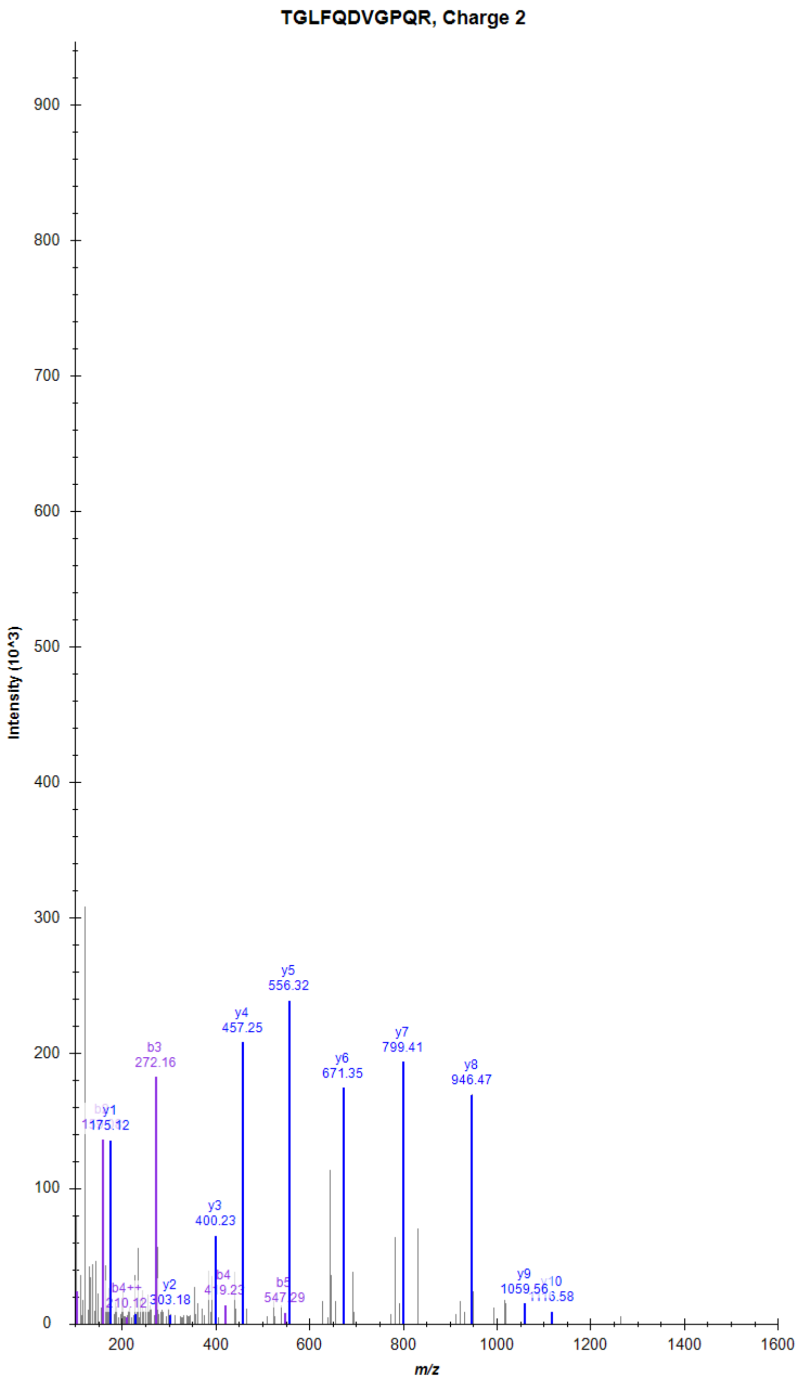

### Adhesion G-protein coupled receptor G6 (ADGRG6)

VILPQTSDAYQVSVAK, Charge 2, m/z = 859.9717

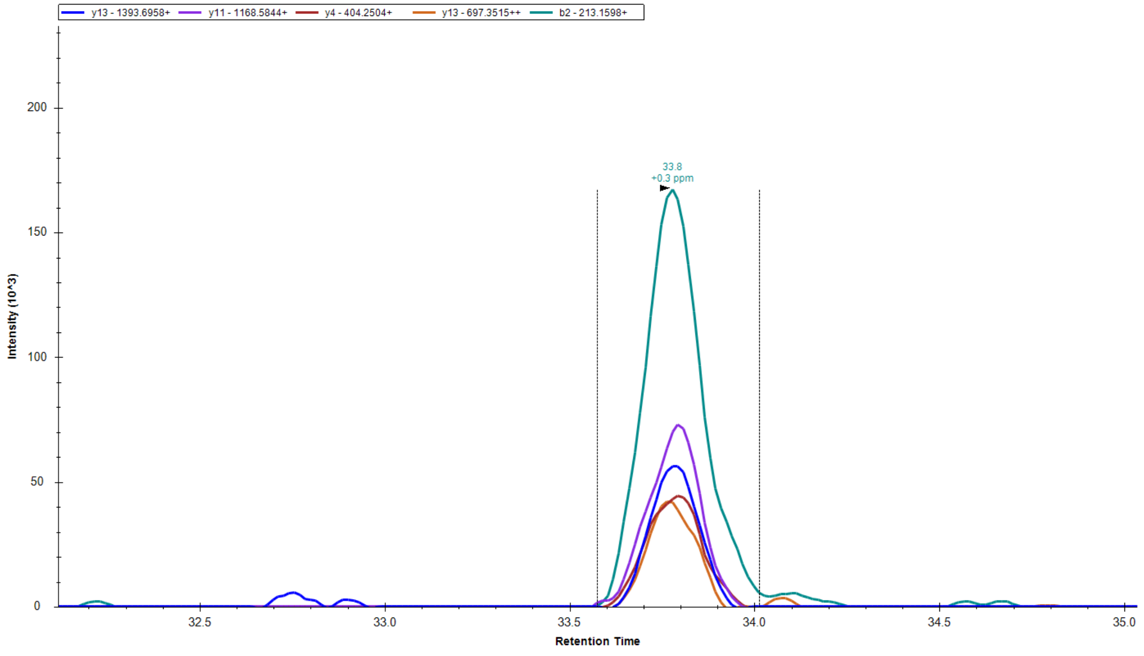


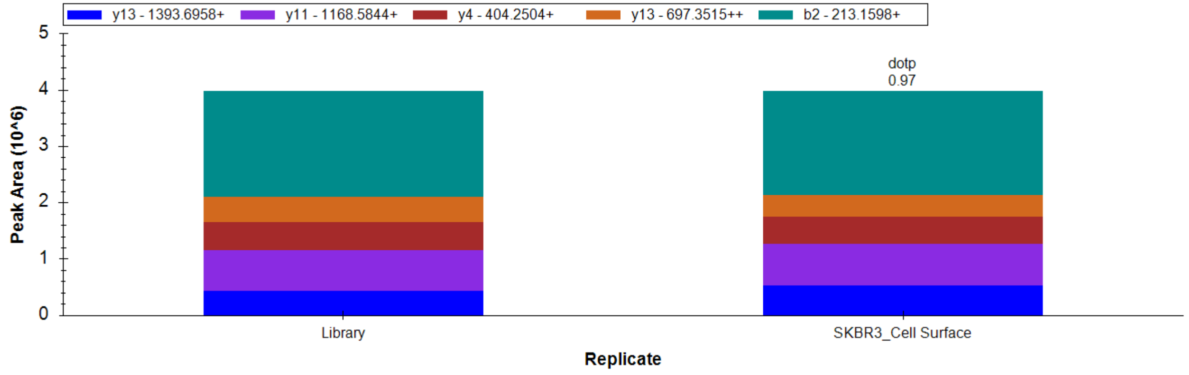


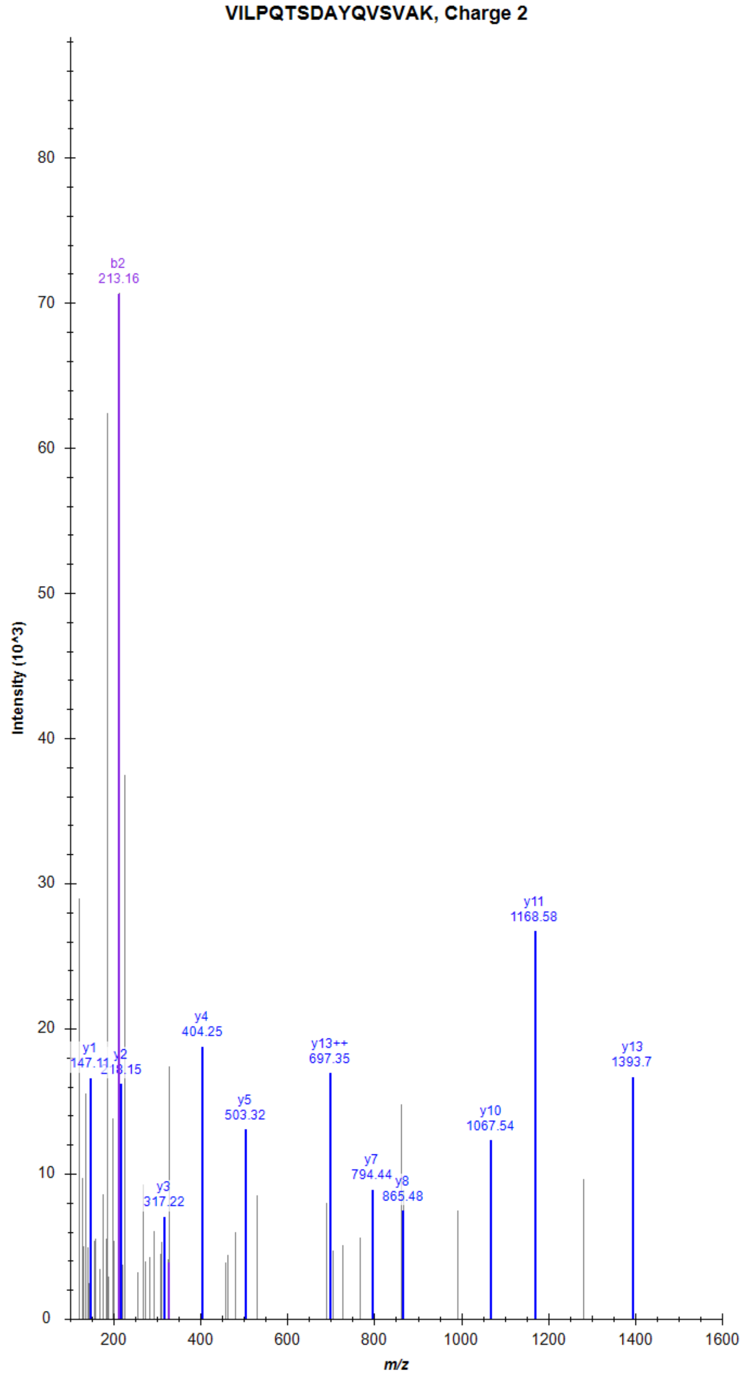

### Leucine-rich repeat-containing G-protein coupled receptor 4 (LGR4)

TLDLSYNNIR, Charge 2, m/z = 604.8171

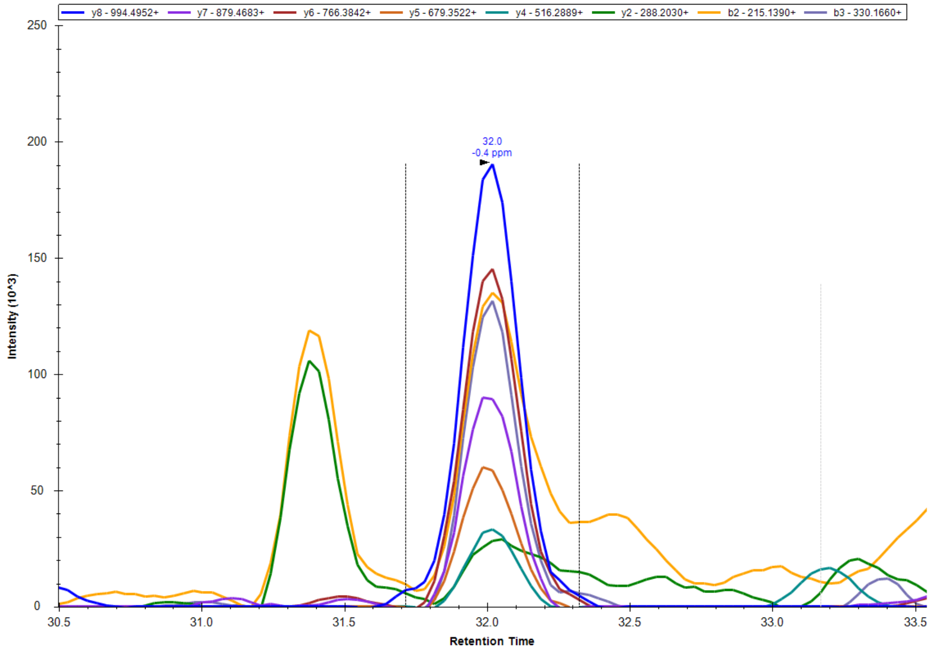


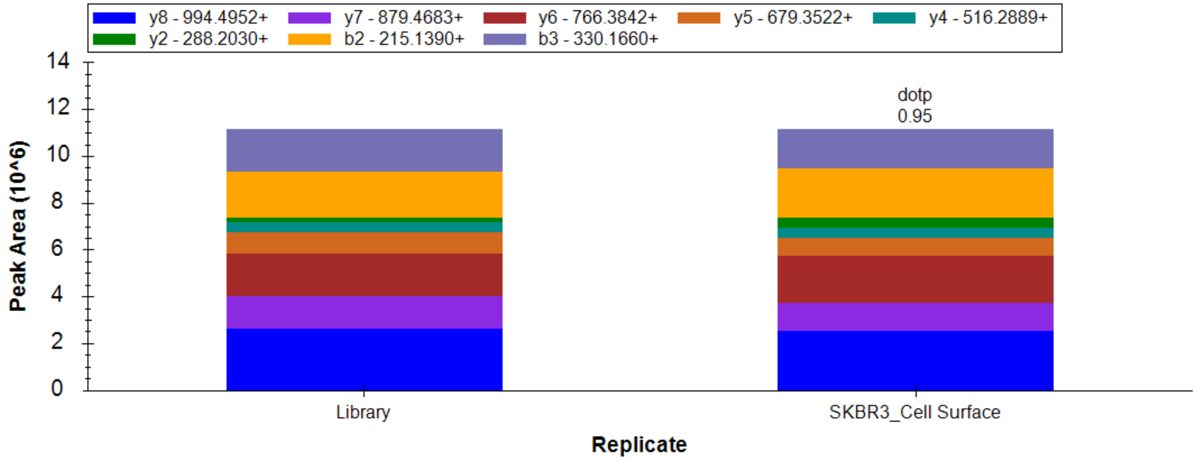


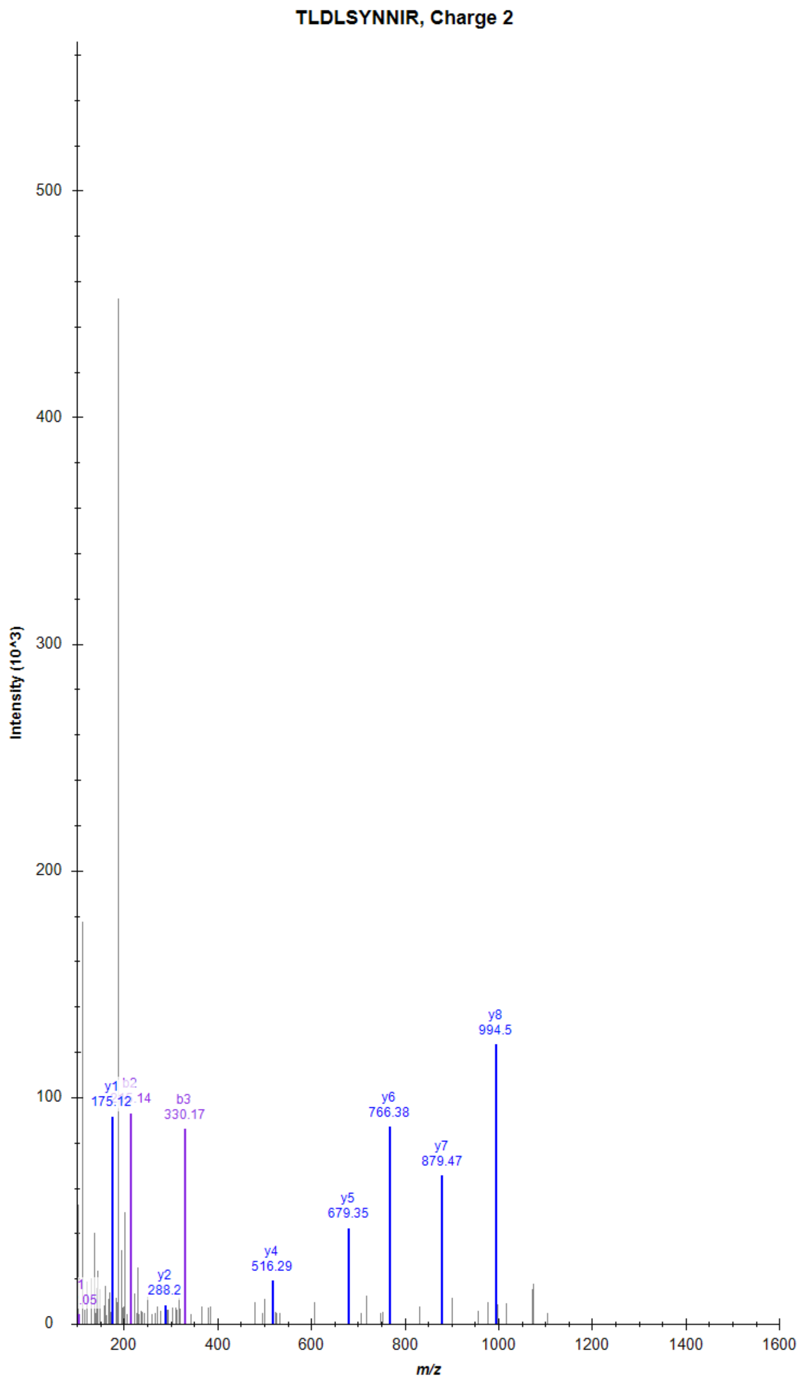

### Leucine-rich repeat-containing G-protein coupled receptor 4 (LGR4)

VLTLQNNQLK, Charge 2, m/z = 585.8464

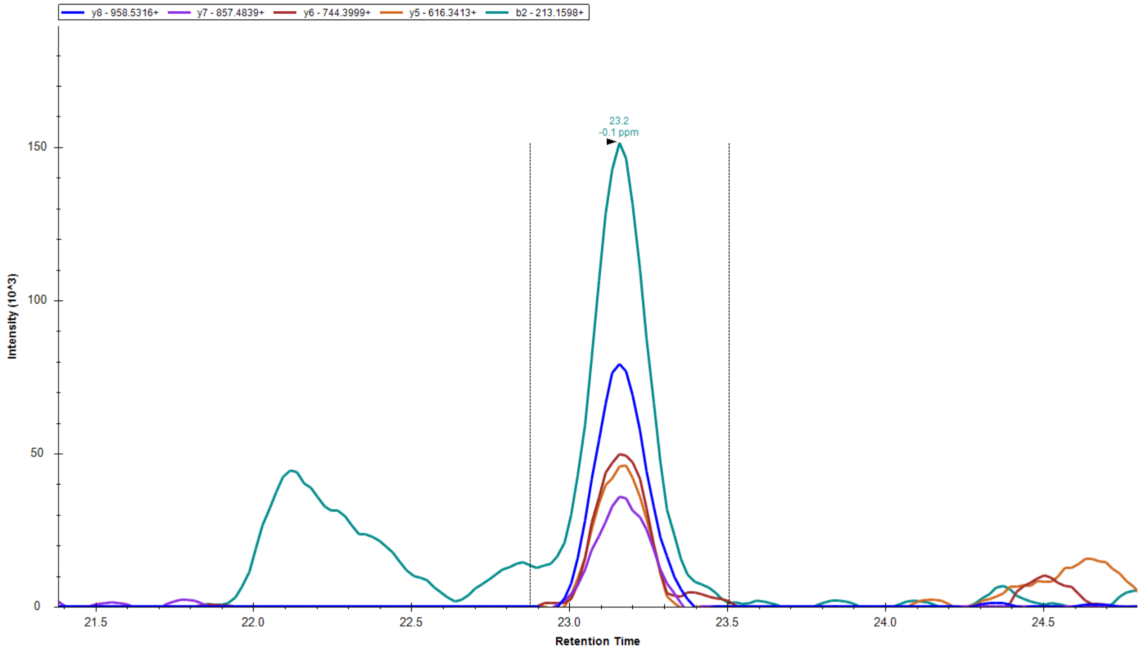


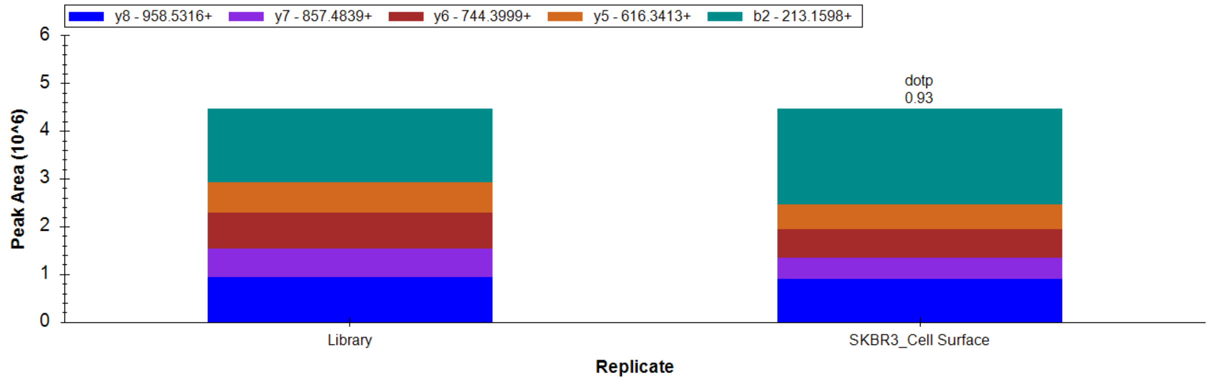


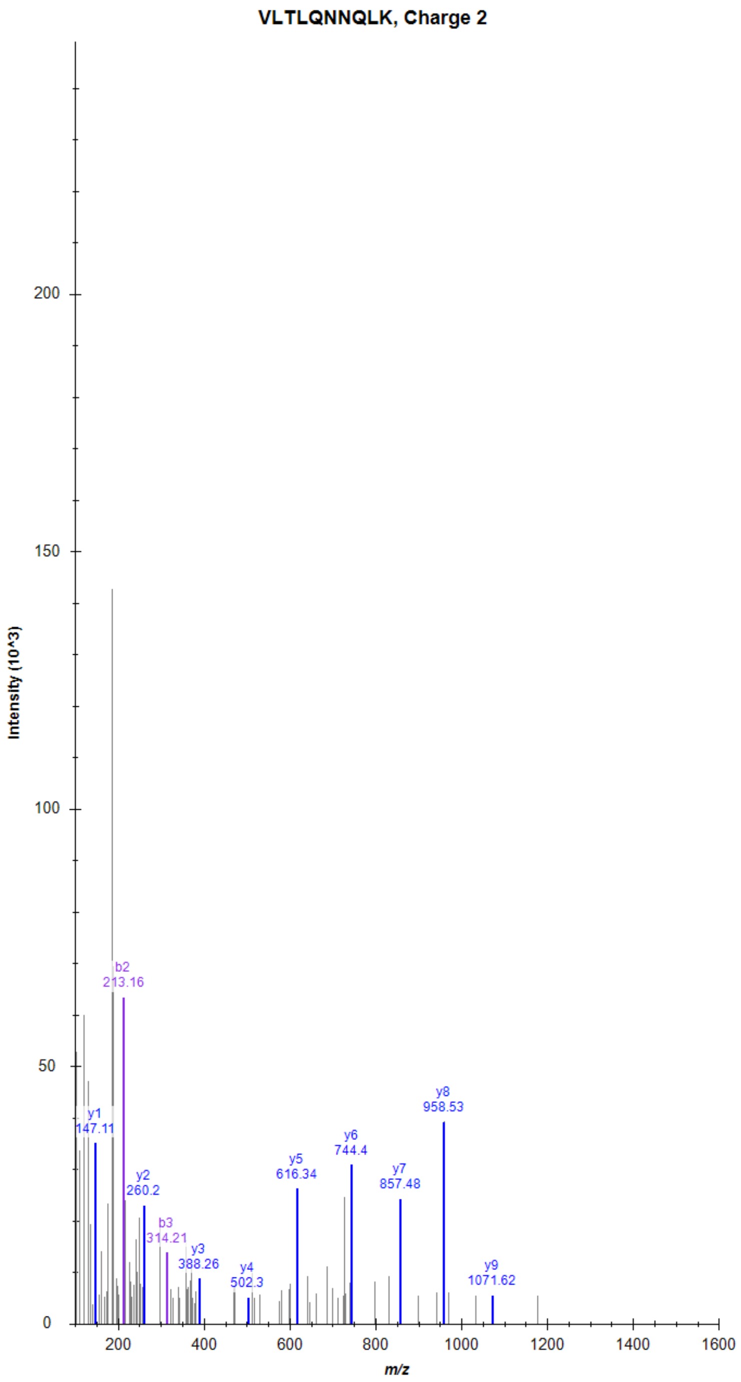

### Cadherin EGF LAG seven-pass G-type receptor 1 (CELSR1)

LVDTASTFLGGGSAGPK, Charge 2, m/z = 789.4125

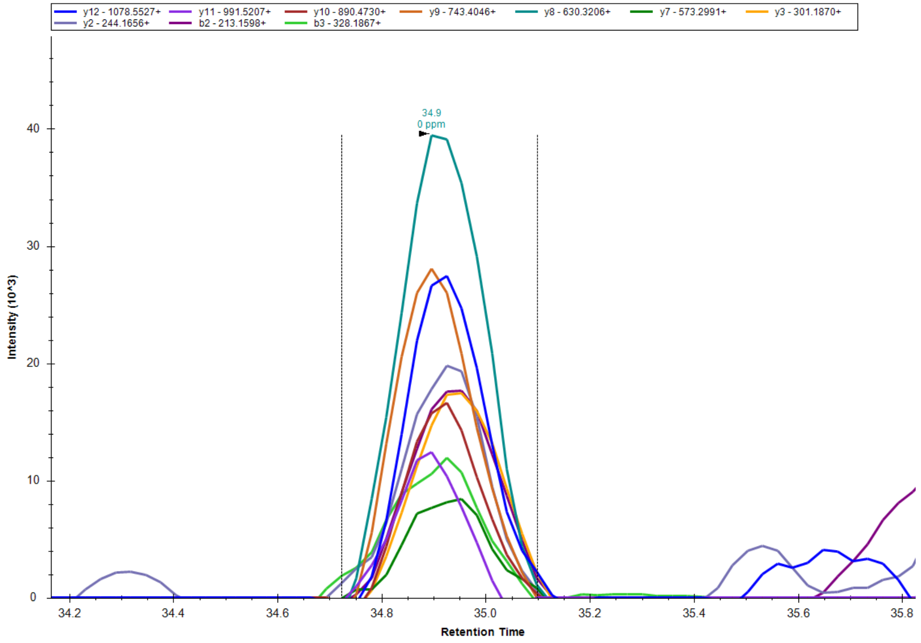


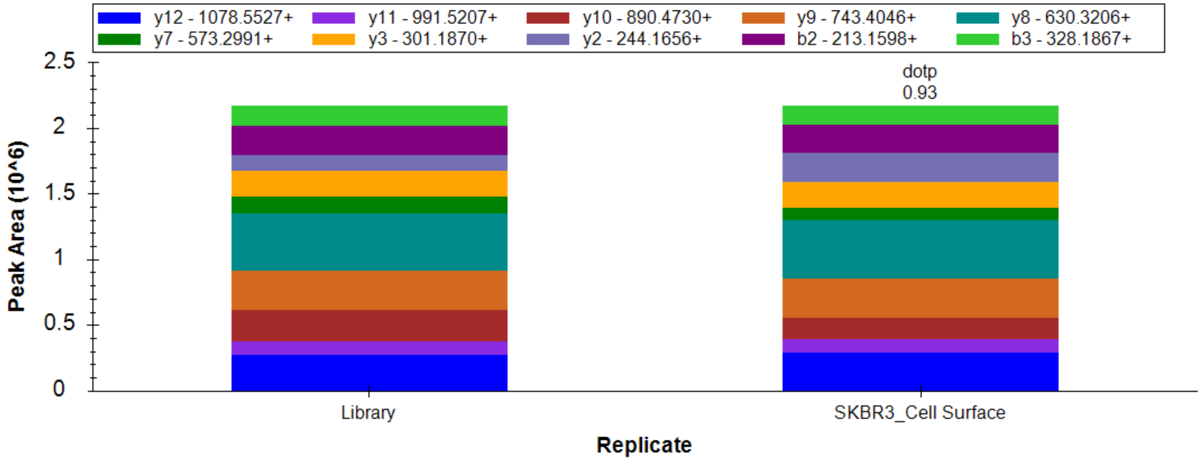


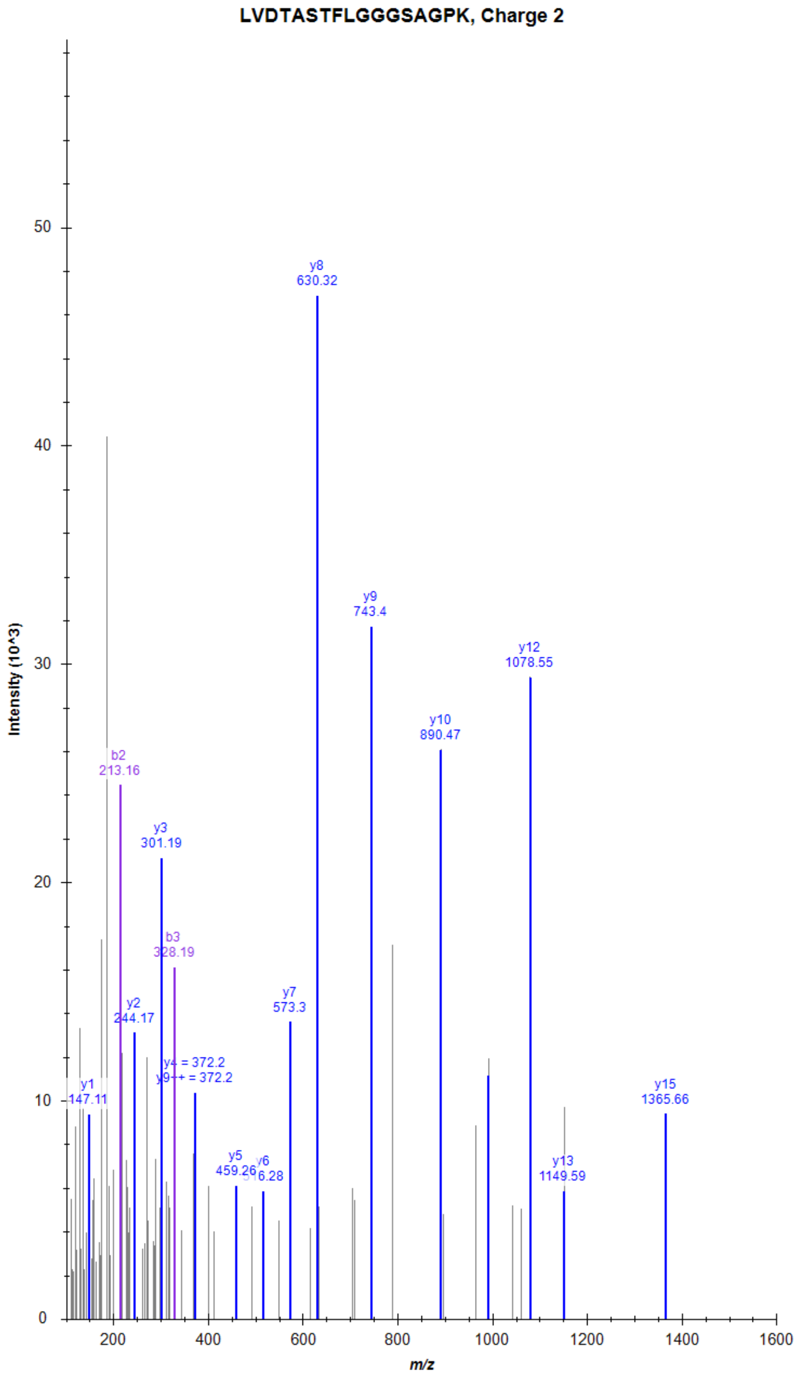

### Cadherin EGF LAG seven-pass G-type receptor 1 (CELSR1)

LLLLDPATGELQLSR, Charge 2, m/z = 819.9744

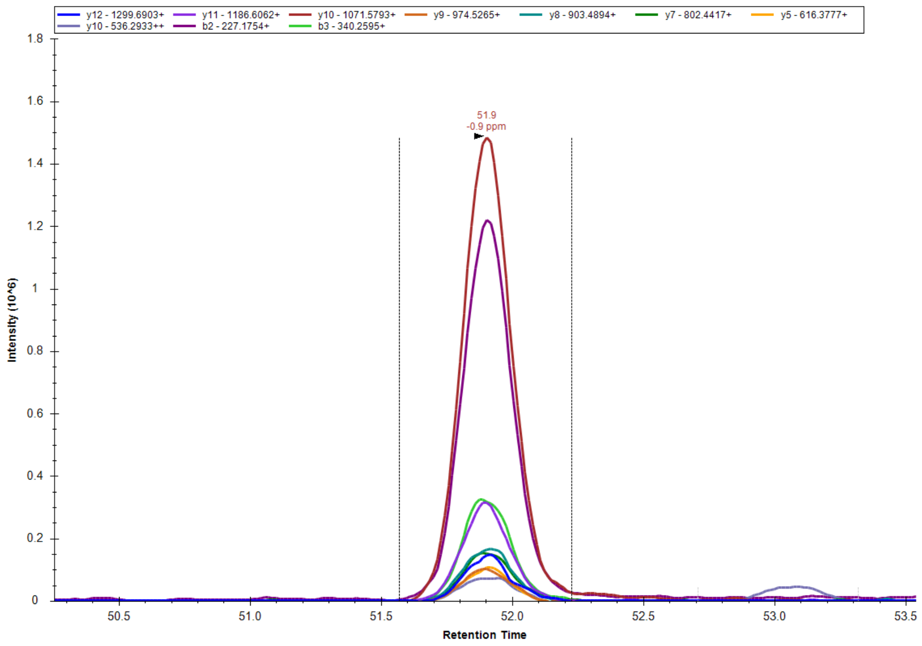


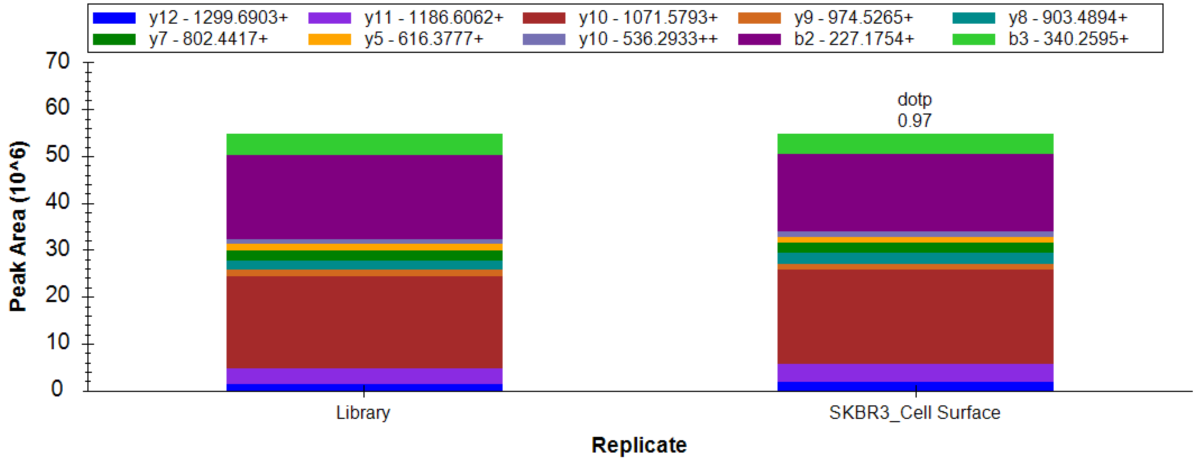


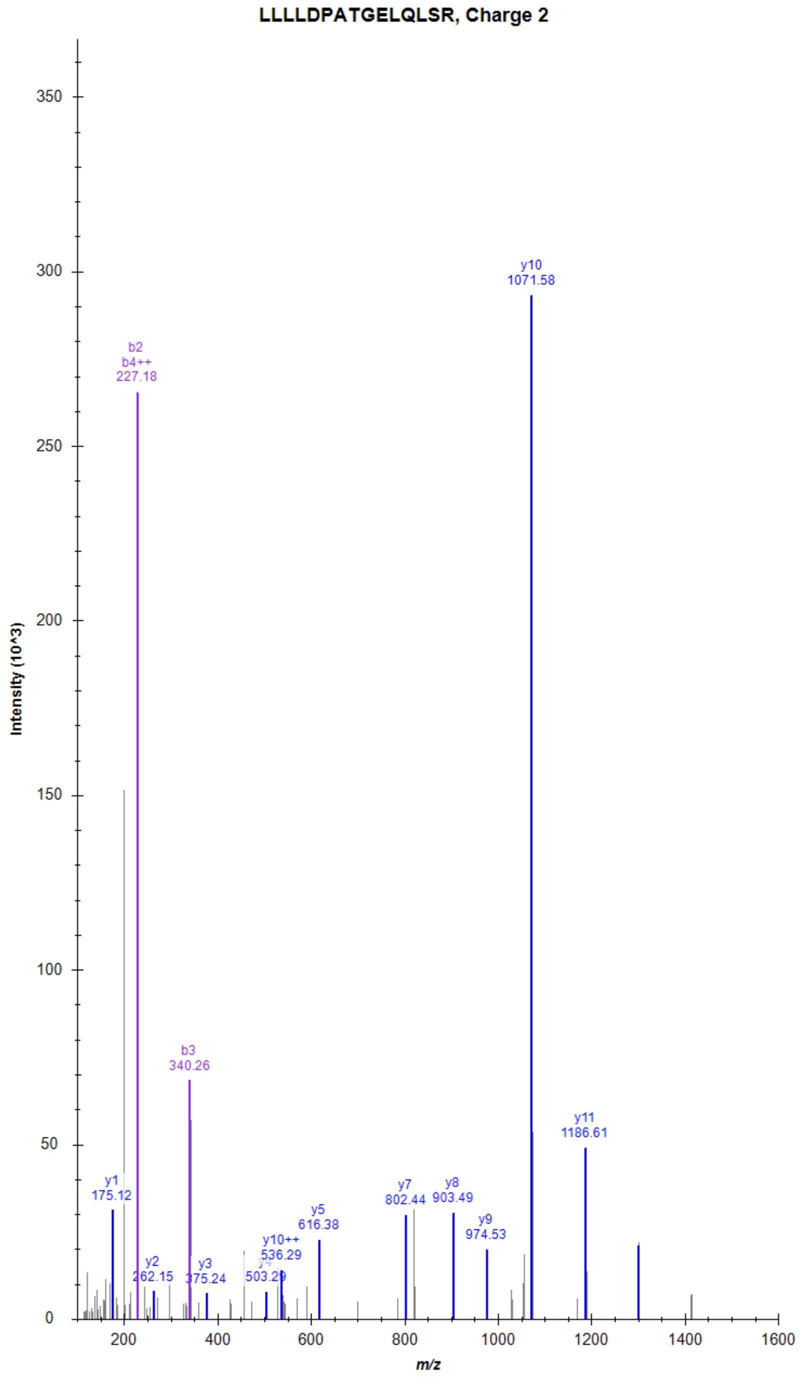

### Adhesion G-protein coupled receptor G1 (ADGRG1)

LQPTAGLQDLHIHSR, Charge 3, m/z = 562.6406

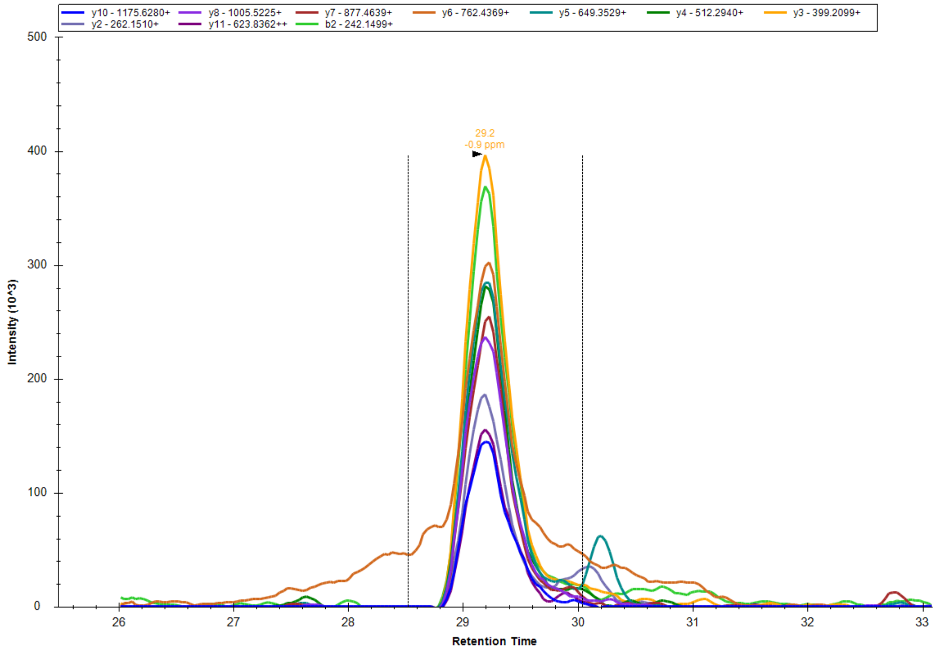


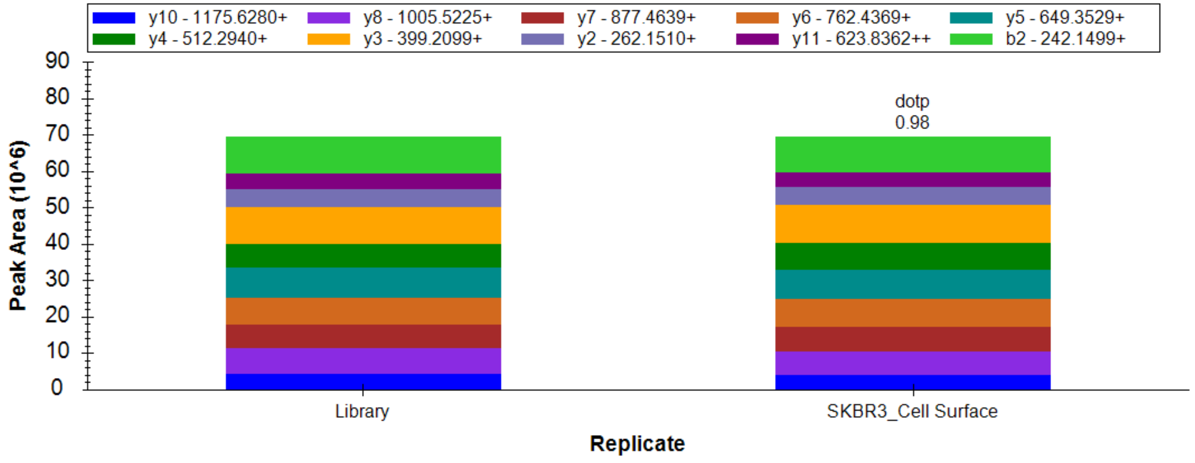


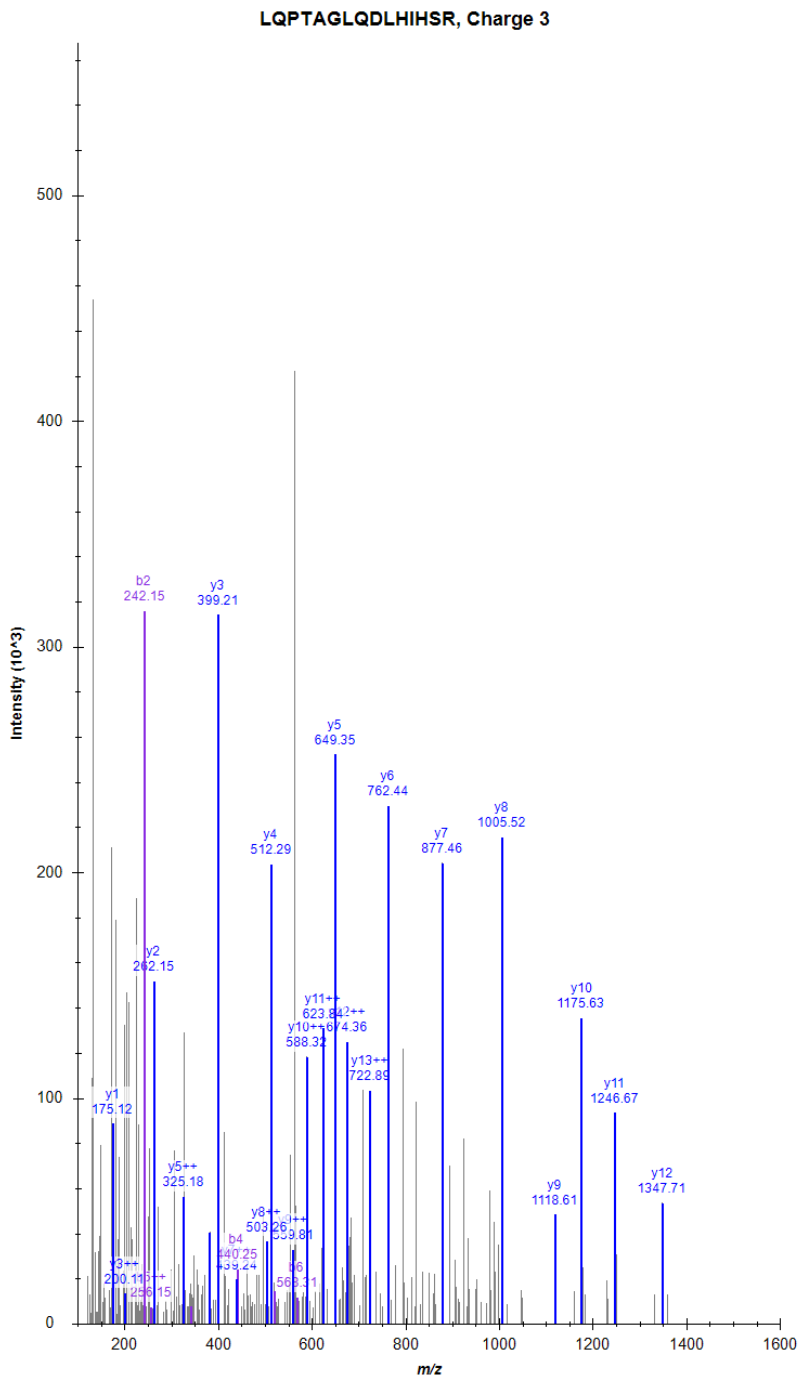

### Adhesion G-protein coupled receptor G1 (ADGRG1)

QEEEQSEIMEYSVLLPR, Charge 2, m/z = 1040.4957

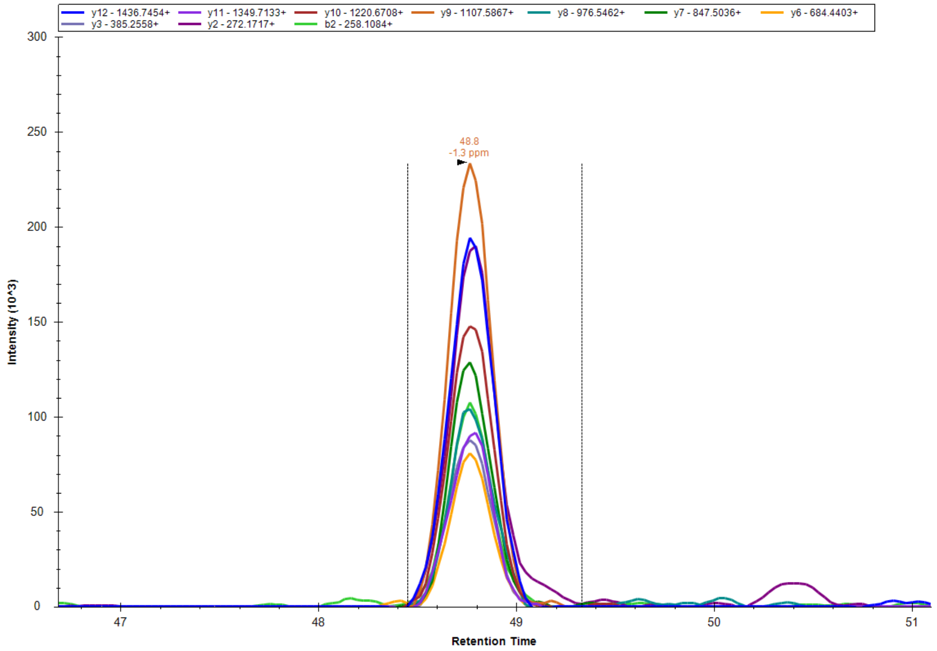


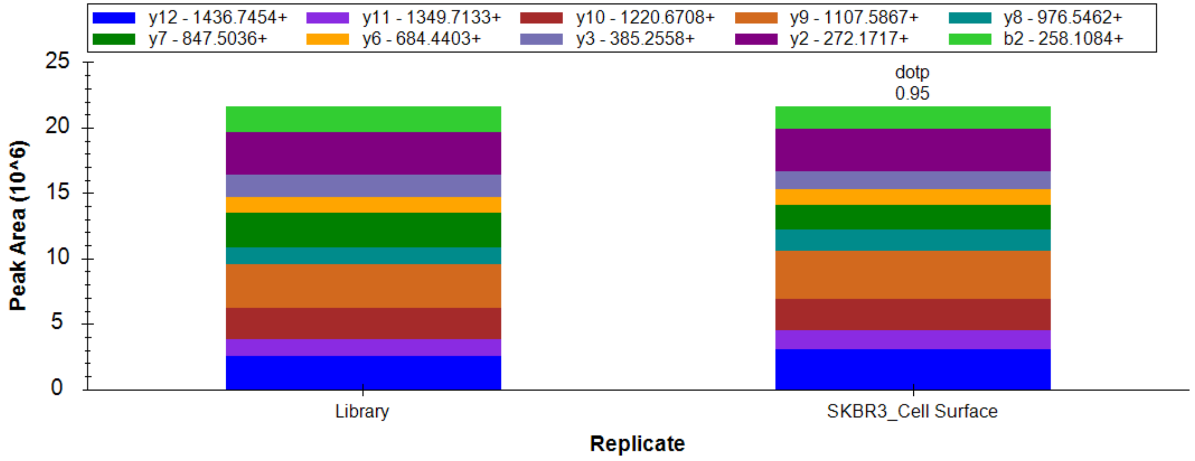


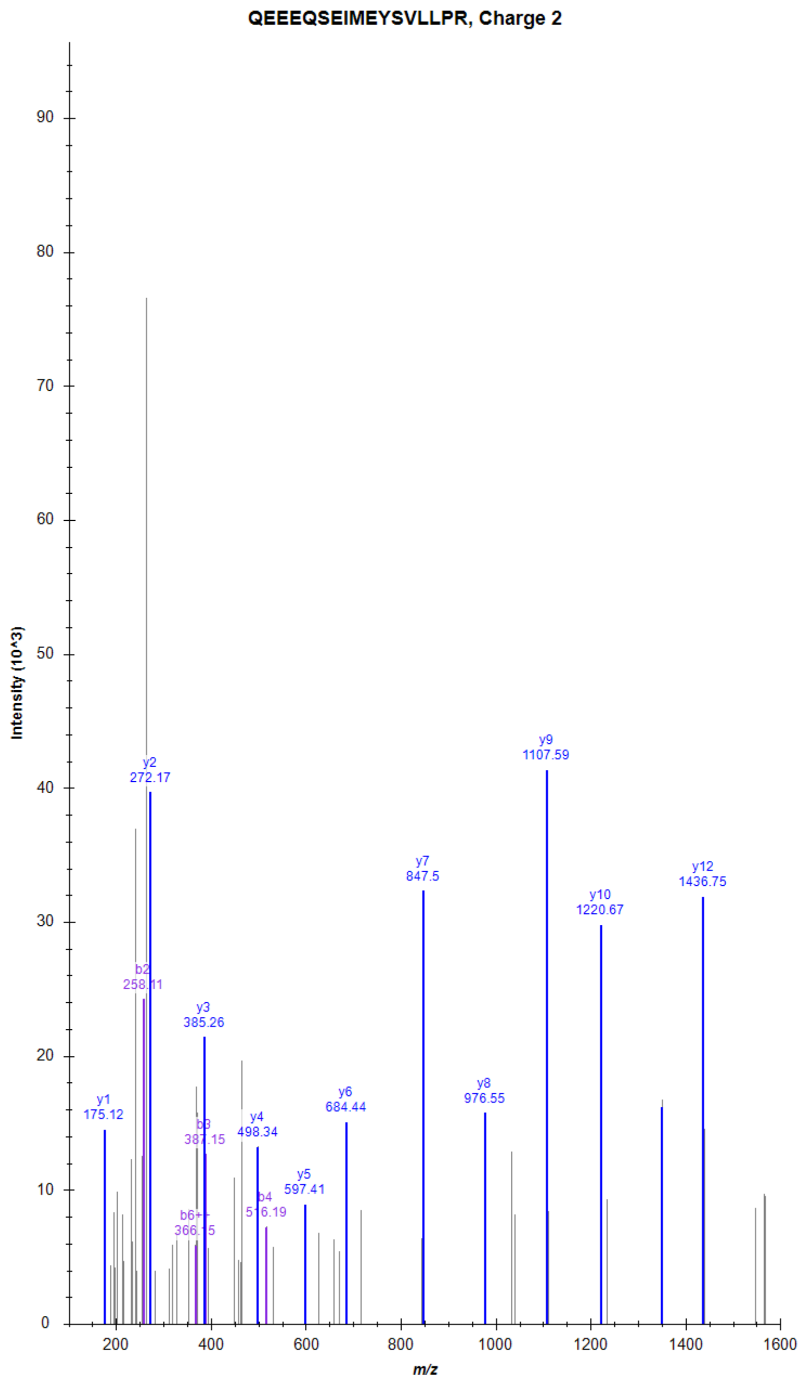

### Adhesion G-protein coupled receptor E2 (ADGRE2)

SGDPGPSVVGLVSIPGMGK, Charge 2, m/z = 877.4613

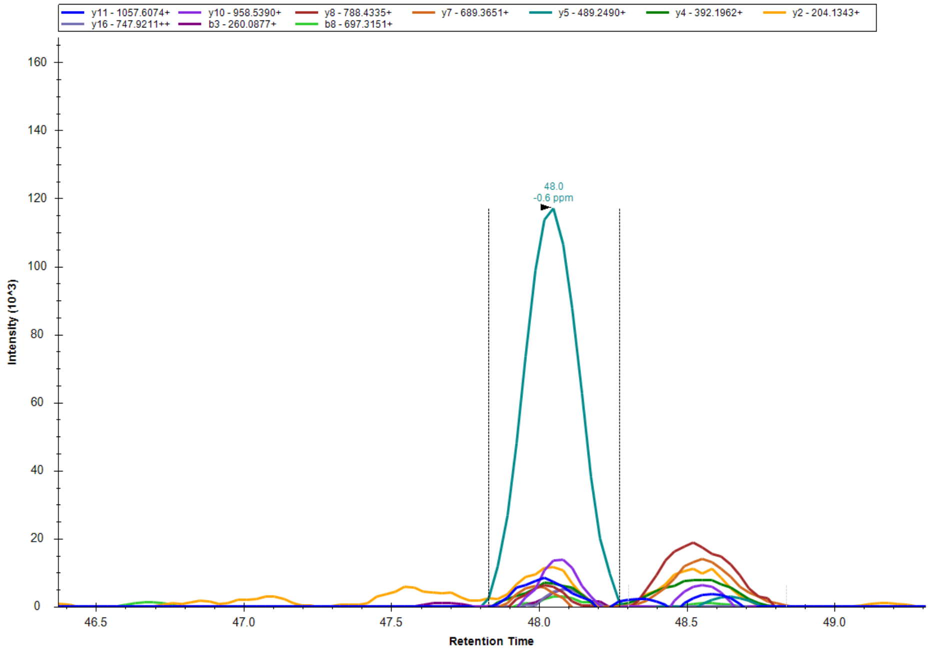


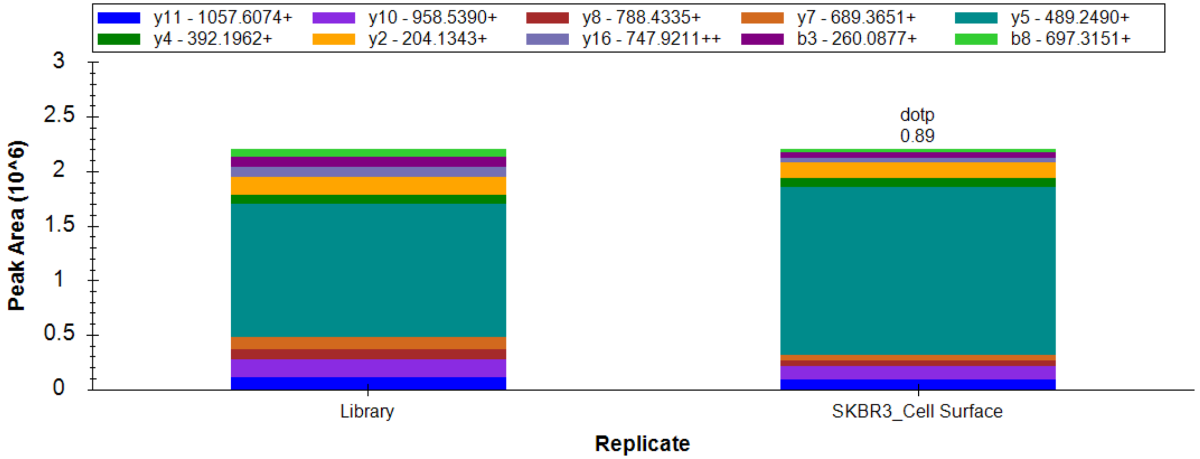


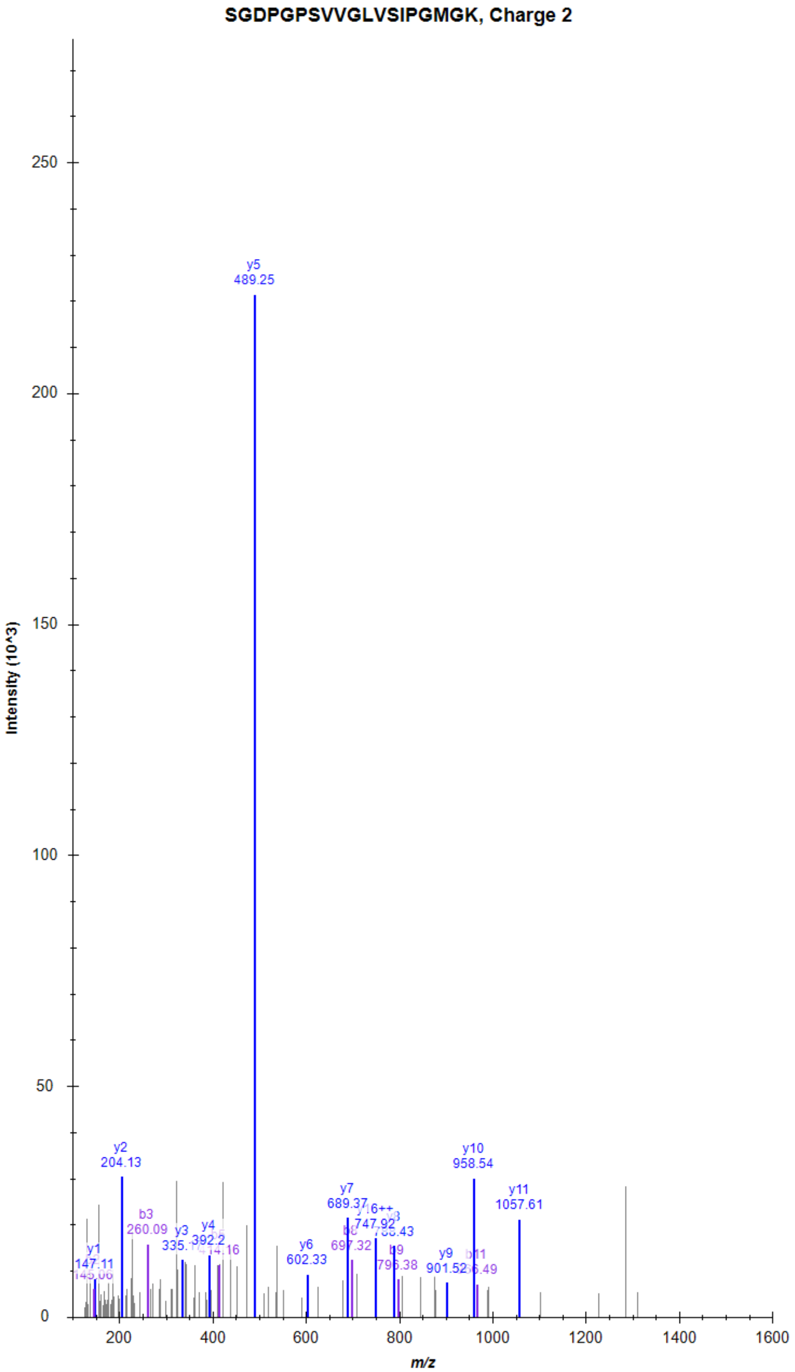

### Adhesion G-protein coupled receptor E2 (ADGRE2)

LLAEAPLVLEPEK, Charge 2, m/z = 711.4155

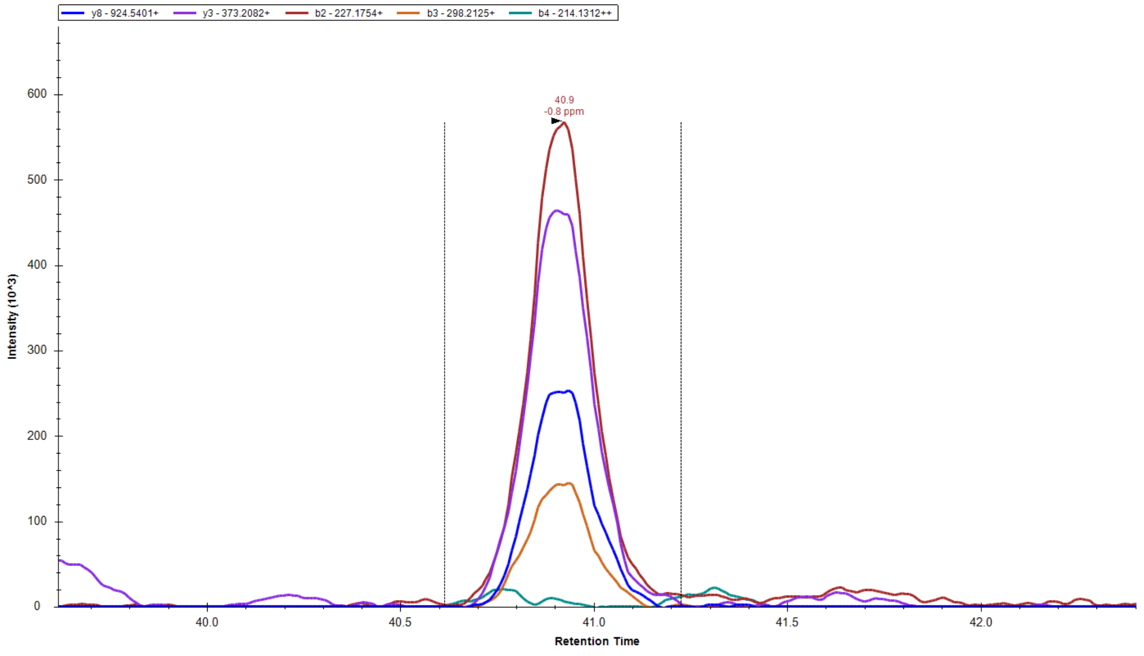


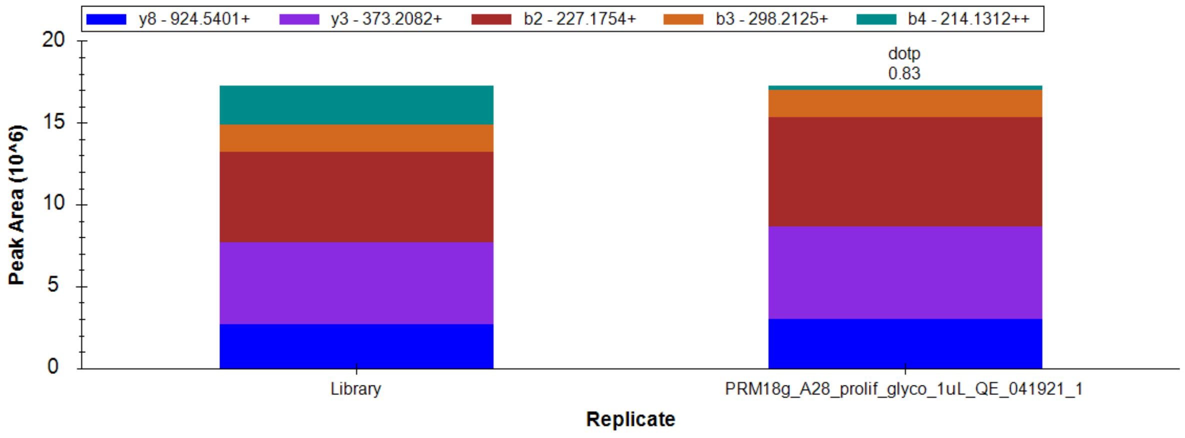


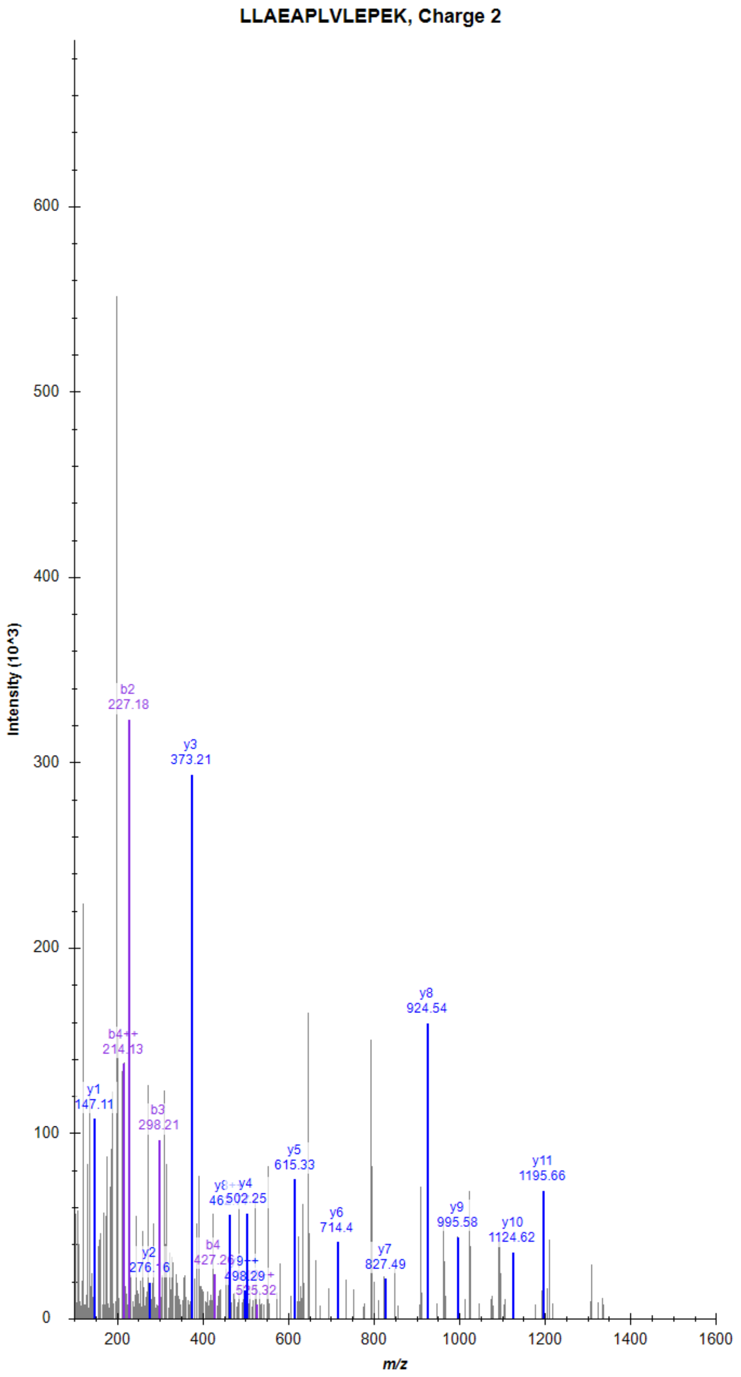

### Muscarinic acetylcholine receptor M1 (CHRM1)

MPMVDPEAQAPTK, Charge 2, m/z = 707.8388

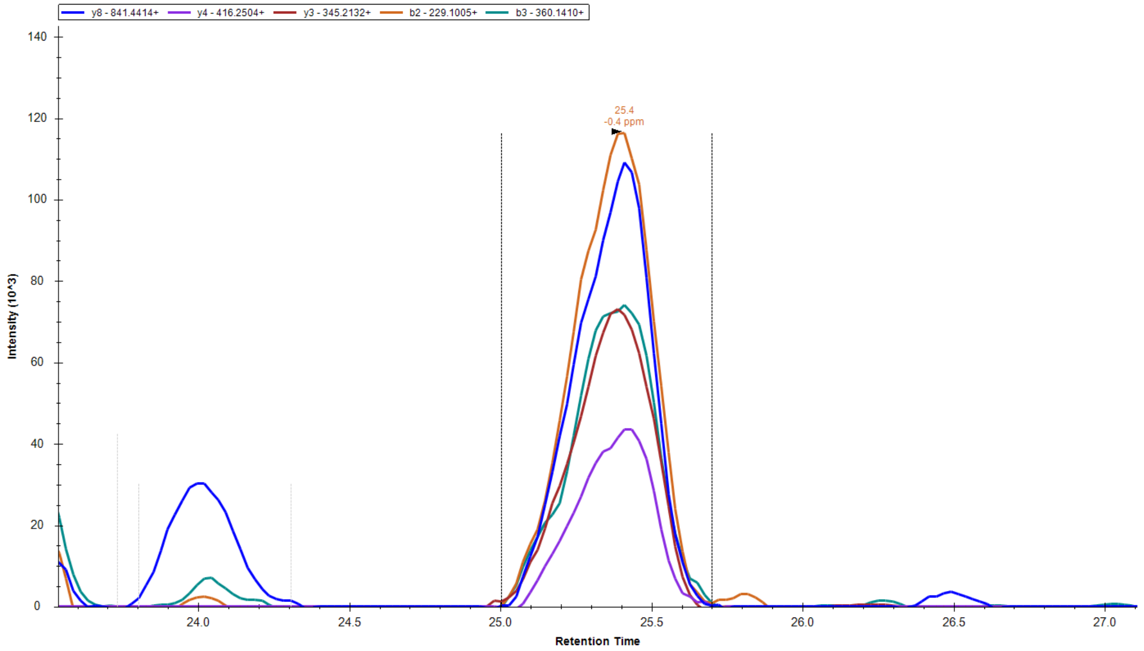


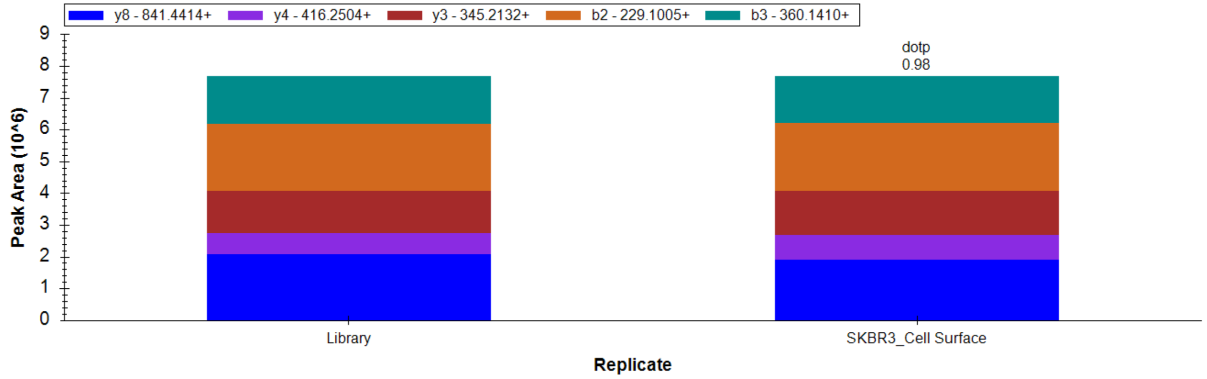


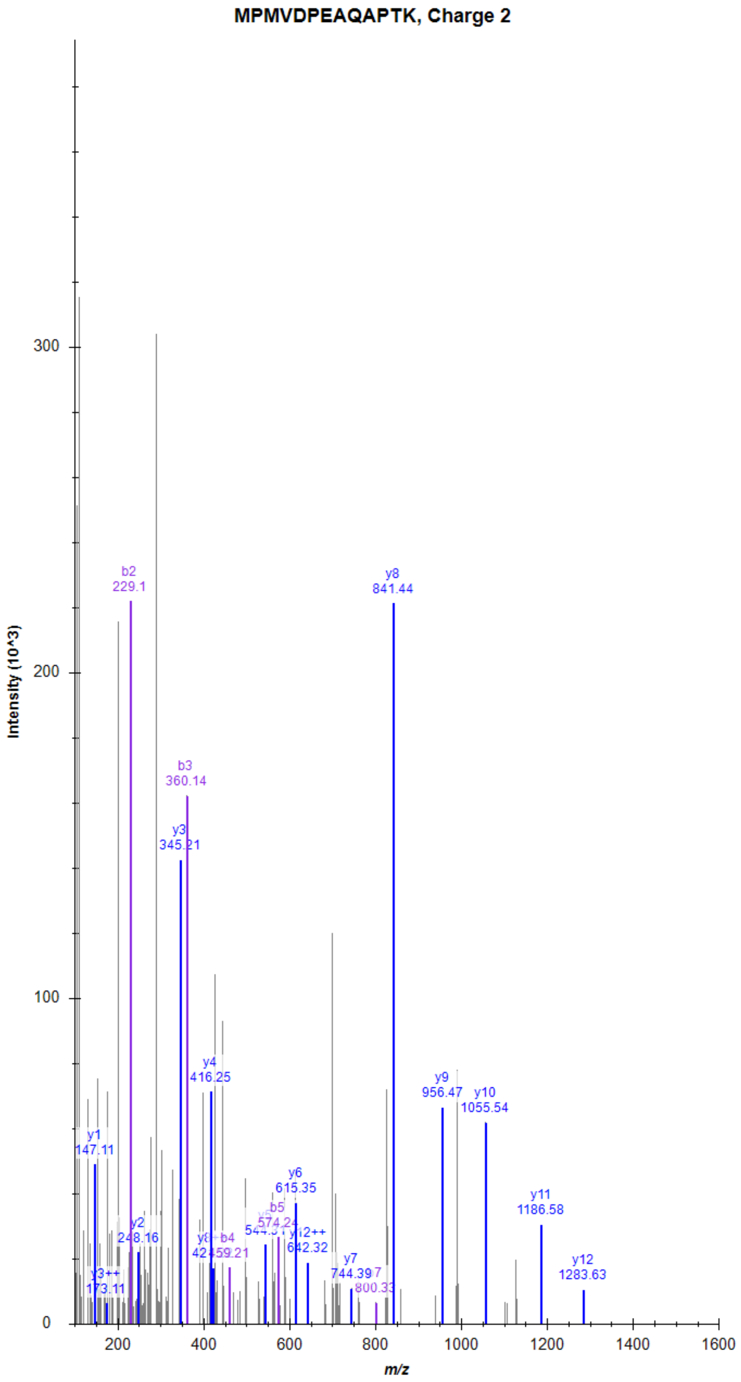

### Muscarinic acetylcholine receptor M1 (CHRM1)

ELAALQGSETPGK, Charge 2, m/z = 650.8407

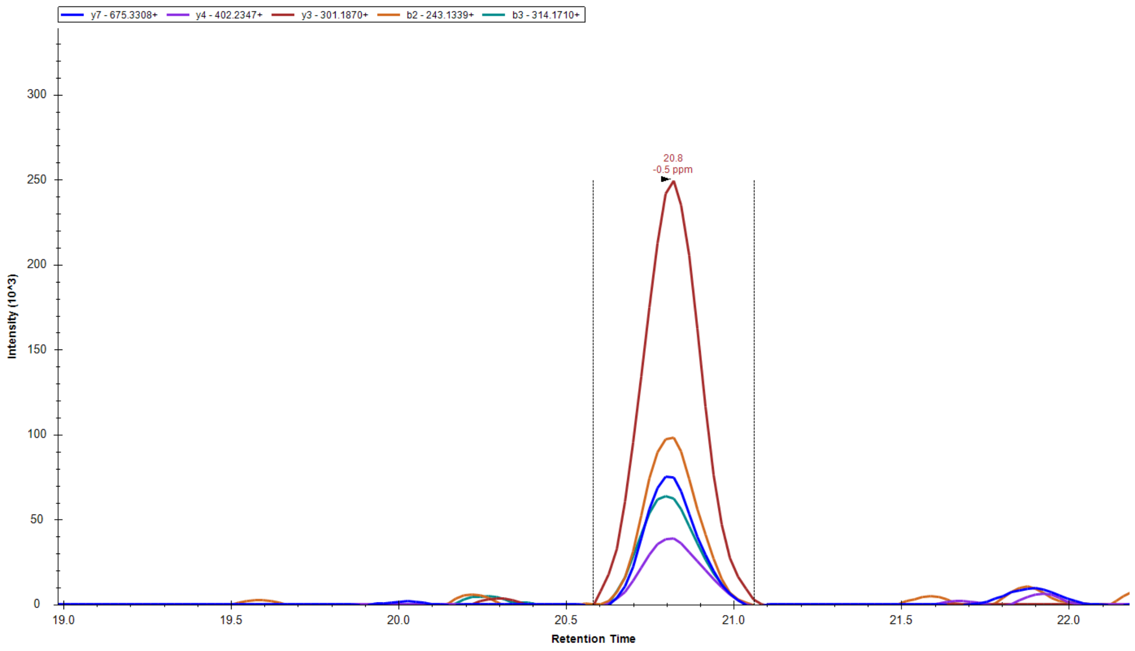


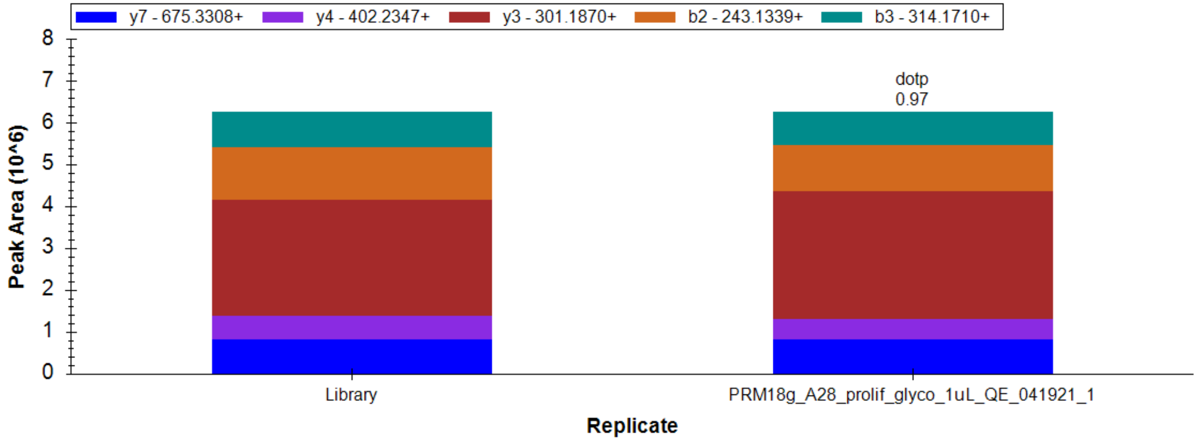


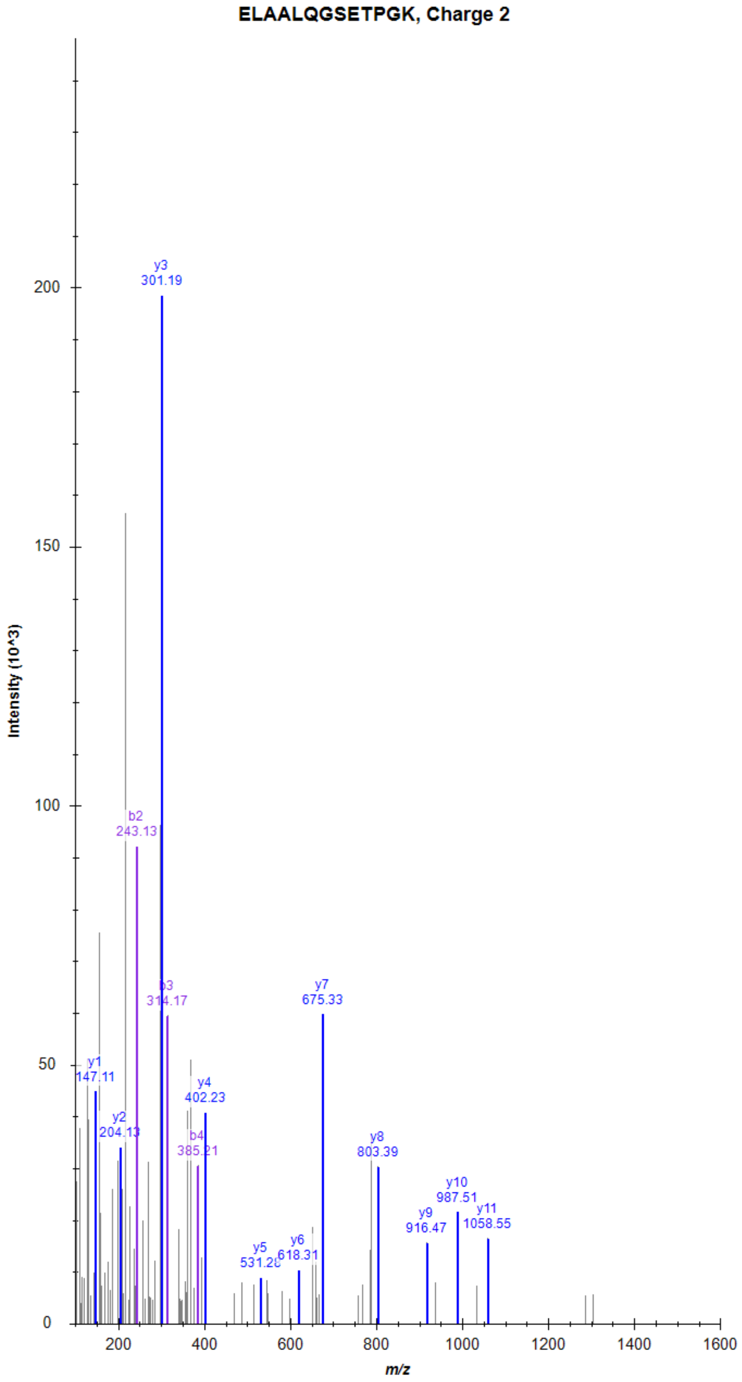

### P2Y purinoceptor 2 (P2RY2)

LLKPAYGTSGGLPR, Charge 3, m/z = 477.2769

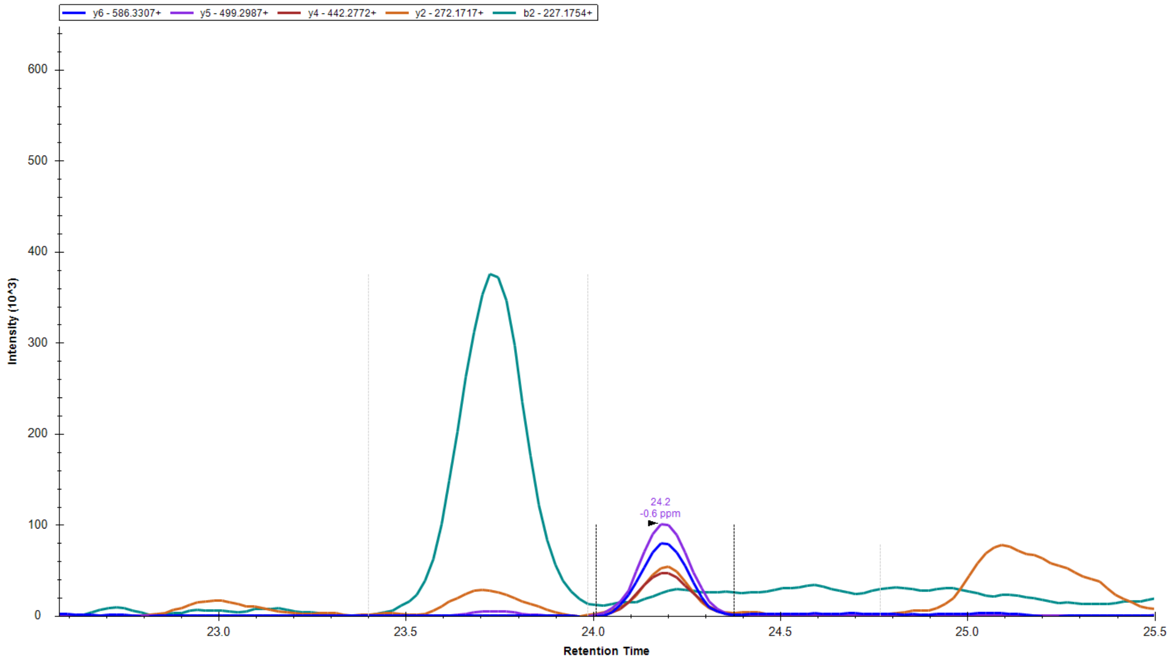


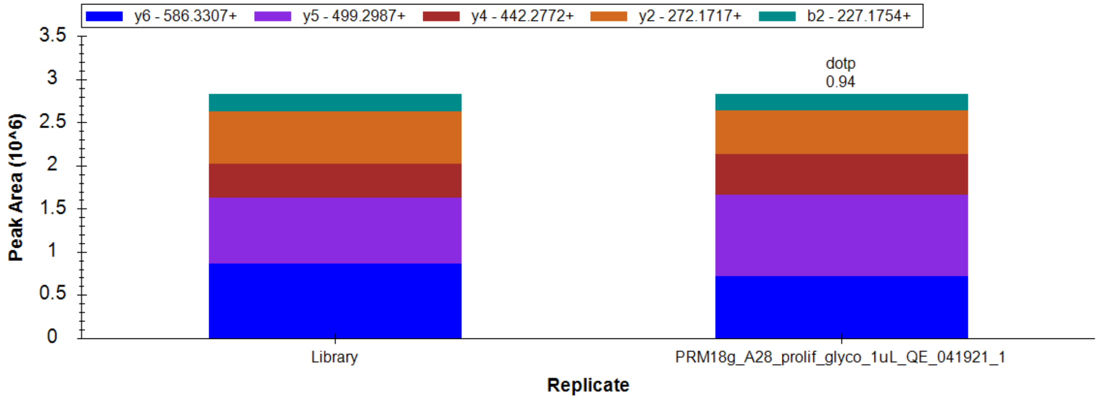


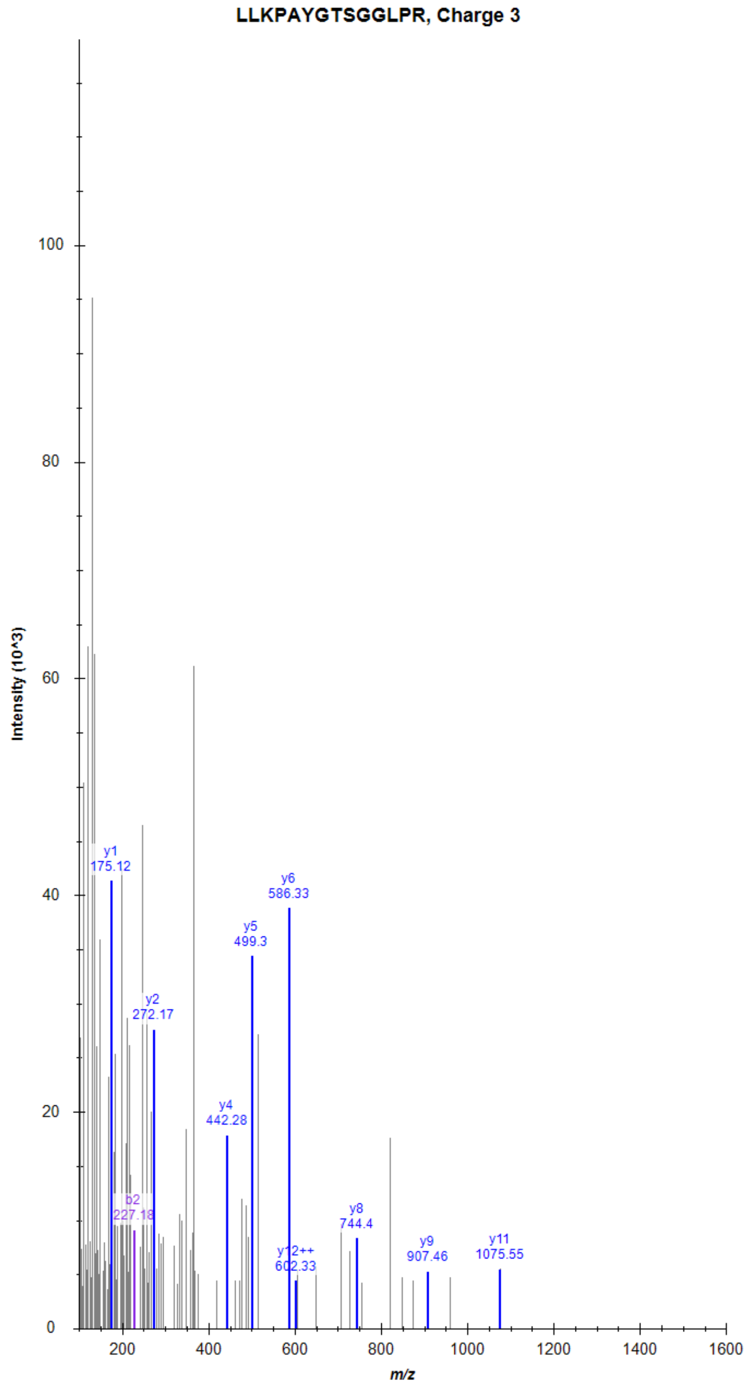

### P2Y purinoceptor 2 (P2RY2)

IEDVLGSSEDSR, Charge 2, m/z = 653.8104

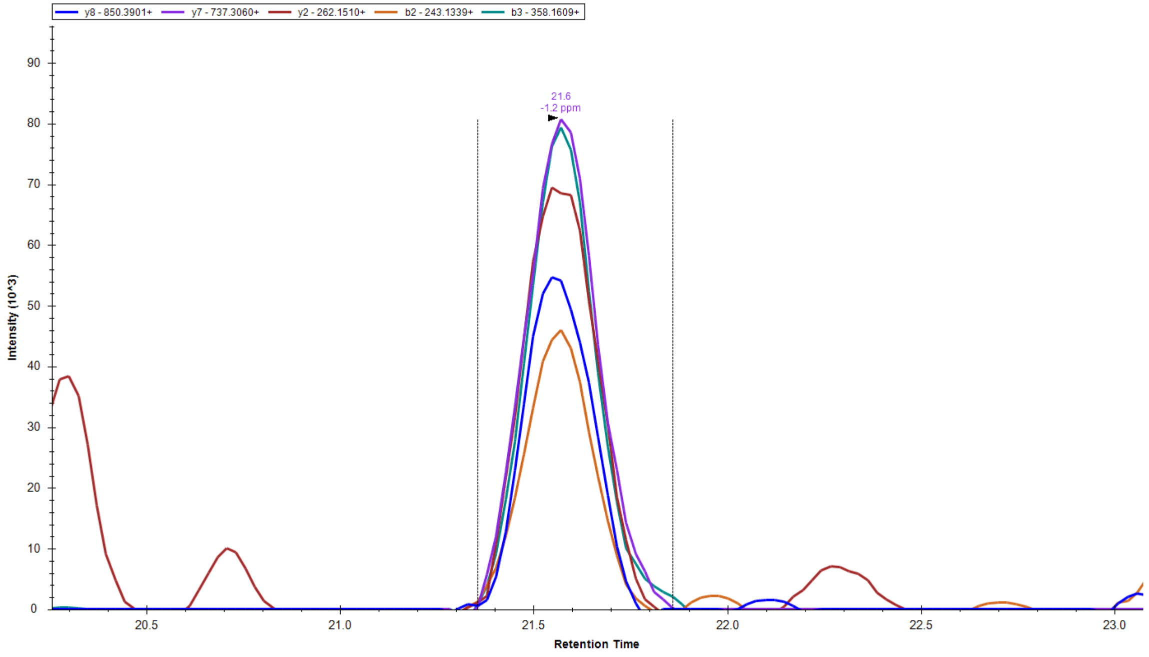


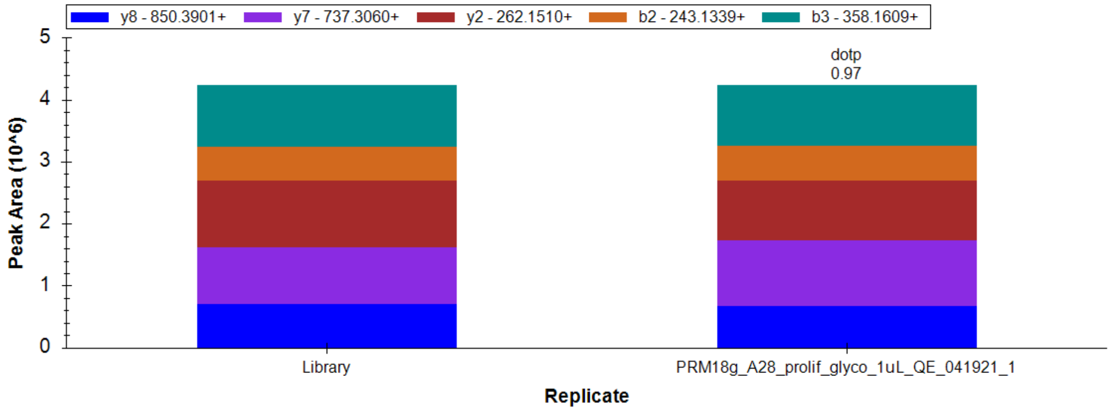


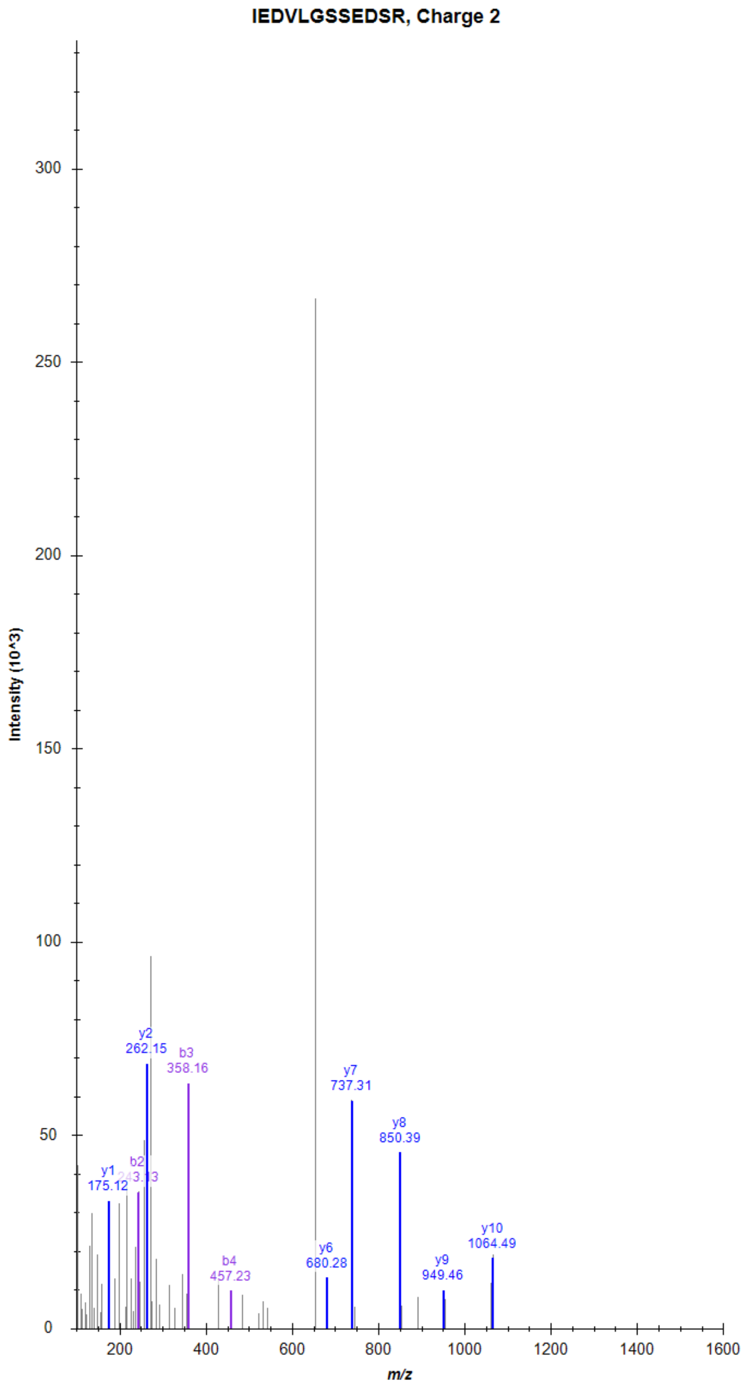

### B2 bradykinin receptor (BDKRB2)

SEPIQMENSMGTLR, Charge 2, m/z = 796.8734

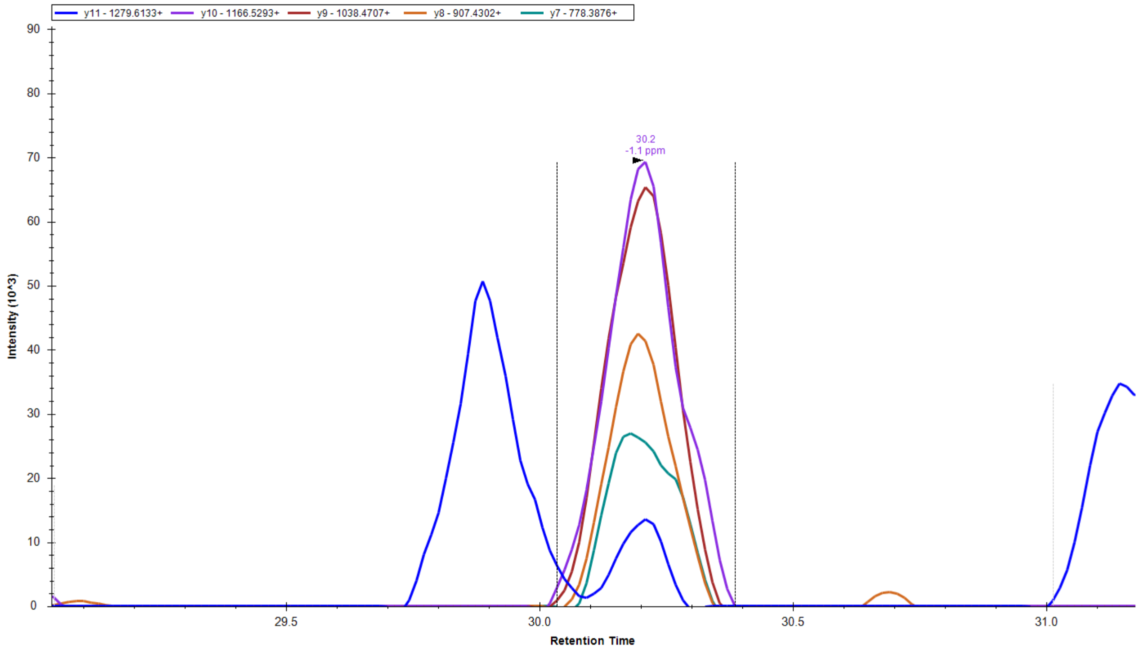


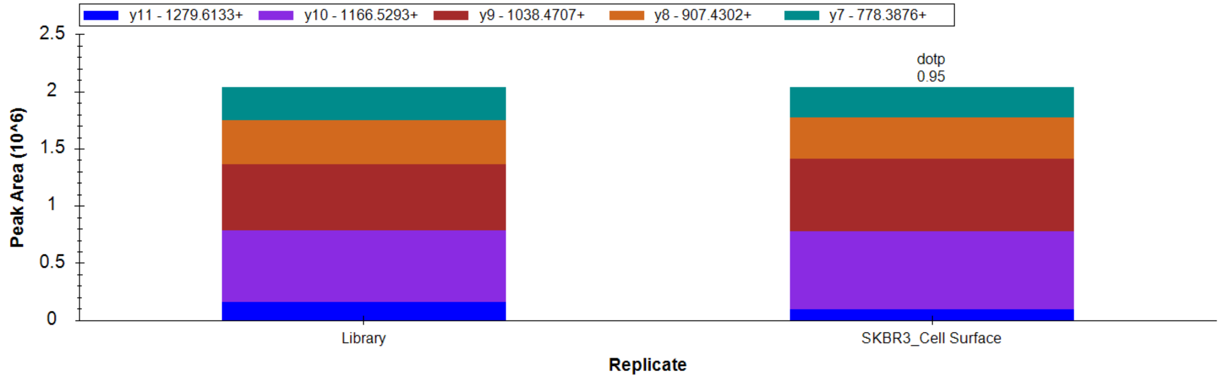


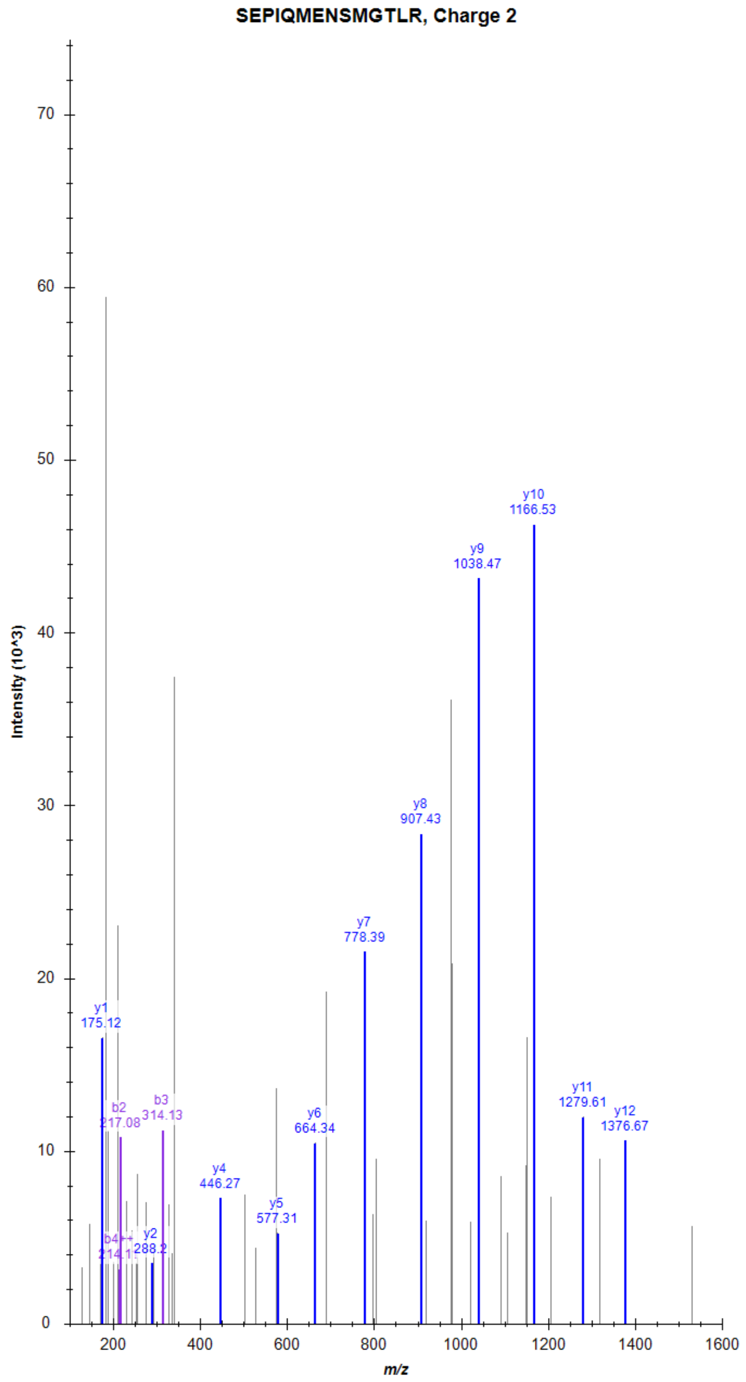

### B2 bradykinin receptor (BDKRB2)

LQDWAGSR, Charge 2, m/z = 466.7322

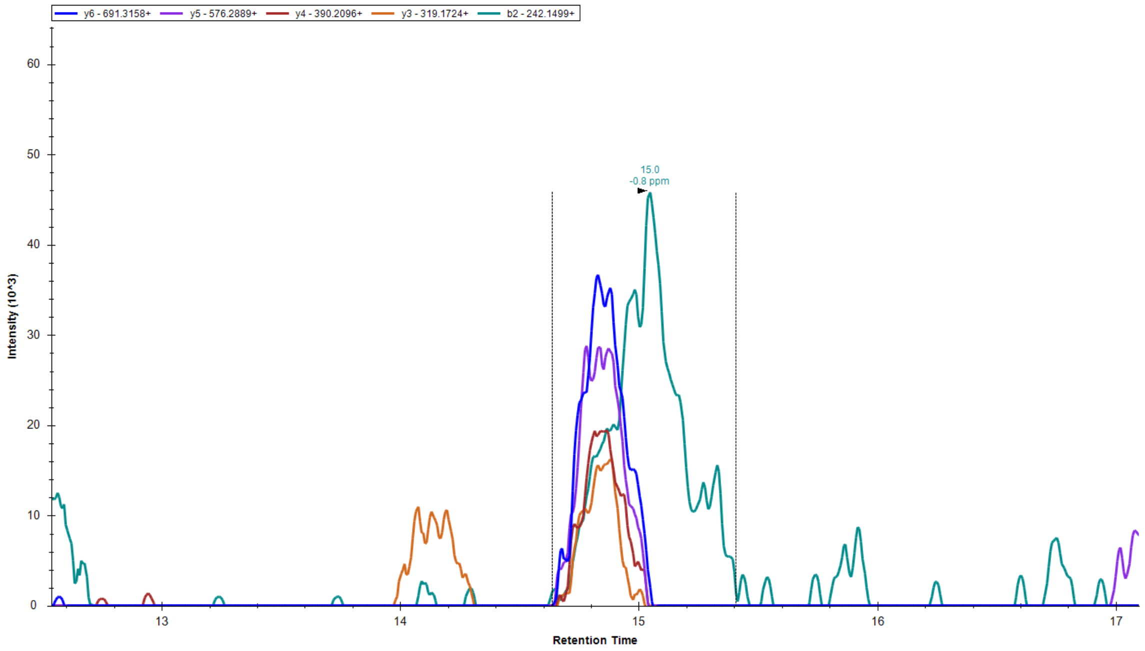


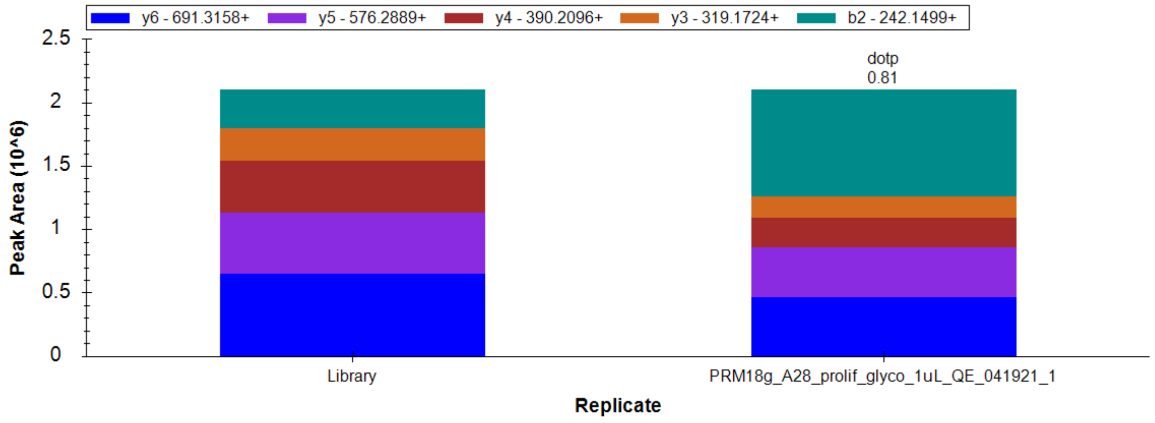


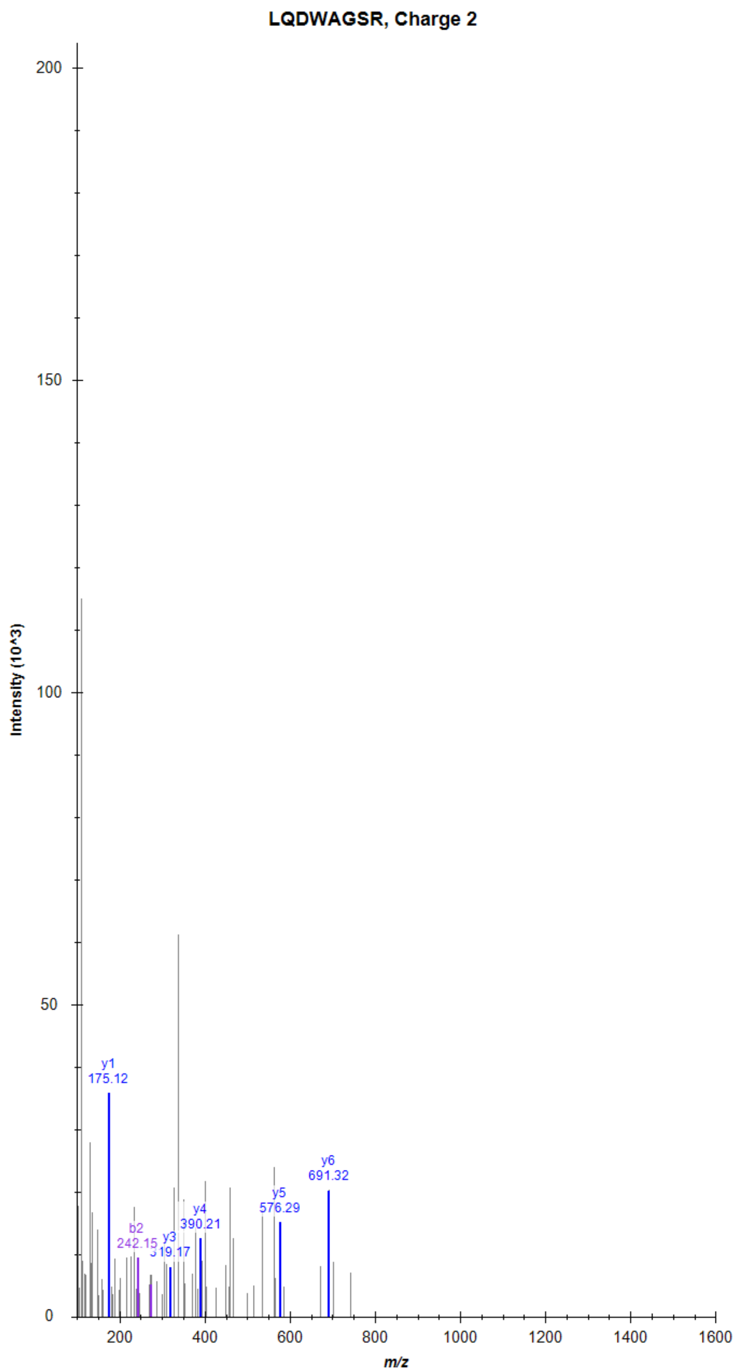

### Frizzled-1 (FZD1)

VYGLMYFGPEELR, Charge 2, m/z = 787.3892

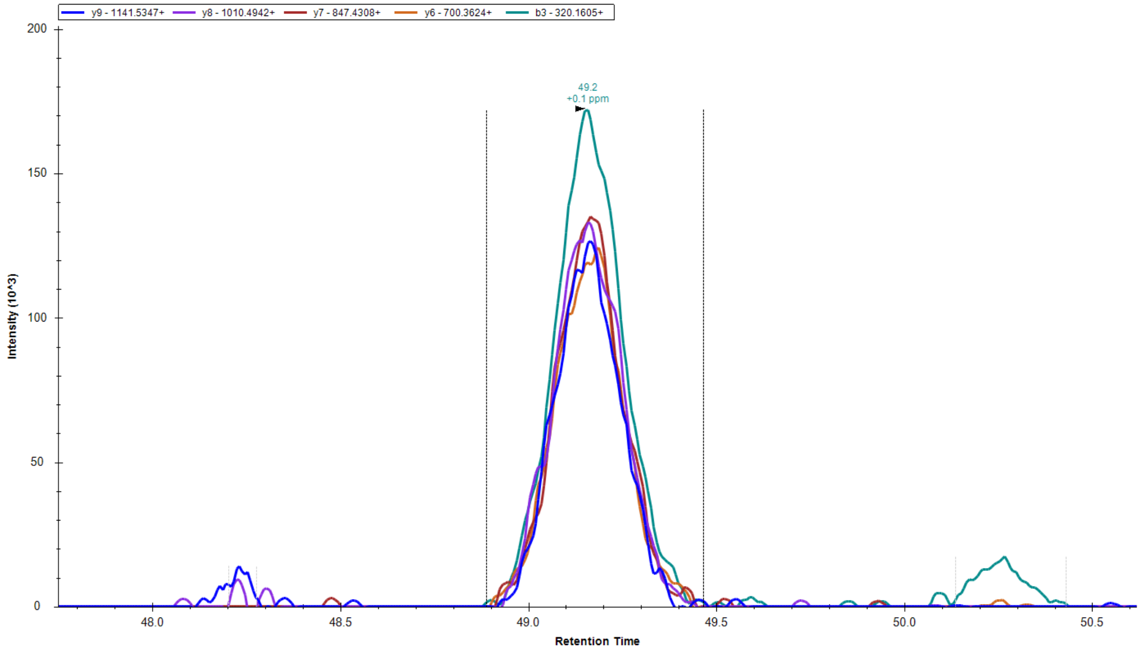


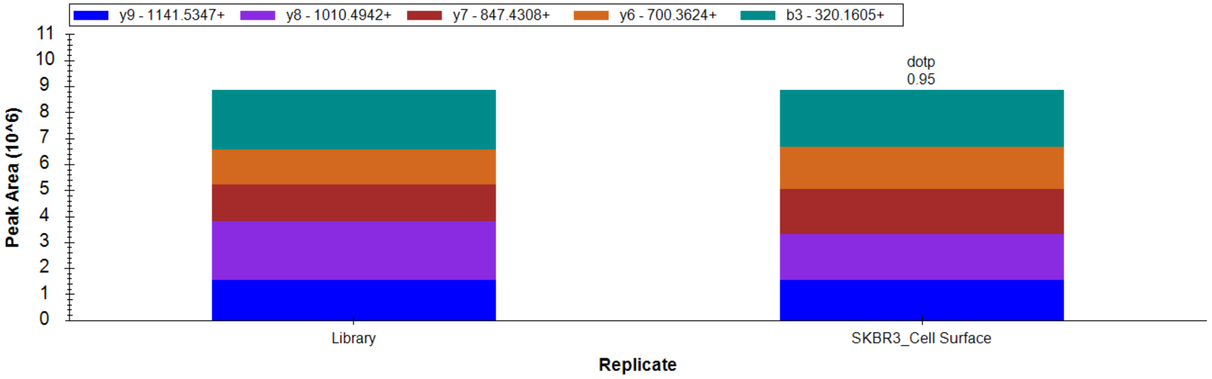


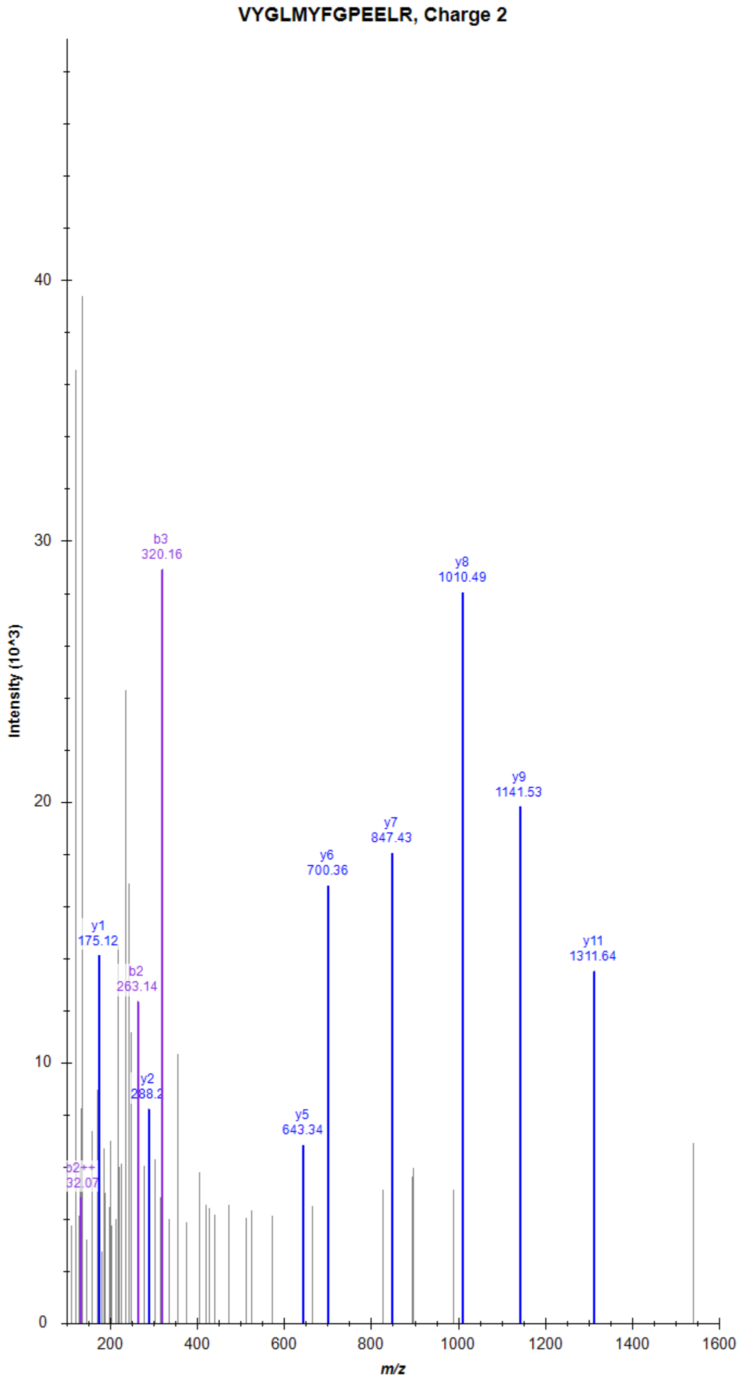

### Cadherin EGF LAG seven-pass G-type receptor 2 (CELSR2)

SLDLTGPLLLGGVPDLPESFPVR, Charge 2, m/z = 1196.663

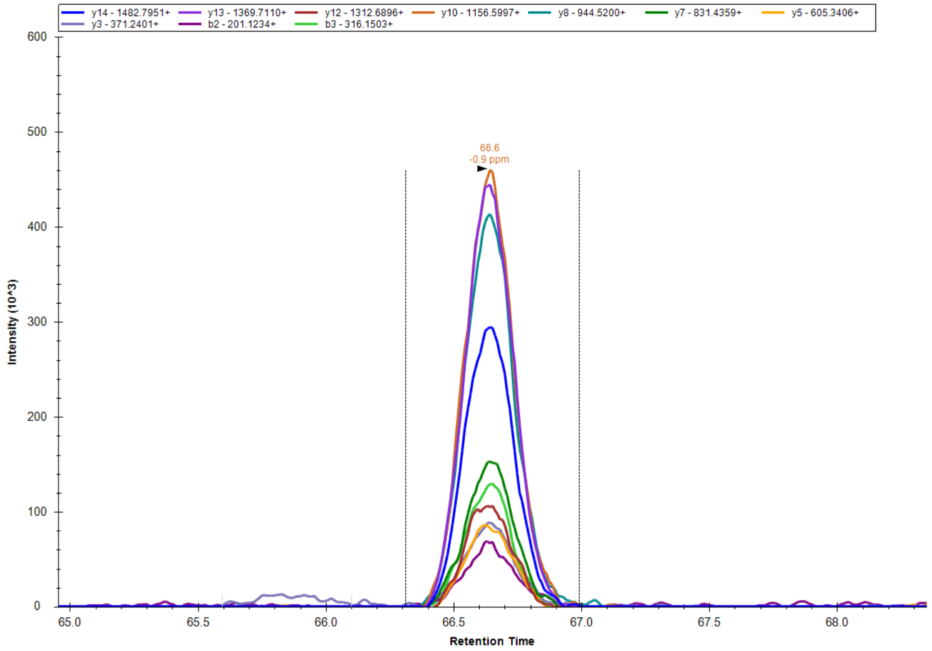


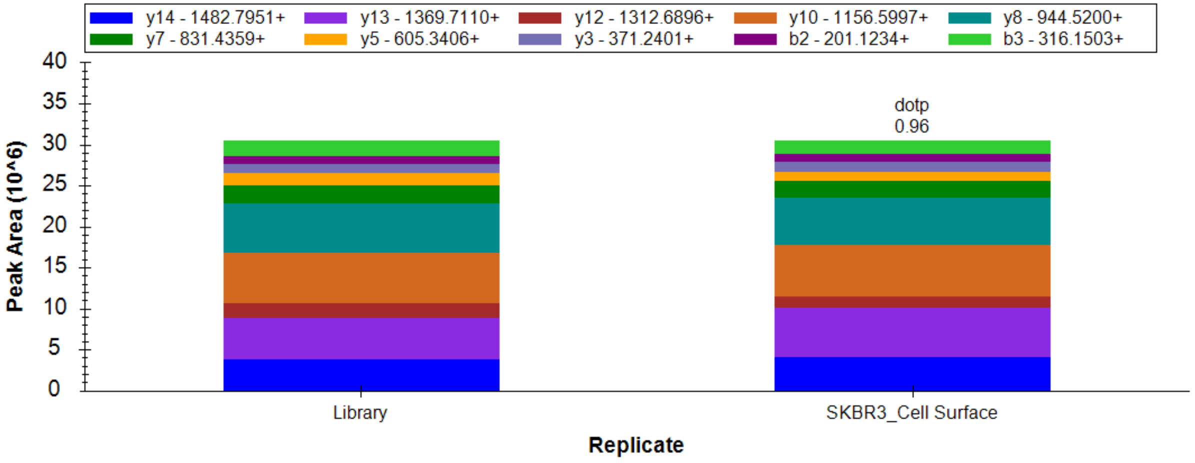


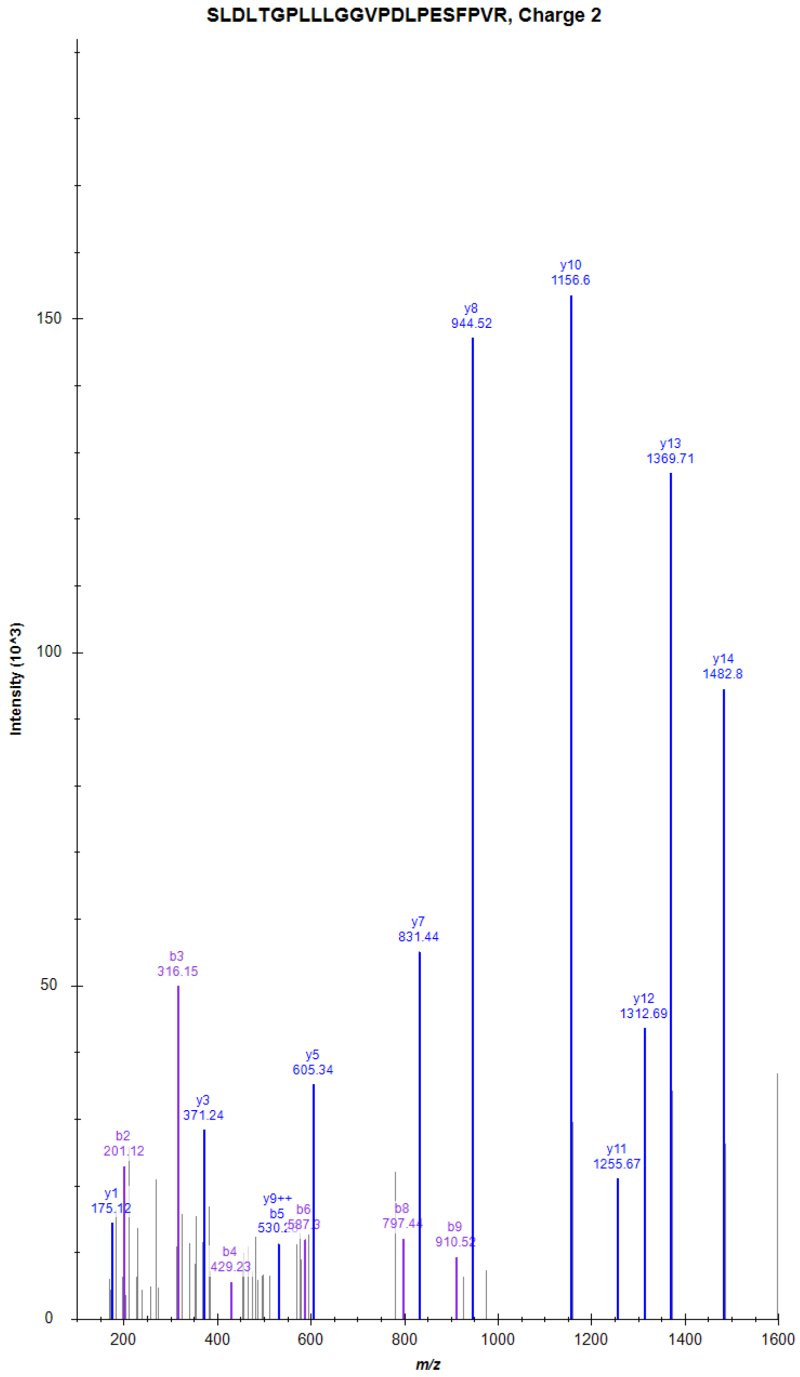

### G-protein coupled receptor 39 (GPR39)

IFLSTFQSEAEPQSK, Charge 2, m/z = 856.4298

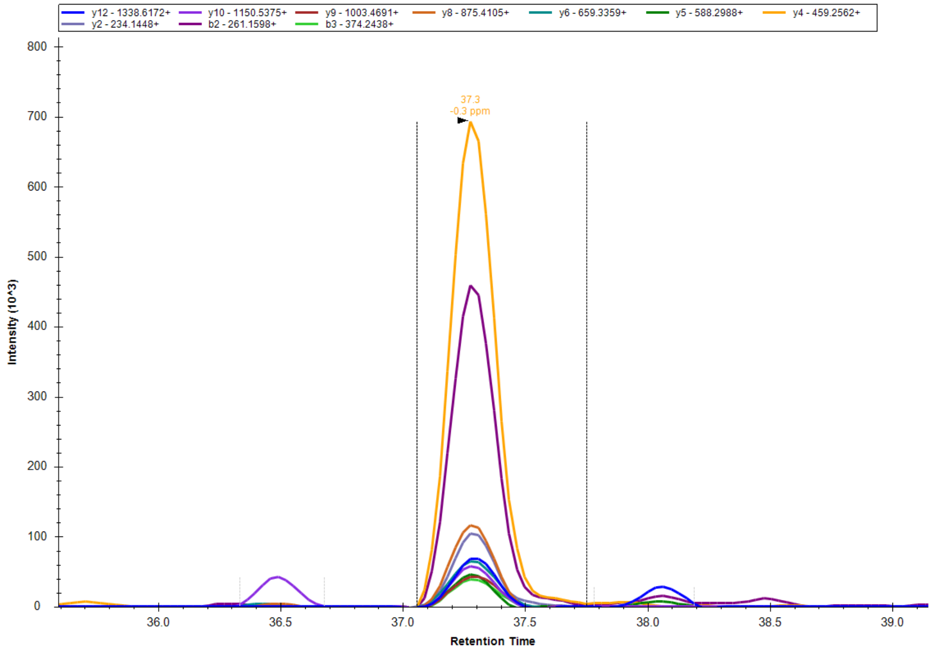


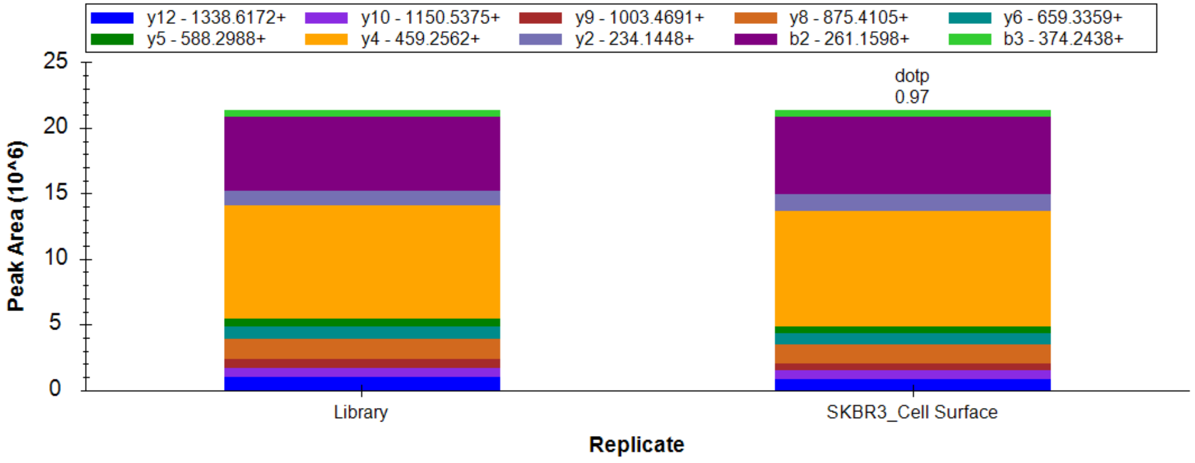


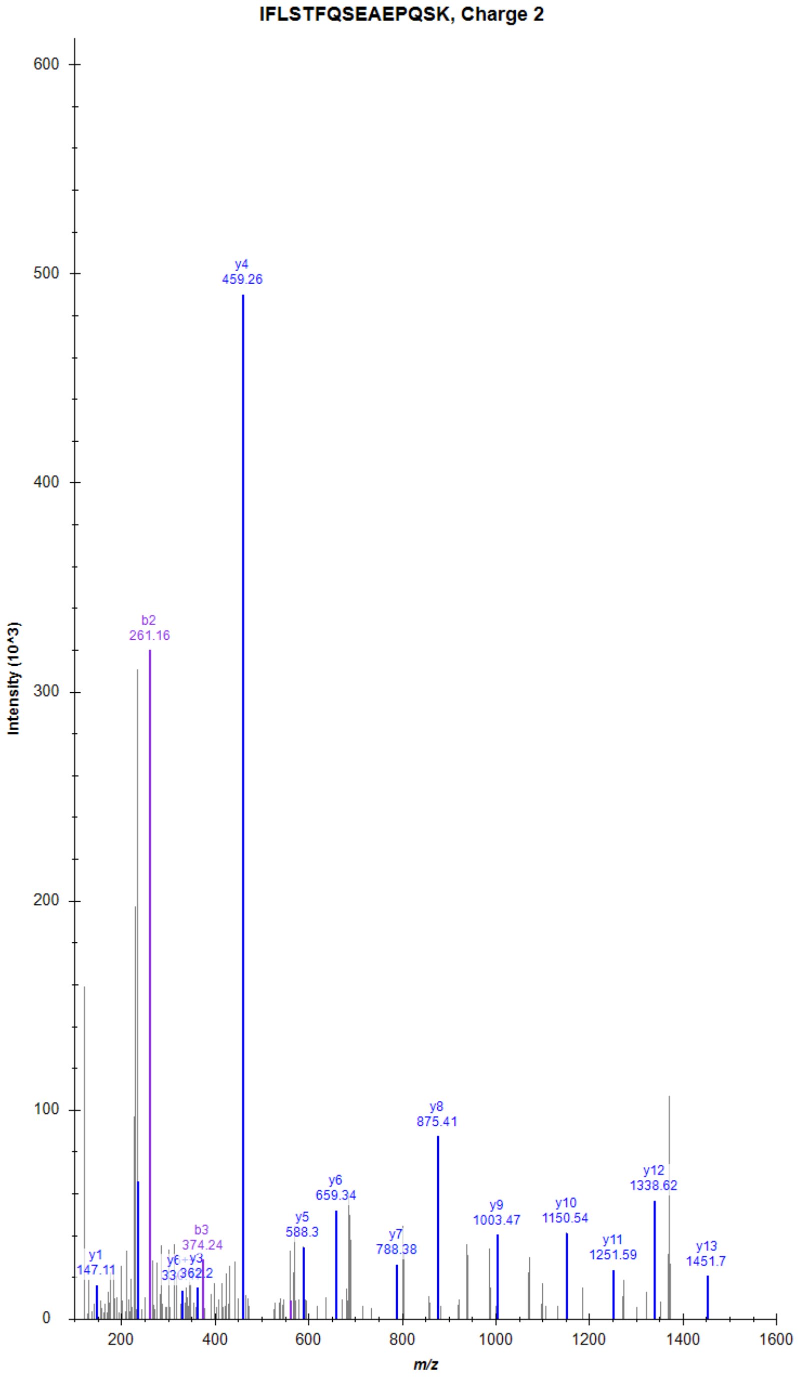
